# Widespread variation in molecular interactions and regulatory properties among transcription factor isoforms

**DOI:** 10.1101/2024.03.12.584681

**Authors:** Luke Lambourne, Kaia Mattioli, Clarissa Santoso, Gloria Sheynkman, Sachi Inukai, Babita Kaundal, Anna Berenson, Kerstin Spirohn-Fitzgerald, Anukana Bhattacharjee, Elisabeth Rothman, Shaleen Shrestha, Florent Laval, Zhipeng Yang, Deepa Bisht, Jared A. Sewell, Guangyuan Li, Anisa Prasad, Sabrina Phanor, Ryan Lane, Devlin M. Campbell, Toby Hunt, Dawit Balcha, Marinella Gebbia, Jean-Claude Twizere, Tong Hao, Adam Frankish, Josh A. Riback, Nathan Salomonis, Michael A. Calderwood, David E. Hill, Nidhi Sahni, Marc Vidal, Martha L. Bulyk, Juan I. Fuxman Bass

**Affiliations:** Center for Cancer Systems Biology (CCSB), Dana-Farber Cancer Institute, Boston, MA, USA; Department of Genetics, Blavatnik Institute, Harvard Medical School, Boston, MA, USA; Department of Cancer Biology, Dana-Farber Cancer Institute, Boston, MA, USA; Division of Genetics, Department of Medicine, Brigham and Women’s Hospital and Harvard Medical School, Boston, MA, USA; Department of Biology, Boston University, Boston, MA, USA; Bioinformatics Program, Boston University, Boston, MA, USA; Department of Epigenetics and Molecular Carcinogenesis, The University of Texas MD Anderson Cancer Center, Houston, TX, USA; Molecular Biology, Cell Biology & Biochemistry Program, Boston University, Boston, MA, USA; Department of Pediatrics, University of Cincinnati College of Medicine, Cincinnati, OH, USA; Division of Biomedical Informatics, Cincinnati Children’s Hospital Medical Center, Cincinnati, OH, USA; TERRA Teaching and Research Centre, University of Liège, Gembloux, Belgium; Laboratory of Viral Interactomes, GIGA Institute, University of Liège, Liège, Belgium; Harvard College, Cambridge MA, USA; European Molecular Biology Laboratory, European Bioinformatics Institute, Wellcome Genome Campus, Hinxton, Cambridge, UK; The Donnelly Centre, University of Toronto, Toronto, Ontario, CanadaA; Department of Molecular Genetics, University of Toronto, Toronto, Ontario, Canada; Lunenfeld-Tanenbaum Research Institute (LTRI), Sinai Health System, Toronto, Ontario, Canada; Department of Molecular and Cellular Biology, Baylor College of Medicine, Houston, TX, USA; Department of Pathology, Brigham and Women’s Hospital and Harvard Medical School, Boston, MA, USA

## Abstract

Most human Transcription factors (TFs) genes encode multiple protein isoforms differing in DNA binding domains, effector domains, or other protein regions. The global extent to which this results in functional differences between isoforms remains unknown. Here, we systematically compared 693 isoforms of 246 TF genes, assessing DNA binding, protein binding, transcriptional activation, subcellular localization, and condensate formation. Relative to reference isoforms, two-thirds of alternative TF isoforms exhibit differences in one or more molecular activities, which often could not be predicted from sequence. We observed two primary categories of alternative TF isoforms: “rewirers” and “negative regulators”, both of which were associated with differentiation and cancer. Our results support a model wherein the relative expression levels of, and interactions involving, TF isoforms add an understudied layer of complexity to gene regulatory networks, demonstrating the importance of isoform-aware characterization of TF functions and providing a rich resource for further studies.

## Introduction

Gene regulatory programs are a major driver of cellular phenotypes in development and disease. Gene regulation is mediated through activities of transcription factors (TFs) which interact in a sequence-specific manner with their target DNA elements to activate or repress gene expression.^1^ The last four decades have seen an explosion in studies and throughput to determine the DNA binding specificities,^2–5^ transcriptional activities,^6–8^ and protein-protein interactions^9–11^ of TFs in physiological and pathological conditions. Historically, these efforts have mostly focused on generating functional profiles for the wildtype, canonical reference isoforms of TFs or for specific domains. However, such studies rarely consider that TF genes, like most other genes, encode multiple “proteoforms” due to: i) alternative transcript isoforms resulting from alternative promoter, splice site/junction, and/or terminal exon usage; ii) naturally occurring coding variation across individuals; and iii) post-translational modifications (**Figure 1A**).^12–14^

**Figure 1:**
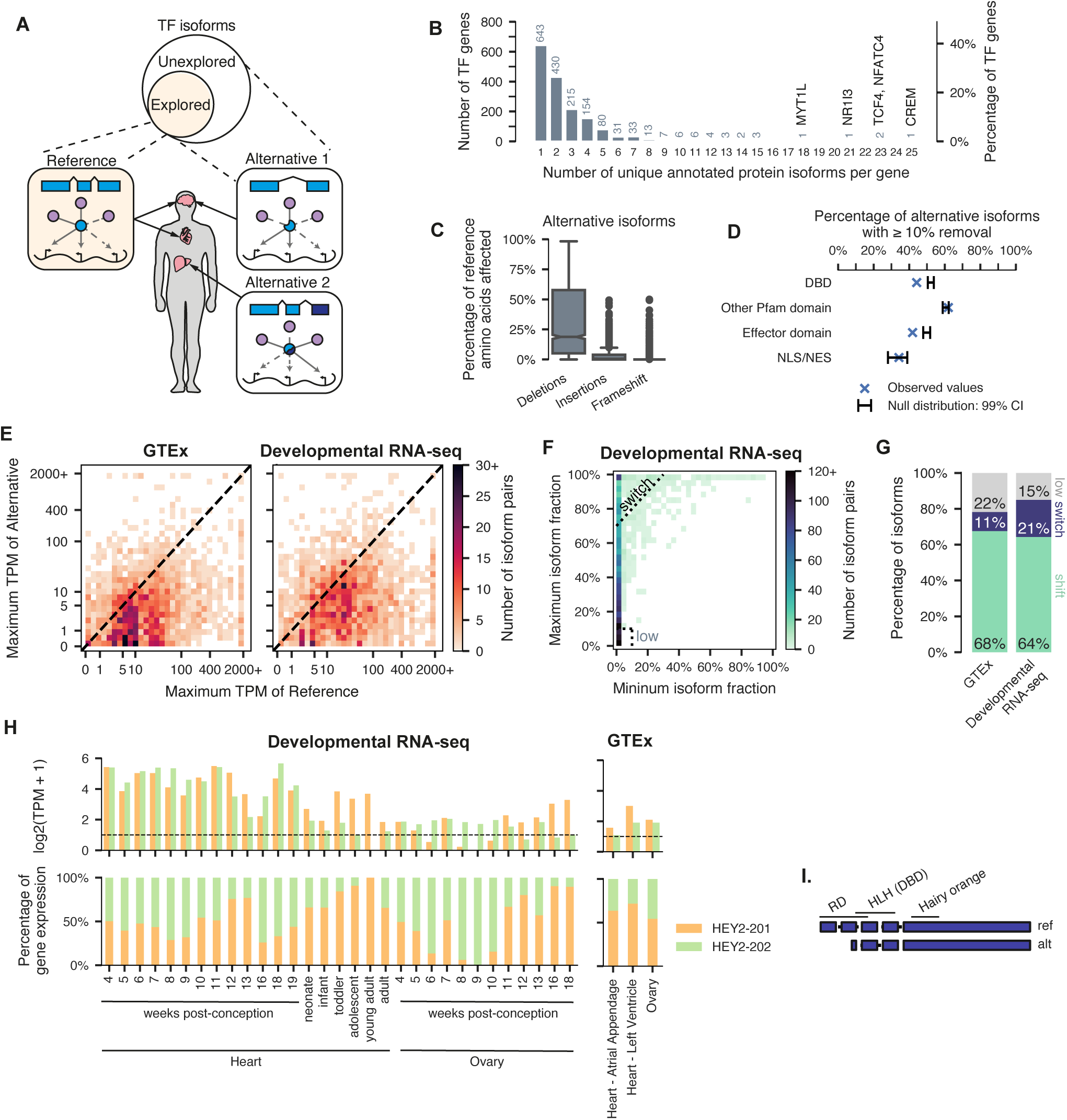
Sequence and expression diversity of annotated TF isoforms. **A.** Study schematic. **B.** Histogram showing the number of unique annotated protein isoforms for each TF gene. **C.** Boxplot showing the total percent of amino acids altered via deletions, insertions, or frameshifts in alternative isoforms compared to their cognate reference isoforms. **D.** Barplot showing the observed fraction of alternative isoforms with ≥ 10% removal of various protein domains (green bars) compared to the expected fraction (black error bars, 99% CI) as defined by a null model assuming the domain is randomly positioned along the protein. DBD = DNA-binding domain; NLS/NES = nuclear localization/export signal. **E.** Heatmap showing the maximum expression value of alternative TF isoforms (y-axis) compared to their cognate reference isoforms (x-axis) across GTEx (left) and developmental (right) RNA-seq datasets. GTEx dataset has been re-sampled to compare to the developmental dataset. **F.** Heatmap showing the maximum isoform fraction (y-axis) compared to the minimum isoform fraction (x-axis) of alternative TF isoforms in developmental RNA-seq data, where isoform fraction is defined as the expression level of an isoform normalized to the total expression level of its host gene. Dashed lines show the definitions used for isoforms that exhibit “switching” events and isoforms that remain lowly expressed. Only isoforms whose host genes are expressed at ≥ 1 TPM in ≥ 1 sample are shown. **G.** Stacked barplot summarizing the number of alternative isoforms defined to exhibit a switch event, a shift event, or be lowly expressed in re-sampled GTEx and developmental RNA-seq data. **H.** Example of an alternative TF isoform (HEY2-202) that exhibits a switch event. Top: log2 TPM values for each HEY2 isoform; bottom: isoform expression as a percentage of total gene expression for each HEY2 isoform. All heart and ovary samples in both GTEx and developmental RNA-seq are shown. Right: exon diagram of HEY2 isoforms with annotated protein domains. RD = repression domain; HLH = helix loop helix; DBD = DNA-binding domain.

Recent studies have globally investigated how coding variants in TFs–which are often associated with disease–affect TF functions such as DNA binding and transcriptional activity. Such mutations can range from having no detectable effect on TF functions to resulting in a complete loss or even gain of functions.^5,15,16^ However different transcript isoforms of common TF genes, herein abbreviated as “TF isoforms”, remain far less studied, despite being widespread. Indeed, TFs are among the most frequently spliced classes of gene^17,18^ and the majority of human TFs are present as multiple distinct isoforms.^19^ Current estimates show that the ~1,600 annotated human sequence-specific TF genes encode ~4,100 individual TF isoforms.^20,21^ This is likely a substantial underestimate of the “space” of isoform complexity, as novel disease-and condition-specific isoforms continue to be detected by long-read RNA-sequencing technologies.^21–25^ Importantly, recent high-coverage mass spectrometry studies suggest that the majority of frame-preserving isoforms are translated, highlighting the importance of studying their functional activities.^26^

Though the majority of TF isoforms remain uncharacterized, some individual TF isoforms have been shown to exhibit drastically different gene regulatory functions.^27,28^ Indeed, alternative TF isoforms can exhibit differential binding to DNA, cofactors, or chromatin-associated proteins,^18,27,29–31^ leading to isoform-specific effects on gene regulatory networks (GRNs). Notably, two isoforms of the Forkhead developmental regulator *FOXP1* gene exhibit different DNA binding specificities and consequently drive opposing phenotypes: one maintains pluripotency while the other drives differentiation.^32^ TF isoforms are also known to play distinct roles in disease. For example, altered expression of an alternative isoform encoded by the Wilms’ tumor *WT1* gene causes Frasier syndrome, a rare developmental kidney disease,^33^ and is also essential for female sex determination in mice.^34^ The alternative isoform differs by only 3 amino acids (a.a.) (-KTS) compared to the WT1 reference isoform, but differs substantially in DNA binding specificity.^35^ Moreover, TF isoforms can be dysregulated in cancer.^36^ Several TFs that act as either canonical oncogenes or tumor suppressors encode dominant negative TF isoforms that compete with reference isoform TF activity in the context of cancer, including STAT3,^37,38^ ESR1,^39,40^ and p53.^41^ For example, the alternative isoform of the known oncogene STAT3, STAT3beta, is missing the C-terminal transactivation domain due to an alternative splicing event; STAT3beta thus inhibits target gene activation by the reference isoform STAT3alpha and can act as a tumor suppressor.^37,38^

These case studies are striking, but few in number. This prompts the question of whether they are unusual scenarios, or represent a more general phenomenon of alternative isoforms diversifying the functions of TFs. In a previous systematic study of protein isoforms across the human proteome, we reported that isoforms exhibit widespread functional differences in their protein-protein interactions, suggesting that the latter hypothesis is more likely.^42^ However, relatively few studies have interrogated isoform-resolved TF functions, primarily due to technical limitations. For example, studies employing ChIP-Seq usually use antibodies that rarely distinguish between isoforms.^43^ Most large-scale studies of human TF DNA binding consider only the reference isoforms or just their DNA binding domains (DBDs).^2,3,44,45^ Additionally, high-throughput transcriptional activity studies have mostly been restricted to short effector domains or peptides, which can miss synergistic or antagonistic effects between domains within full-length isoforms.^6,7,46^ Overall, there is a need for high-throughput, integrative, experimental approaches to dissect the mechanisms by which alternative isoform usage alters the regulatory functions of TFs.

Here, we present an in-depth, experimentally driven investigation into the functional differences between 693 isoforms of 246 TFs. The results reveal system-scale relationships between TF sequence diversity and functional diversity, including DNA binding, transcriptional activation, protein-protein interactions, localization, and condensate formation. Our work builds on previous approaches^9,47^ to create “functional portraits” based on protein-protein and protein-DNA interaction networks to assess TF functions, extending them to assess multiple isoforms of individual TF genes. In this study, we present evidence that most alternative isoforms diversify the functions of TFs, provide a quantitative survey of the mechanisms involved, and propose that this rewiring of molecular functions through alternative isoforms constitutes an often overlooked but important layer of complexity in gene regulation in development and disease.

## Results

### TF isoforms are prevalent and frequently affect functional domains

To investigate the prevalence of TF isoforms that may play differential roles in GRNs, we began by cataloging annotated (known) protein-coding isoforms of TF genes (*i.e.*, transcripts of the same TF gene that encode different open reading frames, ignoring any changes in untranslated regions) using the GENCODE reference transcriptome database.^21^ GENCODE annotates 4,144 high-confidence protein-coding transcripts for the 1,635 human TF genes,^20^ with 992 TF genes (61%) encoding multiple isoforms (**Figure 1B**). On average, each TF gene encoded 2.5 protein isoforms; however, the number of isoforms per gene was highly variable, with extreme cases such as CREM, with 25 isoforms, and TCF4 and NFATC4, each with 23 isoforms. Nuclear hormone receptors have the highest number of annotated isoforms per gene (median = 3, mean = 3.8), whereas homeodomains have the lowest (median = 1, mean = 1.8) (**Figure S1A**). The number of annotated isoforms is likely an underestimate; with the advent of long-read RNA-sequencing platforms, studies in both healthy^23,48^ and diseased^49^ contexts have been uncovering thousands of novel transcripts. Indeed, recent work from the GTEx consortium revealed more than 70,000 novel transcripts across 88 human tissue samples and cell lines using Oxford Nanopore Technologies long-read sequencing.^48^ And a separate study of 30 human tumor and normal breast samples using Pacific Biosciences long-read sequencing revealed more than 94,000 novel transcripts.^49^

We next quantified the amino acid sequence diversity between TF isoforms, since sequence diversity may suggest functional divergence. For each TF gene, we therefore defined a “reference” isoform using the MANE Select representative transcript annotation set^50^ to serve as a baseline. We then compared all “alternative” isoforms of each TF gene to their cognate reference isoforms in a pairwise manner. Amino acid sequence differences between alternative TF isoforms and their cognate reference TF isoforms arise from alternative N-terminal regions, C-terminal regions, and/or alternatively spliced internal exons (**Figure S1B**). Across alternative TF isoforms, the median fraction of amino acids in the cognate reference isoform that is deleted is 18.8% (**Figure 1C**). While insertions and frameshifts are rare, 195 (8.5%) and 68 (3%) isoforms contain insertions or frameshifts affecting > 10% of their total amino acid length, respectively (**Figure 1C**).

Structural domains, such as DNA binding domains (DBDs), and generally unstructured effector domains are vital to the fundamental molecular functions of TFs, as they mediate specific biophysical interactions. Alternative TF isoforms with sequence changes in these annotated domains may therefore differ in their function. To determine the degree to which TF isoforms differ within and outside of annotated functional domains, we mapped three key protein domain types to TF isoforms: (1) annotated protein domains from the Pfam database, separated into DBDs and other domains (*e.g.*, ligand binding domains); (2) effector domains shown to either activate or repress transcription in reporter assays;^6–8^ and (3) nuclear localization/export signals (NLS/NES). Overall, 1,707 alternative TF isoforms (74%) differed by ≥ 1 amino acid in one of these annotated domains. Despite the frequency of domains being affected, however, DBDs and effector domains are statistically significantly affected slightly less than expected by chance, whereas NLS/NES motifs and other Pfam domains (*e.g.*, ligand binding domains, dimerization domains) are not (**Figure 1D, Figures S1C-D**). This finding supports previous reports that splicing boundaries tend to reside outside of annotated domains, perhaps reflecting a negative selection pressure to avoid deleterious alternative splicing variants,^51^ or reflecting evolutionary selection through which entire exons tend to be gained or lost.^52^

In the complex landscape of human gene regulation, understanding the expression patterns of alternative TF isoforms is critical for unraveling the contexts in which they may function. Given the lack of proteomics data that are of sufficient depth and quality to accurately quantify TF isoforms,^26^ which are typically expressed at < 30 transcripts per million (TPM), we used large-scale RNA-seq datasets to address this question. We used RNA TPM pseudo-aligned counts that map to a given unique protein isoform to infer isoform expression. We examined the expression patterns of TF isoforms across GTEx,^53^ which comprises primarily healthy human adult tissues, and a time-course series of human development across seven organs (hereafter referred to as “Developmental” RNA-seq).^54^ To correct for potential biases arising from imbalanced data when comparing the two datasets, we re-sampled GTEx to result in a dataset of equivalent size to the Developmental RNA-seq dataset (**Figure S1E, STAR Methods**). We calculated the maximum expression value for each TF isoform across tissues and developmental stages and, as expected, reference isoforms generally had higher values than alternative isoforms; however, alternative isoforms had higher values than their cognate reference isoforms in 522 (23%) and 551 (24%) cases, in the GTEx and Developmental RNA-seq datasets, respectively (**Figure 1E**).

A handful of TFs, including FOXP1,^32^ REST,^55^ and GRHL1,^56^ are known to dramatically “switch” from expressing one particular isoform to another at key stages of development. Such isoform switching events add an additional layer of complexity to the changes in GRNs that are required for cell state transitions. To determine the prevalence of “switch” events, we calculated the percentage of total TF gene expression for each TF isoform (the “fractional isoform expression”). We considered an alternative isoform to exhibit a “switch” event if it changed its fractional expression by at least 70% between any two conditions. We defined isoforms that did not reach at least 10% in any expression condition (22% in GTEx and 15% in Developmental RNA-seq) to have “low” fractional isoform expression and all other isoforms to exhibit more subtle “shifts’’ across conditions (**Figure 1F, Figure S1F**). The majority of TF isoforms (68% in GTEx and 64% in Developmental RNA-seq) exhibited these graded “shifts” across conditions rather than dramatic switching events (**Figure 1G**). Interestingly, the fraction of alternative isoforms that showed dramatic switches is higher in the Developmental RNA-seq data than in GTEx data (21% versus 10%, respectively), consistent with both TFs and alternative splicing being important regulators of differentiation^27^ and suggesting that many alternative TF isoforms might affect gene regulation in early development.

Overall, the vast majority of TF genes with multiple isoforms have at least one alternative isoform that exhibited either a “switch” or a “shift” event in GTEx (94%) and the Developmental RNA-seq data (96%). One example of a TF that shows variable isoform expression across tissues is HEY2, a Notch-dependent basic helix-loop-helix (bHLH) transcriptional repressor that is a critical regulator of cardiac development (**Figures 1H, 1I**).^57^ The alternative isoform of HEY2 arises due to an alternative transcription start site, which results in an open reading frame that lacks the N-terminal annotated repression domain (**Figure 1I**). While the relative expression of the two annotated HEY2 isoforms is similar across GTEx heart and ovary samples, the expression of these isoforms switches and shifts across the Developmental RNA-seq heart and ovary samples (**Figure 1H**). As expected, the reference isoform is more abundantly expressed in the heart, reaching 100% of total gene expression in young adult heart samples. However, in the ovary, another tissue where Notch signaling is known to play an important role,^58^ the alternative isoform of HEY2 is more abundant, reaching 100% of total gene expression in week 9 ovary samples (**Figure 1H**).

In summary, the majority of annotated alternative TF isoforms show differences in annotated protein domains and variable expression across tissues, particularly in the context of development. Taken together, our results imply that alternative TF isoforms might serve distinct roles in GRNs and thus underscore the need to functionally characterize TF isoforms.

### Systematic characterization of TF isoforms reveals differences in molecular interactions and regulatory activity

Given that TF isoforms exhibit differences in primary sequence, structural domains, and expression patterns, we hypothesized that alternative TF isoforms likely exhibit widespread functional divergence. To investigate the extent of human TF isoform diversity, we systematically assayed molecular functional differences (*e.g.*, biophysical interaction differences) across a large collection of TF isoforms. We first generated a clone collection of human TF isoforms, TFIso1.0, using a PCR-based approach to sample isoforms expressed in fetal and adult brain, heart, and liver tissues that have well-documented differences in splicing isoform expression^59^ (**Figure 2A**). Our clone collection comprises 693 protein-coding TF isoforms, corresponding to 246 TF genes and spanning a wide range of TF classes (**Figure S2A, Table S1**).

**Figure 2:**
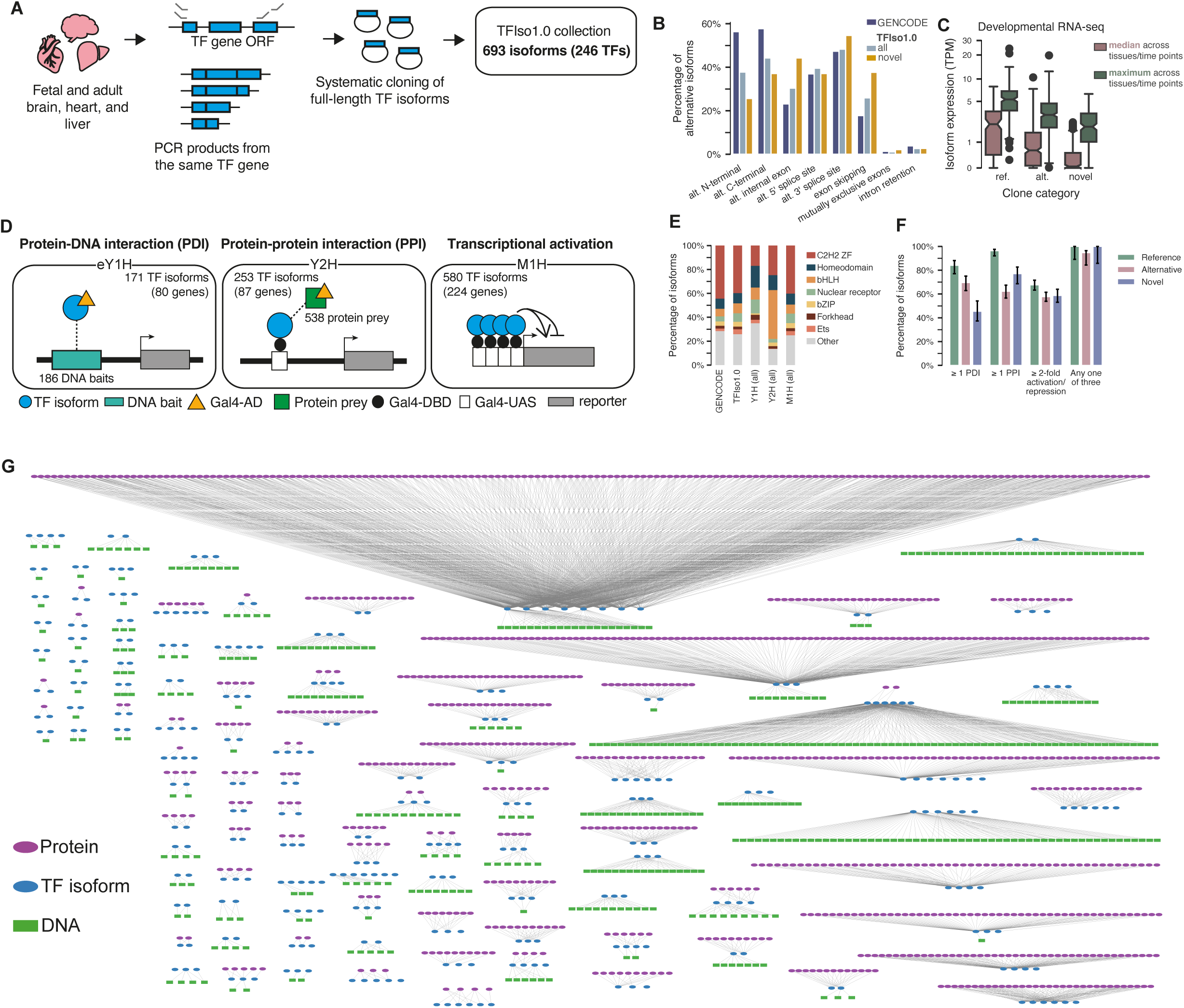
Overview of TFIso1.0 clone collection and TF molecular function assays. **A.** Schematic showing the PCR-based approach used to generate TFIso1.0. **B.** Barplot showing the percentage of alternative isoforms in GENCODE, all of TFIso1.0, and only the novel isoforms in TFIso1.0 exhibiting various sequence differences compared to their cognate reference isoforms. **C.** Boxplot showing the median and maximum expression levels (in TPM) in developmental RNA-seq data of reference, annotated alternative, and novel alternative isoforms in TFIso1.0. **D.** Schematic showing the three primary assays used in this study. eY1H = enhanced yeast one-hybrid; Y2H = yeast two-hybrid; M1H = mammalian one-hybrid; Gal4-AD = Gal4 activation domain; Gal4-DBD = Gal4 DNA-binding domain; Gal4-UAS = Gal4 upstream activation sequence. **E.** Stacked barplot showing the percent of TF isoforms belonging to various TF families in GENCODE, the entire TFIso1.0 collection, and those that have been successfully tested in each assay. **F.** Barplot showing the proportion of isoforms exhibiting ≥ 1 PPI, ≥ 1 PDI, ≥ 2-fold activation/repression in M1H, or any one of the three across reference, annotated alternative, and novel alternative isoforms, normalized to the number of isoforms that were successfully tested in each assay. Error bars are 68.3% Bayesian CI. **G.** The sub-networks of PPIs and PDIs from profiling different TF isoforms.

We compared the full-length sequences of the 693 isoform clones in TFIso1.0 to coding sequences in the GENCODE database,^21^ finding that 510 match existing transcripts, while 183 (26%) were novel. We manually curated these novel isoforms using standards established at GENCODE to ensure high quality (**STAR Methods**). Because our cloning strategy used annotated N-and C-terminal regions for primer design, we are likely to miss unannotated alternative transcription start and polyadenylation sites or unannotated splicing events in the UTRs, and consequently novel TF isoforms were more likely to differ in an internal exon and less likely to differ at the N-and C-terminals, compared to annotated alternative TF isoforms (**Figure 2B**). Although these novel isoforms are generally expressed at lower levels than annotated alternative isoforms, their maximum expression values across conditions are similar, indicating that these novel isoforms may be as-yet unannotated because they are expressed in a more tissue-or developmental-stage-restricted manner (**Figure 2C**, **Figure S2B-C**). For example, we discovered a novel isoform of ZNF414 that is expressed throughout liver development (**Figure S2D**). Uncovering these novel isoforms is an advantage of our PCR-based cloning strategy over approaches that rely on synthesizing ORFs based solely on existing annotations.^60,61^ The inclusion of novel isoforms in our clone collection necessitated a numbering system that expands upon GENCODE annotation: we refer to TF isoforms by their gene name and a clone ID number and supplement these identifiers with the matching GENCODE transcript name for annotated isoforms.

Experimentally solved 3D structures of alternative isoforms were exceedingly rare; therefore, to observe the differences in 3D structure between isoforms, we generated AlphaFold2 predictions for each of our cloned isoforms. TFs are enriched for intrinsically disordered regions (IDRs), particularly in their activation domains.^62^ TF IDRs have recently been implicated in phase separation^63,64^ and shown to affect DNA binding.^65^ We observed that alternative isoforms were predicted to have more of their sequence in IDRs than their reference isoforms (median proportion of residues in IDRs was 57% for alternative and 50% for reference isoforms; *P* = 0.0048, two-sided permutation test, **Figure S2E**).

To systematically characterize TF isoforms, we assessed DNA binding (or protein-DNA interactions, PDIs), transcriptional regulatory activities, and protein-protein interactions (PPIs) for isoforms in TFIso1.0 (**Figure 2D**; **STAR Methods**). We assessed TF-DNA binding using enhanced yeast one-hybrid (eY1H) assays,^4,66^ in which PDI profiles were generated by testing each TF isoform against a collection of 330 DNA-baits consisting of known developmental enhancer or gene promoter elements (**Tables S2, S3**). We assessed TF transcriptional regulatory activities using a modified mammalian one-hybrid (M1H) assay in HEK293T cells (**Table S4**). In M1H assays, full-length TF isoforms are tethered to a Gal4 DBD and transcriptional activity is then measured by co-transfecting with a plasmid containing four arrayed Gal4 upstream activation sequences (UAS) upstream of the firefly luciferase gene (**Figure 2D, Figure S2F**). We assessed PPIs using yeast two-hybrid (Y2H) assays, in which each TF isoform was systematically screened against the human ORFeome v9.1, comprised of 17,408 protein-coding genes,^67^ followed by pairwise testing of each TF isoform with all interaction partners for that TF gene. In total, we successfully tested 3,509 isoform-resolved protein pairs for interaction, where, in each case, at least one isoform of the tested TF gene interacts with the partner. The resulting PPI profiles involved 253 isoforms of 87 TF genes, tested against 538 different TF-binding protein partners (**Table S5**). All major TF families are well represented in the data produced by each of the assays (**Figure 2E, Figure S2G**). Binary PPI and PDI calls validated well when random samples were re-tested in orthogonal assays (**Figure S2H-K**; **Tables S6, S7**; **STAR Methods**), and M1H activities were highly reproducible across biological replicates (Spearman’s rank correlation coefficient = 0.99, **Figure S2L**). Novel TF isoforms showed evidence of functionality (in terms of numbers of DNA interactions, protein interactions, or ability to activate or repress transcription) at levels similar to annotated alternative isoforms (**Figure 2F, Figure S2M**), suggesting that these novel isoforms may serve important biological roles.

In total, we successfully assayed the PDIs, PPIs, and regulatory activities of 171, 253, and 580 TF isoforms, of 80, 87, and 224 genes, respectively. Our isoform-specific PDI and PPI network shows the long-tailed degree distributions typical of biological networks^68^ with a small fraction of TFs binding to a large number of interaction partners (**Figure 2G**). Altogether, our experimental dataset comprises the most comprehensive, systematic characterization of TF isoforms’ molecular interactions and regulatory properties reported to date. We therefore used the results of our survey to characterize the degree to which alternative TF isoforms differ from their cognate reference TF isoforms in terms of these key molecular functions.

### DNA binding of alternative TF isoforms is influenced by differences both inside and outside the DBD

One of the most important functions of sequence-specific TFs is to recognize and bind to short DNA motifs, thereby initiating the complex process of gene regulation. TF binding to DNA is canonically achieved through structured DBDs. We therefore first compared the PDI profiles of alternative TF isoforms to see how changes inside and outside the DBD affect DNA binding compared to their cognate reference TF isoforms (**Tables S2, S3**). Unsurprisingly, alternative TF isoforms that are completely missing the DBD completely lose the ability to bind DNA in our eY1H assay (**Figure 3A**). In almost every case, alternative TF isoforms that contain only a partial DBD also lose DNA binding. The one exception to this is ZIC3, which has two alternative isoforms that lose 3 amino acids in a C2H2 zinc finger DBD at the C-terminal end of an array of five zinc fingers (**Figure S3A**). These alternative ZIC3 isoforms bind different subsets of DNA baits than the reference isoform (**Figure S3A**). One of these alternative ZIC3 isoforms, known as ZIC3-B, is conserved in mouse and co-expressed with the reference isoform in development,^69^ suggesting that these isoforms may perform distinct roles in GRNs.

**Figure 3:**
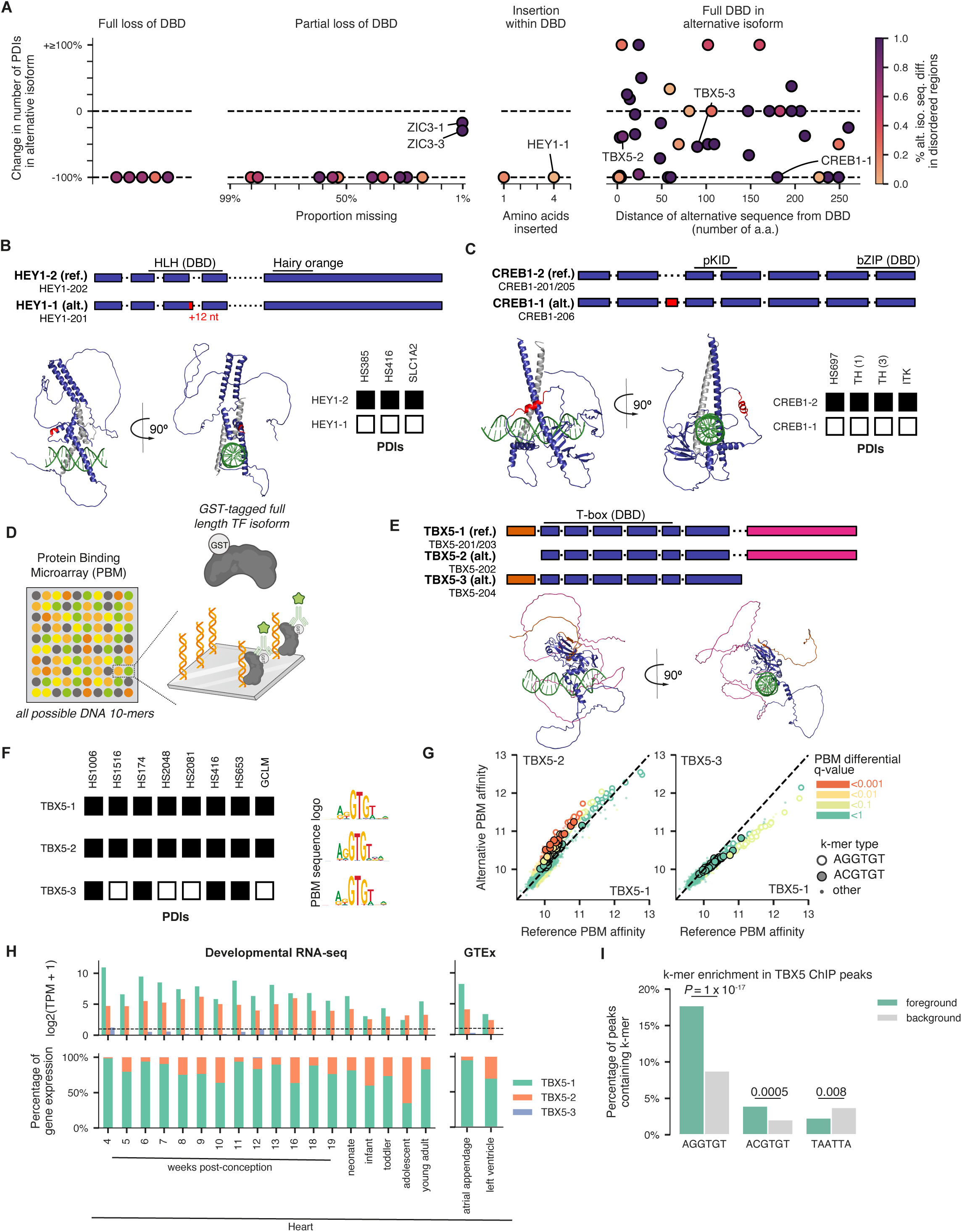
DNA binding preferences of TF isoforms. **A.** Change in the number of PDIs in the alternative isoform compared to the reference isoform for alternative isoforms with full loss of DBD, partial loss of DBD, insertions within the DBD, or that contain the full DBD. Each point is colored by the percentage of the sequence difference in the alternative isoform stemming from a predicted disordered protein region. **B.** Top: exon diagrams of cloned HEY1 isoforms with annotated Pfam domains. 12 nt = location of the 4 amino acid insertion in the alternative isoform, at the end of exon 3. Bottom left: AlphaFold model of the alternative isoform of HEY1 aligned to an experimental structure of a homologous protein in a dimer bound to DNA (PDB ID 4H10), with DNA in green and dimerization partner in gray. Bottom right: PDI results from Y1H assay for the 3 baits successfully assayed for both isoforms of HEY1; black box = binding and white box = no binding. **C.** Top: exon diagrams of CREB1 isoforms with annotated Pfam domains. pKID = phosphorylated kinase-inducible domain. Bottom left: AlphaFold model of the alternative isoform of CREB1 aligned to an experimental structure of a CREB1 homodimer bound to DNA (PDB ID 1DH3), with DNA in green and dimerization partner in gray. Bottom right: PDI results from the Y1H assay for the 4 baits successfully assayed for both isoforms of CREB1. **D.** Schematic showing protein-binding microarray (PBM) experiments. Scores for all possible 8-mers are calculated from universal “all 10-mer” PBMs. **E.** Top: exon diagrams of TBX5 isoforms with annotated DBD. 3 nt = TBX5-2 is missing 1 amino acid at the start of its first exon compared to the reference. Bottom: AlphaFold model of the reference isoform of TBX5 aligned to an experimental DNA-bound structure (PDB ID 5FLV), with DNA in green. **F.** Left: PDI results from the Y1H assay for the 3 isoforms of TBX5. Showing 8 baits that were successfully tested against all 3 isoforms. Right: Sequence logo derived from the top 50 8-mers as determined via PBMs for each of the 3 TBX5 isoforms. **G.** Scatter plots showing the PBM affinity scores for the alternative isoform (y-axis) compared to the reference isoform (x-axis) of TBX5 for every 8-mer, for either TBX5-2 (left plot) or TBX5-3 (right plot), each compared to TBX5-1. Points are colored by the differential affinity q-value calculated by the upbm package.^78^ Open circles correspond to 8-mers containing the canonical TBX5 6-mer AGGTGT (or its reverse complement); filled circles correspond to 8-mers containing the altered 6-mer ACGTGT (or its reverse complement). **H.** Expression of TBX5 isoforms in developmental RNA-seq (left) and GTEx (right). Top: log2 TPM values for each TBX5 isoform; bottom: isoform expression as a percentage of total gene expression for each TBX5 isoform. All heart samples are shown. **I.** Barplot showing the enrichment of the canonical TBX5 6-mer AGGTGT, the altered TBX5 6-mer ACGTGT, or a negative control Homeodomain 6-mer TAATTA (or each of their reverse complements) in TBX5 ChIP-seq peaks (foreground) compared to matched genomic negative control regions (background). P-values shown are from a Fisher’s exact test.

Only two alternative isoforms for which we have PDI data have insertions within their DBD, and both completely lose PDIs. One of these is an alternative isoform of the transcriptional repressor HEY1, which contains a four amino acid insertion in a short alpha-helical section within the loop region of the bHLH DBD. Previous work has shown that the loop region of MLX–another bHLH TF with a long (> 14 a.a.) loop region–is important for stabilizing complexes of bHLH dimers.^70^ The alternative isoform of HEY1 with this insertion fails to bind to any of the three DNA baits that the reference isoform of HEY1 binds (**Figure 3B**). This longer alternative isoform of HEY1 shows lower repression of the dopamine transporter *DAT1* than the reference isoform, despite maintained localization to the nucleus.^71^ Our results are consistent with previous studies that found that disruption of a DBD by the insertion of only a few amino acids can have strong effects on TF function; for example, a three amino acid insertion within the C2H2 zinc finger array of WT1 was previously shown to completely abrogate DNA binding.^72,73^

Most of the assayed alternative TF isoforms, 42/63 (67%), contained the complete, unaltered DBD (**Figure 3A**, right). However, only 8/42 (19%) showed identical DNA binding profiles to their cognate reference isoforms. Moreover, 9 of these 42 alternative isoforms (21%) containing the full, unaltered DBD gained PDIs that their cognate reference isoforms lack (**Figure S3B**). We find that sequence differences in regions close to DBDs are often associated with dramatic differences in DNA binding, consistent with evidence that flanking regions can play a pivotal role in TF-DNA binding;^74^ alternatively, this may suggest uncertainty in the prediction of exact domain boundaries.^75^ Surprisingly, however, sequence differences in regions far from the DBD and commonly in IDRs (**Figure 3A**, right panel) often affect DNA binding. For example, 13/20 (65%) alternative isoforms with sequence differences that are at least 100 a.a. away from the DBD have differences in their DNA binding, and of those, in 9/13 (69%) cases, the sequence differences are in regions predicted to be fully disordered.

One striking example of sequence differences in IDRs far from the DBD affecting DNA binding is seen for the TF CREB1. An alternative isoform of CREB1 differs from the reference isoform by the inclusion of a small, in-frame 14-a.a. exon, 165 amino acids N-terminal of the bZIP DBD. This 14-a.a. exon is predicted by AlphaFold2 to be in a long disordered region (**Figure 3C**). We observed a complete loss of binding for this alternative isoform of CREB1 across the approximately 500-2,000-bp DNA sequences assayed by Y1H (**Figure 3C**). This alternative isoform of CREB1 retains its ability to activate transcription of the reporter gene in the M1H assay, suggesting that the alternative isoform is expressed and folded (**Figure S3C**). Reasoning that the loss of eY1H DNA binding in the alternative isoform of CREB1 might be due to differential DNA binding affinity or specificity between the two CREB1 isoforms, we performed *in vitro* universal protein binding microarrays (PBM) using full-length CREB1 proteins (**Figure 3D, Table S8, STAR Methods**).^76,77^ Universal PBMs provide a measure of the relative affinity of a protein for all possible 8-bp sequences, allowing for higher resolution measurement of the TF isoform’s sequence preferences.^78^ The alternative isoform of CREB1 showed lower affinity for DNA than the reference CREB1 isoform, particularly for the 8-mers that comprise the canonical CREB1 binding motif. However, this relative affinity difference was subtle (**Figure S3D**), suggesting that small differences in DNA binding affinity may lead to marked changes in binding to longer DNA targets, resulting in binding signal below the sensitivity of the eY1H assay. It is possible that the alternative isoform of CREB1 may still retain binding to other DNA targets that were not assayed by eY1H.

Another TF that shows differences in IDRs across isoforms is TBX5. TBX5 is a member of the Brachyury family and a critical regulator of heart development.^79^ Mutations in TBX5 cause the developmental disease Holt-Oram syndrome.^80^ There are three annotated isoforms of TBX5, all of which are represented in TFIso1.0 (**Figure 3E**). One alternative isoform of TBX5, TBX5-2, also known as TBX5e, differs from the reference isoform, also known as TBX5a,^81^ in the disordered N-terminal region of the protein, adjacent (but outside of) the T-box DBD. The other alternative isoform, TBX5-3, differs from the reference isoform in the disordered C-terminal region of the protein, affecting an annotated activation domain but distal to the DBD. TBX5-3 loses binding to half (4 of 8) of the TBX5 reference DNA baits in eY1H, whereas TBX5-2 retains binding (8 of 8) (**Figure 3F**). Given the biological importance of TBX5, we further profiled these full-length isoforms using PBMs (**Table S9**). Consistent with the eY1H assay, we see that TBX5-3 has slightly lower affinity for most 8-mers on PBMs (**Figure 3G**). Interestingly, while TBX5-2 and the reference TBX5 isoform have similar affinity for the highest affinity 8-mers that include the canonical TBX5 motif AGGTGT, TBX5-2 shows significantly higher affinity than the reference TBX5 isoform for a subset of moderate affinity 8-mers (**Figure 3G**). TBX5-2 is co-expressed with the reference TBX5 isoform throughout heart development (**Figure 3H**). Analysis of human TBX5 ChIP-seq data in cardiomyocytes^82^ showed that these moderate affinity 8-mers are highly enriched in TBX5 binding sites (**Figure 3I, Figure S3E**), indicating that these 8-mers, which are preferentially bound by the alternative isoform TBX5-2 *in vitro*, may be playing an important role *in vivo*. Future isoform-specific ChIP-seq will be important to disentangle the roles of these two isoforms in regulating cardiac GRNs.

Altogether, our results support recent findings that IDRs of TFs, outside of the structured DBD, play critical roles in determining TF DNA binding, particularly in the context of a chromatinized genome.^65,83^ Moreover, our work highlights that changes in DNA binding are extremely challenging to predict from sequence (and predicted structure) alone. Thus, it remains highly necessary to perform complementary DNA binding assays such as eY1H and PBMs to more fully elucidate the DNA binding activities of TFs.

### TF isoforms often differ in transcriptional activities due to changes in annotated and putative novel effector domains

In addition to changing the DNA targets of a TF, alternative isoforms can also rewire GRNs by altering transcriptional activity through changes in PPIs with cofactors, other TFs, or signaling proteins. We therefore measured the extent of differences in transcriptional activity between the reference and alternative isoforms using M1H assays (**Table S4**). We annotated our cloned TF isoforms with activation and repression domains from literature-curated and systematic experimental datasets.^6–8^ As expected, TF isoforms with transcriptional activity above basal levels in the assay are enriched in isoforms containing activation domains, whereas TF isoforms with activity below basal levels are enriched in isoforms containing repression domains (**Figures S4A-B**). Overall, 125 of the profiled alternative isoforms (49%) showed at least a 2-fold difference in M1H activity compared to their cognate reference isoform, with more alternative isoforms losing, rather than gaining, activity (**Figure S4C**). Moreover, we found that four TF genes (FOXP3, MAX, MAZ, ZNF544) encode both activator and repressor isoforms. For example, while the reference isoform of FOXP3 is a repressor, the four alternative isoforms are weak activators (**Figure S4D**). Thus, TF isoforms can differ greatly in their transcriptional activities, leading to potentially different effects on gene regulation.

As expected, alternative TF isoforms that have full or partial loss of activation domains often showed reduced transcriptional activity compared to their cognate reference isoforms (*P* = 0.02, paired two-sided Wilcoxon test, **Figure 4A**, **Figure S4E**). In contrast, the overall effect of partial or full loss of repression domains was less clear (**Figure 4A**), potentially because of (1) lower sensitivity to detect repression in the version of M1H assay used, which employs a minimal promoter with low background activity, (2) the cellular context of the assay in which some corepressors may not be expressed or active, or (3) dominance of other (potentially unannotated) effector domains. Alternative isoforms that show loss of both annotated activation and repression domains tended to lose transcriptional activity, suggesting a more dominant effect of activation domains (**Figure 4A**). For example, an alternative isoform of E2F3 that loses an entire annotated activation domain (E2F3-2) has, as expected, strongly decreased activity, whereas a second alternative isoform of E2F3 (E2F3-4) that loses most of an annotated repression domain does not show increased activity (**Figure S4F**).

**Figure 4:**
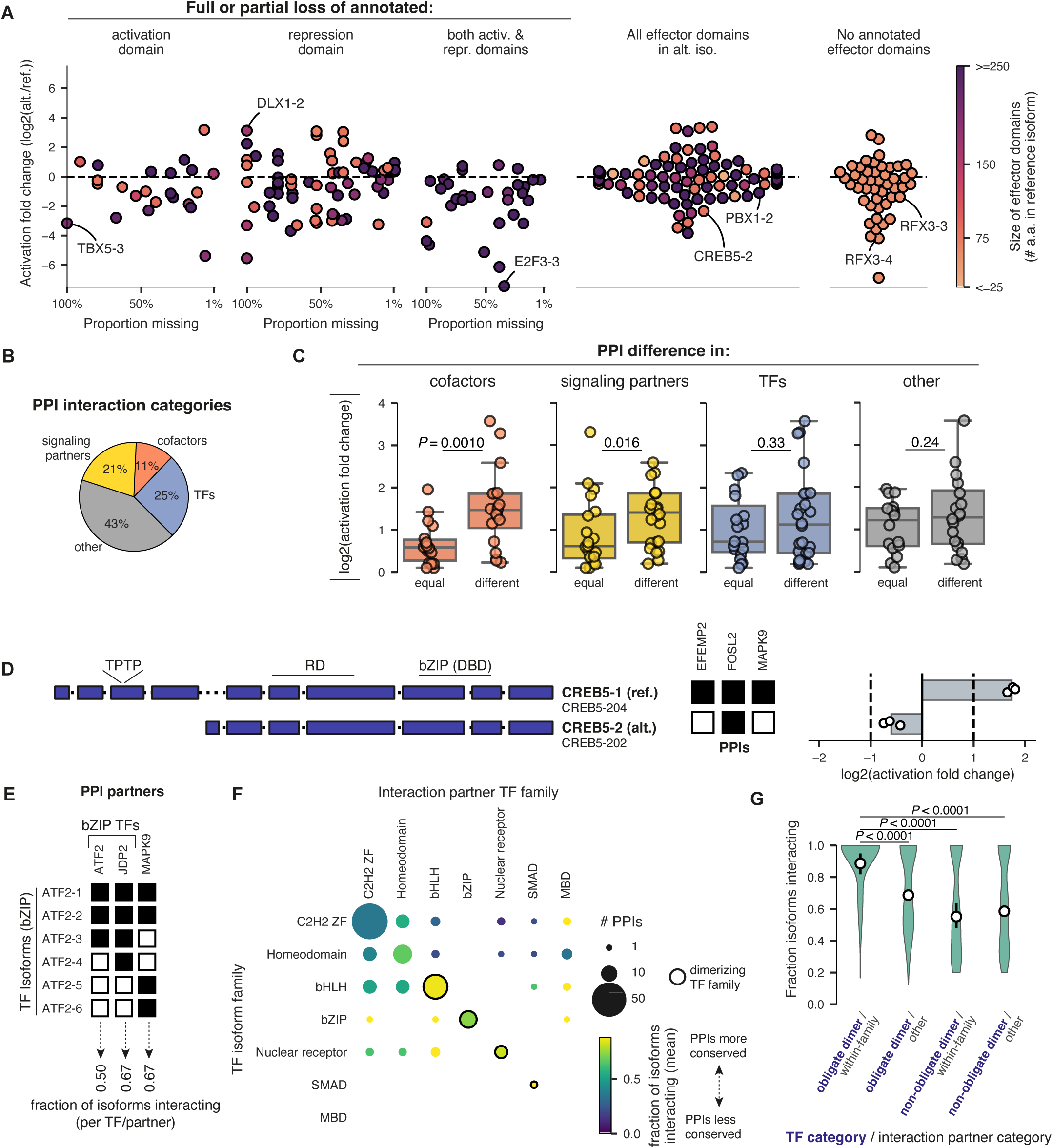
Transcriptional activity and protein binding preferences of TF isoforms. **A.** Summary plot showing the change in the transcriptional activity (log2 fold-change, as determined via M1H assays) of the alternative isoform compared to the reference isoform for alternative isoforms with full or partial loss of annotated activation or repression effector domains, no loss of annotated effector domains, or containing no annotated effector domains. Each point is colored by the total number of amino acids in annotated effector domains for a given isoform. **B.** Pie chart showing the categories of PPI partners (as determined via Y2H assays) found to interact with ≥1 TF isoform. **C.** Box plots showing the absolute change in transcriptional activity associated with no change in PPIs (equal) or a change in PPIs (change) for various categories of PPI partners. P-values shown are from a one-sided Mann Whitney U test. **D.** Left: exon diagrams of CREB5 isoforms. RD = repression domain; 18 nt = CREB5-1 reference isoform clone is missing 6 amino acids at the N-terminus compared to annotated CREB5-204. Middle: PPI results from the Y2H assay for the 2 isoforms of CREB5; black box = binding and white box = no binding. Right: transcriptional activity from the M1H assay for the 2 isoforms of CREB5. **E.** Schematic showing how to calculate the fraction of isoforms interacting, using the PPI results for the 6 isoforms of ATF2. Showing PPI partners that were successfully tested against all 6 isoforms. **F.** Heatmap showing the rewiring score for combinations of families of TF isoforms (y-axis) and families of TF PPI partners (x-axis). Within-family dimerizations are therefore denoted on the diagonal of the heatmap. TF families that bind DNA as obligate dimers are marked with outlined black circles on the diagonal. The size of the circle denotes the number of PPIs, whereas the color denotes the mean fraction of isoforms interacting. Only TF isoform families with ≥3 TF partner interactions are shown; for the full heatmap see **Figure S4I**. **G.** Violin plot showing the fraction of TF isoforms (categorized before the slash in bolded blue text) that retain interactions with various TF PPI partner types (categorized after the slash). P-values shown are from a two-sided permutation test.

We observed multiple instances of isoforms showing marked differences in transcriptional activity that did not differ in any annotated effector domains (**Figure 4A**). This highlights the incompleteness of current effector domain annotations, many of which are determined by testing short protein segments in reporter assays that tile across TFs and cofactors in a limited number of cell types.^6,7^ Such approaches may miss effector domains that require the full protein context for their function. For example, an alternative isoform of RFX3, RFX3-4, which lacks the C-terminal domain, loses transcriptional activity relative to the reference isoform, while an additional alternative isoform which largely retains this domain also retains activity (**Figure S4G**). This suggests the presence of an activation domain in the C-terminal region of RFX3 that was not detected in previous tiling screens^6,7^–either due to cell-type dependent activation or requirement of the full protein context–and illustrates how our profiling of full-length TF isoforms can be used to identify putative effector domains.

### Changes in PPIs with cofactors and signaling proteins are associated with differences in activity between TF isoforms

We hypothesized that differences in transcriptional activity between isoforms likely result from differences in PPIs. We generated isoform-resolved PPI profiles, testing multiple isoforms of TF genes against a single isoform of protein interaction partners of those TF genes. In total, we successfully tested 3,509 isoform-resolved protein pairs for interaction, where, in each case, at least one isoform of the tested TF gene interacts with the partner, corresponding to 936 PPIs at the gene-gene level (**Table S5**). Of the gene-gene level PPIs, 684 (73%) varied across isoforms. We were able to predict the two interacting domains^84^ for 152 PPIs (16%) that involved the reference isoform and test the association of the disruption of the domain with disruption of the PPI in the alternative isoforms. Complete loss of the binding domain always resulted in loss of the corresponding PPIs and changes outside the domain often resulted in loss of PPIs (**Figure S4H**), similar to the effects of DBD disruption on PDIs (**Figure 3A**). We next focused on 3 major classes of PPI partners that are likely to affect transcriptional activity: (1) transcriptional cofactors,^85^ such as chromatin remodelers and histone-modifying enzymes; (2) signaling partners (**STAR Methods**), such as kinases and other post-translational modifying enzymes; and (3) TFs (**Figure 4B**). We found that changes in cofactor binding among TF isoforms are associated with strong changes in transcriptional activity (**Figure 4C**). Changes in signaling partner binding are also associated with changes in transcriptional activity across isoforms. For example, an alternative isoform of CREB5 has strongly reduced transcriptional activity compared to the reference isoform (**Figure 4D**). The alternative isoform loses interaction with 2 partners: EFEMP2, an extracellular matrix protein that is not predicted to affect transcriptional activity, and MAPK9 (also known as JNK2), a key signaling kinase (**Figure 4D**). Indeed, the alternative isoform of CREB5 is missing two consecutive, conserved threonine-proline motifs (T59-P60 and T61-P62) present in the reference isoform, that are known to be substrates for the JNK family of kinases,^86^ and which have been found to be phosphorylated in multiple independent experiments.^87^ Our results therefore suggest that phosphorylation at these sites may be important for the transcriptional activity of the reference isoform of CREB5, as is known to be the case for CREB1.^88^

### TFs that bind DNA as obligate dimers tend to maintain intra-family protein-protein interactions across isoforms

TF function often relies on PPIs with other TFs, as many TFs bind to DNA as dimers or multimers. Some TF families, such as bZIPs and bHLHs, bind to DNA as obligate dimers.^20^ We therefore tested whether TF isoforms within obligate dimer TF families are more likely to retain within-family dimerizing interactions compared to other families, such as C2H2 zinc finger TFs, which predominantly bind as monomers. For every TF gene in our clone collection and every PPI partner found to interact with at least 1 isoform of that TF gene, we calculated the fraction of isoforms of that TF that interacted with the PPI partner. For example, 50% (3/6) of ATF2 isoforms are able to homodimerize with the reference isoform of ATF2; 67% (4/6) of ATF2 isoforms interact with TF partner JDP2; and an overlapping but nonidentical 67% of ATF2 isoforms interact with MAPK9 (**Figure 4E**). Of the within-family TF-TF PPIs tested, 94% are heterodimers and only 6% are homodimers. We found that, on average, interactions between TFs of the same family are more often retained across isoforms for obligate dimer TF families than other TF families (**Figures 4F-G, Figure S4I**). By definition, obligate dimer TFs require these dimerizing PPIs in order to bind DNA; therefore, taken together with the observation that DBDs tend to be preserved (**Figure 1D**), there appears to be a selective pressure on alternative TF isoforms to retain DNA binding function.

### TF isoforms can be as distinct in molecular functions as TF paralogs

Gene duplication and alternative splicing are two different processes that can each produce novel proteins (**Figure 5A**). The interplay between these two processes has been studied at the level of the genome and transcriptome,^89,90^ but outside of a few examples,^91–93^ little is known about how these processes compare in their effect on the molecular functions of proteins. We therefore evaluated how paralogous TFs (always comparing the two reference isoforms with each other) compare to TF isoforms in TFiso 1.0 (always comparing an alternative isoform with its cognate reference isoform) in terms of their molecular functions. We found that our measured PPI profiles, PDI profiles, and activation levels are more similar between alternative and reference isoforms of the same TF gene than they are between reference isoforms of paralogous genes, which are in turn more similar than reference isoforms of non-paralogous TFs (**Figures 5B-D, Tables S10, S11**). However, these observations are confounded by sequence similarities between isoforms and paralogs: indeed, on average, paralogs tend to vary more at the sequence level than isoforms (**Figure 5E**). When controlling for these overall differences in sequence similarities, isoforms tended to show similar differences in their molecular functions compared to paralogs (**Figures 5F-H, Figures S5A-C**). In fact, TF isoforms tended to show more dramatic differences in PDIs than paralogous TFs.

**Figure 5:**
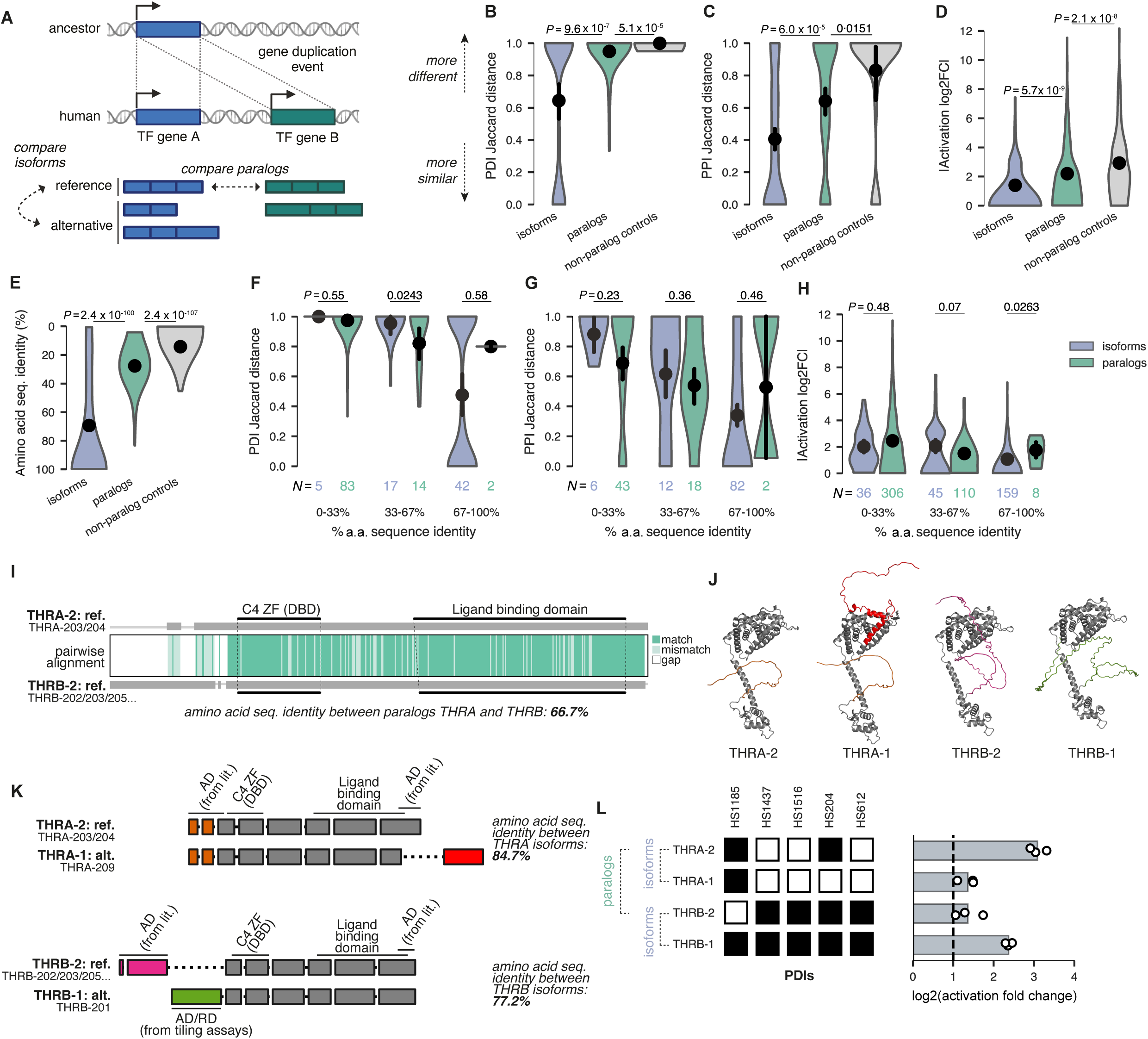
Functional differences between TF isoforms and TF paralogs. **A.** Schematic showing the definition of TF paralogs (blue vs. green) compared to TF isoforms (blue series or green series). **B-C.** Violin plots showing the Jaccard distance in PDIs (B) and PPIs (C) across reference/alternative isoform pairs, reference paralog pairs, or non-paralog reference pairs as a negative control. A Jaccard distance of 0 corresponds to entirely similar binding profiles, whereas a Jaccard distance of 1 corresponds to entirely dissimilar binding profiles. **D.** Violin plot showing the absolute log2 fold-change in M1H activation between isoforms, paralogs, and non-paralog controls. **E.** Violin plot showing the amino acid sequence identity (note that 100% identity is at the bottom of the y-axis, to remain consistent with the other plots) between isoforms, paralogs, and non-paralog controls. **F-H.** Analogous to B-C, but with isoform and paralog pairs broken up into bins based on their amino acid sequence identity. Number of pairs in each bin are denoted below the violins. **I.** Middle: pairwise sequence alignment of the reference isoforms of paralogs THRA and THRB, with darker green denoting perfectly matched amino acids and lighter green denoting mismatched amino acids. White regions indicate a gap in the alignment, and the gray schematics above and below the colored alignment denote which sequence is considered (thick gray block) or gapped (thin gray line). DBD and hormone receptor domains are denoted in each of the two paralogs. C4 ZF = C4 zinc finger. Right: AlphaFold2 predicted structures for isoforms of THRA and THRB. **J.** Exon diagrams of THRA isoforms (top) and THRB isoforms (bottom). AD = activation domain; RD = repression domain **K.** Left: PDI results from the Y1H assay for the isoforms of paralogous TFs THRA and THRB. Right: Transcriptional activity from the M1H assay.

For example, there are two thyroid hormone receptor genes in humans, *THRA* and *THRB*. These paralogous TFs evolved from an ancestral gene that was duplicated in vertebrates 500 million years ago.^94^ THRA and THRB share an amino acid sequence identity of 66.7%, and are particularly conserved within both their C4 zinc finger DBDs and their hormone receptor domains, with the largest differences between their unstructured N-terminal effector domains (**Figure 5I**). We have cloned alternative isoforms of both THRA and THRB, each alternative isoform affects an annotated effector domain; the alternative isoform of THRA differs from its cognate reference isoform at the C-terminal activation domain, and the alternative isoform of THRB differs from its cognate reference isoform at the N-terminal activation domain (**Figure 5J, K**). We found that the differences in PDIs were more subtle between the reference and alternative isoforms, THRA-1 vs THRA-2 and THRB-1 vs THRB-2, than between the paralogous reference isoforms, THRA-2 vs THRB-2 (**Figure 5L**, left). In contrast, the alternative isoforms and paralogous reference isoforms both showed strong differences in transcriptional activity, with THRA-1 having a lower level of activation than THRA-2, and THRB-1 being a stronger activator than THRB-2 (**Figure 5L**, right). Thus, both gene duplication and alternative splicing have modulated the molecular functions of the human thyroid hormone receptors, which is consistent with the model that both evolutionary mechanisms have affected this important metabolic pathway in mammals.^95^

To further explore the differences between TF isoforms and paralogs, we investigated the largest TF family, the C2H2 zinc finger proteins. In principle, the modular nature of individual zinc fingers in an array could enable alternative isoforms to exhibit different DNA binding specificities by splicing individual zinc fingers in or out. However, we found that alternative isoforms of TFs containing zinc finger arrays generally either completely preserve (67%) or completely remove (25%) the entire zinc finger array (**Figure S5D**). This is in stark contrast to zinc finger TF paralogs, which have been shown to alter DNA binding due to differences in the number and spacing of zinc fingers.^74^ Altogether, our results support prior studies that found that, compared to gene duplication, alternative splicing results in sequence changes that are more concentrated within specific regions of proteins and are predicted to affect physico-chemical protein properties more dramatically.^96^

### Widespread differences in cellular localization and condensate formation between isoforms

Eukaryotic gene regulation involves the organization of DNA, RNA, and transcriptional machinery into nuclear condensates, which spatially sequester macromolecules into regions of local enrichment.^97^ The formation of many nuclear condensates is aided by TF IDRs and associated with increased gene activation^63^ and pioneering activity.^98^ We therefore sought to determine whether isoforms of TFs are differentially able to contribute to condensate formation in mammalian cells. To do this, we expressed monomeric, enhanced green fluorescent protein (mEGFP)-tagged forms of 189 isoforms across 60 TF genes in HEK293T and U2OS cells and evaluated both their subcellular localization and ability to form condensates using high-throughput confocal fluorescence microscopy (**Figure 6A, B**).^99^ We focused on a subset of TFs that were found to show differences in either PDIs, PPIs, or transcriptional activation across isoforms in our systematic molecular function assays (**Figure 6B**, **Table S12**).

**Figure 6:**
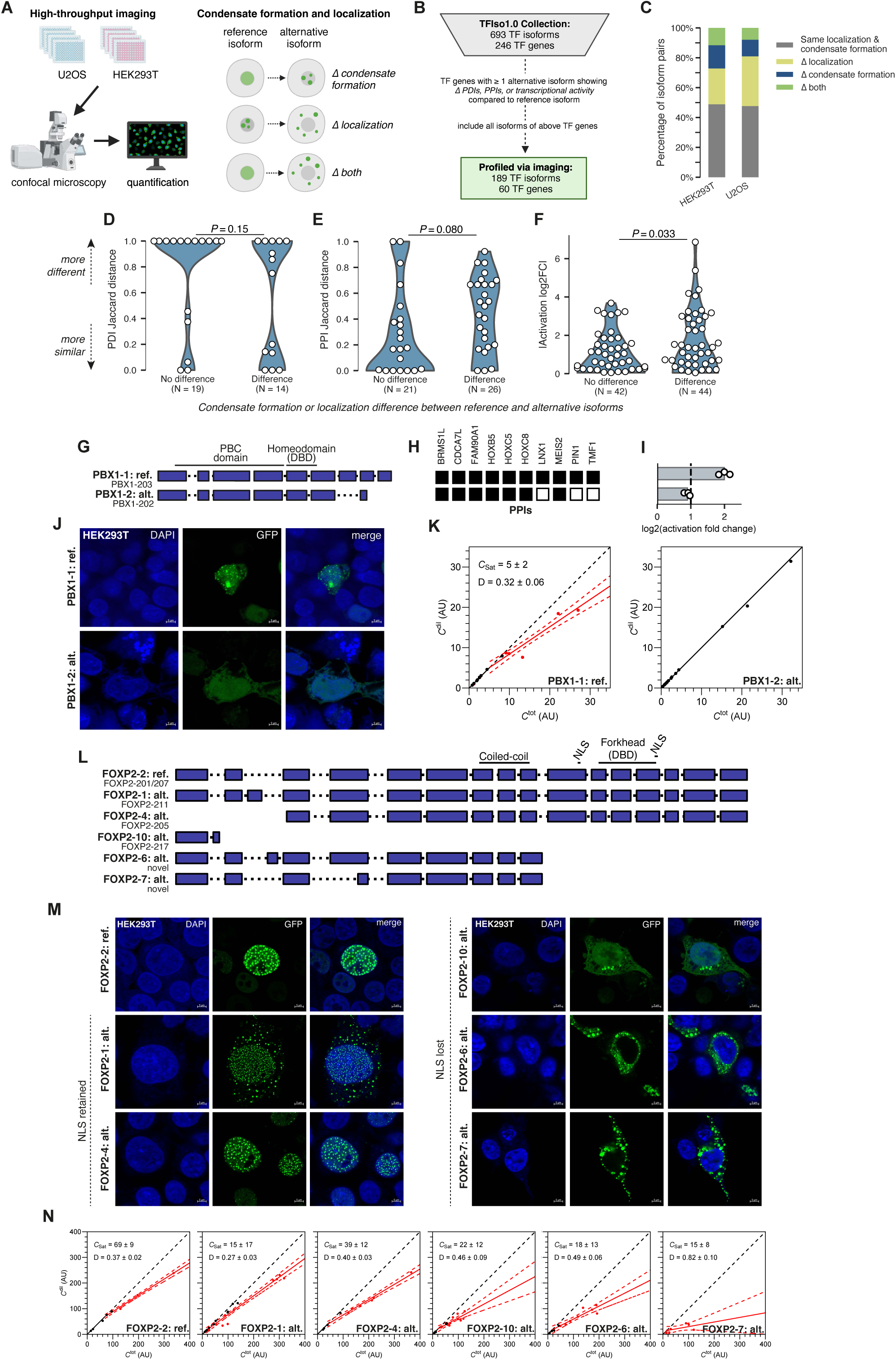
Condensate formation and subcellular localization differences between TF isoforms. **A.** Schematic showing the assessment of condensate formation and subcellular localization using high-throughput imaging across 2 cell lines, HEK293T and U2OS. **B.** Description of the TF isoforms that were selected for profiling in the high-throughput imaging assays. **C.** Stacked bar plot showing the percent of alternative isoforms that show differences in localization, condensate formation, both, or neither as compared to their reference isoform in either HEK293T or U2OS cells. **D.-F.** Violin plots showing the differences in TF molecular functions (PDIs, **D**; PPIs, **E**; transcriptional activity, **F**) between alternative-reference TF isoform pairs that either show no difference in condensate formation or localization or those that do. For these analyses, only TF isoform pairs with consistent results across the two imaging cell lines were considered. P-values calculated using a two-sided permutation test. **G.** Exon diagram showing the two cloned isoforms of PBX1 in TFIso1.0, with Pfam domains annotated. **H.** Y2H PPI results for the two isoforms of PBX1. **I.** M1H transcriptional activation results for the two isoforms of PBX1. **J.** Representative images of PBX1 isoform expression in HEK293T cells (63x magnification). **K.** Saturation (Csat) curve analysis of PBX1 isoforms. Dots represent individual cells, x-axis shows total protein concentration from fluorescence (C^tot^), y-axis shows concentration in the dilute phase (C^dil^). Arbitrary units (AU) are at reference settings. C_sat_ = saturation concentration; D = dominance. **L.** Exon diagram showing the six cloned isoforms of FOXP2 in TFIso1.0, with Pfam domains and nuclear localization sequence (NLS). **M.** Representative images of FOXP2 isoform expression in HEK293T cells (63x magnification). **N.** Csat analysis of FOXP2 isoforms.

Our observed localization of reference isoforms agreed well with the localization of the same proteins in the Human Protein Atlas,^100^ which measured localization using endogenous immunofluorescence (**Figure S6A**). Additionally, there are no significant differences in endogenous expression levels, from RNA-seq, between reference isoforms that form condensates in our assay compared with those that do not (**Figure S6B**). Altogether, these analyses suggest that our observed results are not an artifact of the exogenous expression system used in our high-throughput microscopy assay.

In both cell lines, approximately 50% of these alternative isoforms showed differences in either condensate formation or localization (**Figure 6C**, **Figures S6C-G**). Alternative isoforms that showed differences in condensate formation or localization compared to their cognate reference isoforms tended to also show differences in transcriptional activity (**Figures 6D-F**). This agrees with recent studies showing that the formation of nuclear condensates plays an important role in gene activation.^101,102^ We did not observe a significant association between localization/condensates and differences in PPIs or PDIs between isoforms. The lack of a significant association with PPIs could be due to masking by PPIs which are unrelated to either localization or phase separation, to protein-RNA interactions that are important for phase separation of TFs and were not considered in the analysis, or to a limited number of TF isoforms tested. The lack of a significant association with PDIs could reflect NLS and NES being located across the length of the TF protein, not just in or around the DBD, and to DNA binding not being a major driver of condensate formation.

In total, 19 alternative isoforms (15%) differ in their condensate formation compared to their cognate reference isoforms consistently across cell lines (**Figure S6G**). One example of this is seen for PBX1, a homeodomain TF with known roles in cancer and development.^103,104^ We assayed two isoforms of PBX1, the reference isoform, PBX1a, and an alternative isoform with a truncation and short frame shift at the C-terminus, PBX1b (**Figure 6G-I**). PBX1b has been associated with differentiation,^105–107^ but the molecular mechanism underlying this association is currently unknown. PBX1a forms nuclear condensates consistently in both cell lines, whereas PBX1b does not (**Figures 6J, S6H**). To further characterize the condensate formation in live cells expressing the PBX1 isoforms, we analyzed the relationship between total protein levels (as determined by total cellular GFP signal) compared to the protein level found only in the dilute phase, *i.e.* outside of condensates, across multiple cells displaying a range of overall TF-GFP fusion expression levels.^108^ Proteins that form condensates via phase separation have a critical threshold (the saturation concentration, C_sat_) at which the protein becomes saturated, exhibited by the total concentration of protein being substantially higher than the dilute concentration of protein. C_sat_ analyses confirmed that PBX1a phase separates to form condensates (**Figure 6K**), whereas PBX1b does not. Moreover, the complement of the slope of the line above the C_sat_, the ‘dominance’, is determined by whether the protein is sufficient to phase separate on its own, as indicated by a flat slope, or if it requires other factors, as indicated by a steeper slope.^109^ PBX1a has low dominance (**Figure 6K**), indicating that PBX1a likely phase separates via interactions with other factors rather than on its own. Consistent with this result, in our PPI assay, PBX1a showed interactions with three protein partners that PBX1b does not: LNX1, PIN1, and TMF1 (**Figure 6H**). Of these three differential partners, TMF1 is a co-activator whose known partner TRNP1 regulates nuclear condensates in neural differentiation.^110–112^ Additionally, PBX1b shows substantially reduced transcriptional activity in the M1H assay compared to PBX1a (**Figure 6I**). Together, these results suggest a potential model where PBX1a forms transcriptionally active nuclear condensates via its interactions with protein partners that are not retained by PBX1b.

Overall, TF alternative isoforms were more likely to differ in their cellular localization than in their ability to form condensates (**Figure 6C**). For example, while all isoforms of FOXP2, a forkhead TF that plays important roles in language development,^113^ show a striking ability to form condensates, the localization of these condensates differ (**Figures 6L-N, S6I**). Alternative isoforms of FOXP2 exhibit complex tissue and cell specificity, but their functional significance remains unclear.^114–116^ The reference isoform of FOXP2 contains two NLS regions flanking its DBD.^117^ Alternative isoforms that are missing the NLS form condensates in the cytoplasm, whereas those retaining the NLS form condensates in the nucleus. These results are consistent with a study showing that truncated mouse FOXP2 isoforms missing the NLS are localized to the cytoplasm in Purkinje cell development.^118^ Moreover, whereas FOXP2 isoform localization can be explained by the annotated NLS, in general, we found very few annotated NLSs within our clone collection and, as a result, we find no clear association between localization and the presence of an NLS (**Figure S6J**). This highlights the incompleteness of NLS/NES annotation databases^119^ and the challenge of predicting localization and condensate formation from sequence alone given their dependence on specific PPIs. As the non-overlapping isoforms FOXP2-4 and FOXP2-10 both appear in cellular condensates, this implies that the sequence determinants of condensate formation are not restricted to one protein region, highlighting the challenge of predicting condensate behavior from sequence alone. Altogether, our data reinforce the need for expanded, isoform-aware characterization of TF condensate formation and subcellular localization, which are shaped by a complex network of macromolecular interactions.

### Multi-dimensional characterization of TFs reveals two major classes of alternative isoforms: negative regulators and rewirers

Several well-characterized examples of alternative TF isoforms act as negative regulators of their cognate reference isoforms.^27,36^ For example, the alternative isoform of STAT3, STAT3beta, is missing the C-terminal transactivation domain while retaining the DBD. STAT3beta therefore binds STAT3 targets without activating transcription, thus inhibiting the reference isoform, STAT3alpha, from activating STAT3 GRNs.^37,38^ Indeed, whereas STAT3alpha acts as a canonical oncogene, STAT3beta acts as a tumor suppressor.^120^ However, the extent to which these known examples of negative regulators are representative of the global landscape of TF isoform function is unknown. We therefore sought to use our large-scale dataset of TF isoform properties to address whether putative negative regulators are common among alternative TF isoforms.

We classified the 174 alternative TF isoforms for which we have data in at least two molecular functional assays (PDIs, PPIs, and transcriptional activity) into three categories: negative regulators, rewirers, and those that are similar to their cognate reference isoforms (**Figure 7A, Table S13**). In contrast to the reference, negative regulator alternative isoforms are those expected to negatively affect the function of their cognate reference isoforms. For example, if an alternative isoform fails to bind key cofactors but binds to the same genomic targets, it could prevent the reference isoform from activating target genes (analogous to STAT3beta). We therefore defined’’negative regulators’’ as alternative TF isoforms that completely *lose* function in at least one assay (PDIs, transcriptional activity, or PPIs) while *retaining* function in another (**Figure 7B**). We defined complete loss of function of an alternative isoform compared to its reference as follows: alternative isoforms with (i) 0 PDIs or loss of ≥ 10% of the DBD, (ii) loss of activation or repression (M1H log2FC relative to control between 1 and-1, and ≥ 2-fold difference compared to the reference isoform), or (iii) 0 PPIs or loss of all PPIs of either within-family TFs of obligate dimers, signaling proteins, or transcriptional cofactors. For example, the alternative isoform of CREB1, CREB1-1, fails to bind DNA but retains its ability to strongly activate transcription and thus might interfere with the function of the reference isoform by sequestering key cofactors (**Figure 7E**). We considered any alternative TF isoforms that have identical PDI and PPI profiles and ≤ 2-fold difference in M1H to their cognate reference isoforms to be “similar”, and any alternative isoforms that were otherwise different in PDI, PPI, or M1H profiles (without losing function in ≥ 1 assay) to be “rewirers” (examples shown in **Figure S7A-B**). Only one isoform, an alternative isoform of PPARG, loses function across all tested axes (**Figure S7C**); we considered this isoform “likely non-functional” and filtered it out of downstream analyses. In addition, we considered the subcellular localization, as determined from our high-throughput imaging data, for 129 alternative isoforms in our clone collection. We classified any alternative isoforms whose localization changed from nuclear or both nuclear/cytoplasmic in their cognate reference isoform to exclusively cytoplasmic in either HEK293T or U2OS cells as negative regulators, as they have the potential to sequester the reference isoform in the cytoplasm; isoforms showing any other differences in localization from that of the reference isoform were considered to be rewirers.

**Figure 7:**
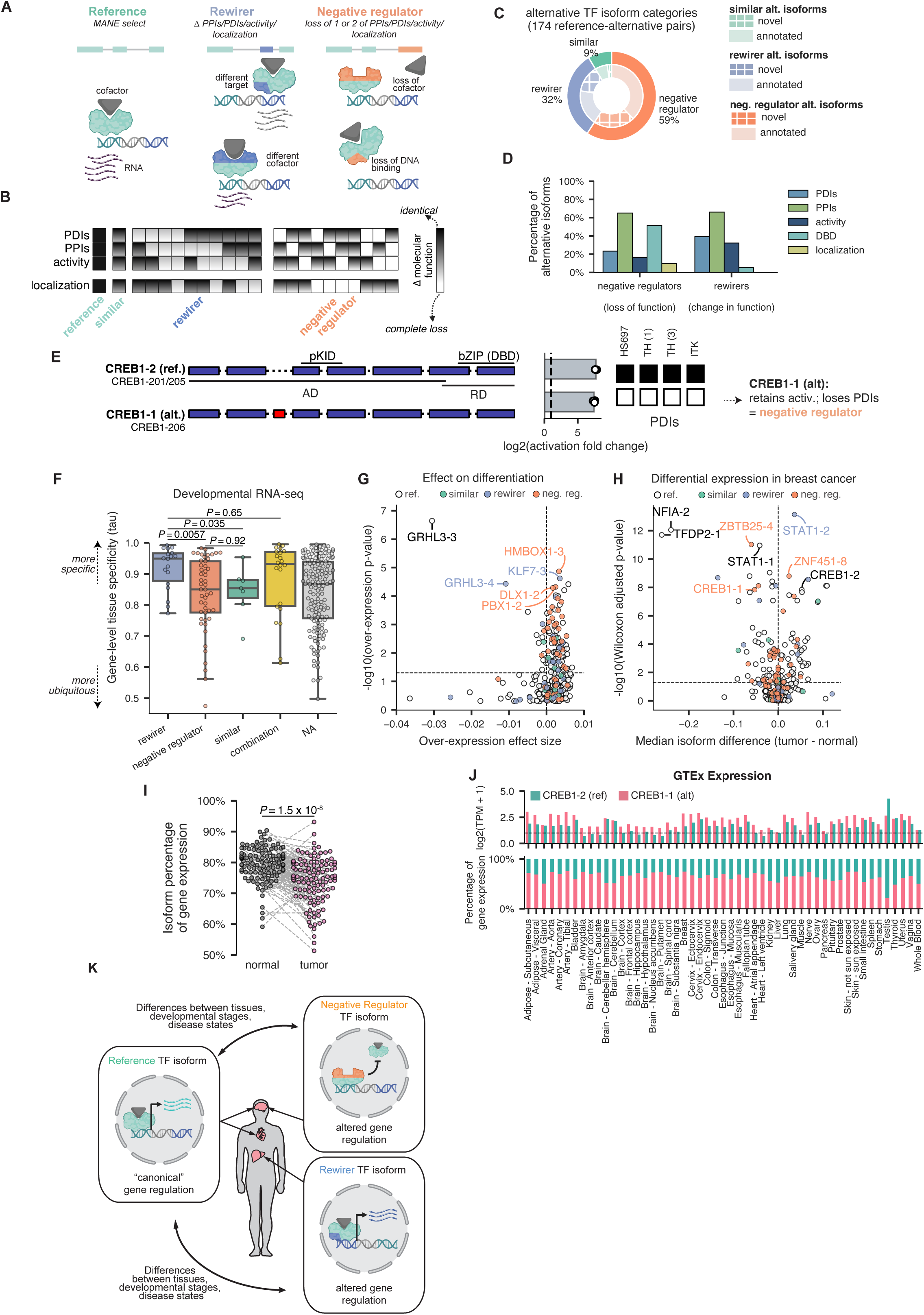
Alternative TF isoforms can function as negative regulators. **A.** Schematic showing examples of TF isoforms classified as either rewirers or negative regulators. **B.** Cartoon heatmap showing how molecular function assay results were used to classify alternative isoforms as either similar to the reference, rewirers, or negative regulators. **C.** Nested pie chart showing the number of alternative isoforms categorized as either similar to reference, rewirers, or negative regulators (outer circle) and the number of annotated (solid colors) and novel (hatched colors) isoforms that comprise each category. **D.** The percent prevalence of various changes in molecular function among rewirers and negative regulators (left graph) compared to the percent prevalence of each assay among all alternative isoforms. Note that because most TFs have been assessed in ≥1 assay, these categories are not mutually exclusive with each other. **D.** The percent of alternative isoforms that either show loss of function in a particular assay (if categorized as a negative regulator, left) or change in function in a particular assay (if categorized as a rewirer, right) as compared to their reference isoforms. Note that because most TFs have been assessed in at least one assay, these categories are not mutually exclusive with each other; for a full plot of negative regulator classification reasons, see **Figure S7D**. **E.** Example of a negative regulator TF isoform (CREB1-alt). Left: exon diagram showing domain annotations. AD = activation domain; RD = repression domain; pKID = phosphorylated kinase-inducible domain. Middle: M1H results. Right: PDIs. **F.** Boxplot showing the gene-level tissue specificities (tau metric)^143^, calculated from the Developmental RNA-seq data^54^, among TF genes with either only rewirer alternative isoforms, only negative regulator alternative isoforms, only alternative isoforms that are similar to reference, some combination of the above, or only alternative isoforms that were unable to be classified (NA). P-values shown are from a two-sided Mann Whitney test. **G.** Volcano plot showing the effect of TF over-expression on differentiation. The over-expression effect size (x-axis) is the Diffusion difference and the p-value (y-axis) is the-log10 of the Diffusion P-value, both as calculated in the TF mORF Atlas.^61^ **H.** Volcano plot showing differential abundance of TF isoforms in breast cancer. The median isoform difference (x-axis) is the median difference of fractional isoform expression among paired tumor/normal breast cancer samples and the p-value (y-axis) is the-log10 of the adjusted, paired Wilcoxon p-value. **I.** Paired swarm plot showing the relative expression of the alternative isoform of CREB1 (as a fraction of total CREB1 gene expression) in matched breast cancer tumor and normal samples (from the same patient). P-value shown is from a two-sided Mann Whitney test, adjusted for multiple hypothesis correction. **J.** Expression levels of CREB1 isoforms in GTEx. Top: log2 TPM values for each CREB1 isoform; bottom: isoform expression as a percentage of total gene expression for each CREB1 isoform. **K.** Schematic model showing one example mechanism of how, whereas rewirer isoforms lead to altered GRNs, negative regulator isoforms can lead to misregulation of canonical GRNs either in the absence or presence of the reference isoform. Negative regulator TF isoforms that outcompete their reference isoforms in the same cell can be thought of as naturally-occurring dominant negatives.

Of the classified alternative isoforms, 103 (59%) were negative regulators, 56 (32%) were rewirers, and 15 (9%) were similar to their cognate reference isoforms. Thus, the vast majority (91%) of alternative isoforms differed substantially from their cognate reference isoforms in at least 1 molecular property. Novel isoforms and annotated isoforms were distributed equally among both negative regulators and rewirers. Negative regulators were found to display loss of any of the three major TF functions: DNA binding, PPIs, or transcriptional activity (**Figure 7D, Figure S7D**), or by loss of subcellular localization, and were found across all major TF families (**Figure S7E**). Intriguingly, TF genes with only negative regulator alternative isoforms tended to be more ubiquitously expressed than TFs with only rewirer alternative isoforms, in both the Developmental RNA-seq (**Figure 7F**) and GTEx data (**Figure S7F)**. Overall, our functional assay results revealed that alternative TF isoforms that can act as negative regulators are widespread among TFs. This suggests that TF negative regulators might commonly serve as an additional layer of regulation through which target gene expression levels are regulated across cell states and tissues.

### Alternative TF isoforms are associated with differentiation and cancer

TFs play important roles in early development. Recently, Joung *et al.* performed a scRNA-seq over-expression screen of > 3,000 annotated TF isoforms in human embryonic stem cells and found many TF isoforms that significantly affected differentiation.^61^ We therefore sought to intersect the results of their screen with our functional data to determine whether there is a differential impact of negative regulators and rewirers on differentiation. 220/246 (89%) of our reference TF isoforms and 183/446 (41%) of our alternative TF isoforms were included in their clone library. As expected, because Joung *et al.* relied on gene annotations to create their library, the majority of our alternative isoforms that were missing in their library are our novel isoforms (167/263 (63%)). Reference isoforms and both rewirer and negative regulator alternative TF isoforms all significantly affected differentiation in the over-expression screen, but the TF isoforms with the strongest effect sizes tended to be reference or rewirer isoforms (**Figure 7G**). The TF that showed the strongest overall effect on differentiation is the reference isoform of GRHL3, a known regulator of many stages of embryogenesis, including neural tube closure and craniofacial development.^121^ GRHL3 has three annotated alternative isoforms: two negative regulators, which lose their ability to bind to (and thus heterodimerize with) GRHL2, and one rewirer (**Figure S7G**). In addition to the reference isoform, only the rewirer isoform of GRHL3, GRHL3-203, drove differentiation when over-expressed, suggesting that the ability of GRHL3 isoforms to dimerize is key to their biological function (**Figure S7G**). This is consistent with missense mutations in the dimerization domain of *grhl3* (homologous to the human reference GRHL3 isoform) resulting in embryonic development problems in zebrafish.^122^ Interestingly, only one of the two negative regulator isoforms that failed to interact with GRHL2 is missing the C-terminal, annotated dimerization domain,^123^ further highlighting the importance of functional assays to characterize TF isoforms rather than relying on domain-based computational predictions of protein function.

Since many of the most well-characterized TF isoforms are negative regulators that are dysregulated in cancer (*e.g.*, STAT3beta),^36^ we next examined the expression of our TF isoform collection in The Cancer Genome Atlas (TCGA). We focused on breast cancer since it has the highest number of clinical samples in TCGA, including 112 paired tumor/normal samples from the same patients (**Table S14, STAR Methods)**. We found that 191 TF isoforms in our clone collection, many of which were reference TF isoforms (78/191 (41%)), showed significant differential abundance between paired tumor and normal samples (adjusted p-value < 0.05, two-sided paired Wilcoxon test) (**Figure 7H, Table S15**). Interestingly, however, several negative regulator isoforms were among the most differentially expressed TF isoforms, including isoforms of ZBTB25, ZNF451, and CREB1. In total, 34/114 (30%) of the alternative TF isoforms that showed significant differential abundance in breast cancer patients were classified as negative regulators, while in contrast only 10/114 (9%) were classified as rewirers, suggesting that misregulation of negative regulator TF isoforms plays specific roles in rewiring GRNs in the context of cancer.

The alternative isoform of CREB1, CREB1-1, is an example of a negative regulator that is significantly misregulated in breast cancer. CREB1 (cyclic AMP response element-binding protein) is a bZIP TF that plays important roles in a number of developmental processes, including cell cycle progression, DNA repair, and differentiation.^124^ CREB1 is considered an oncogene in several cancer types; indeed, small molecule CREB inhibitors have shown therapeutic promise in preclinical studies of leukemia, breast cancer, and glioma.^125^ The alternative isoform of CREB1 differs from the reference isoform by the inclusion of a small, in-frame, 14-a.a. exon in an unstructured region of the protein (**Figure 7E**, **Figure 3C**). This alternative isoform retained its ability to strongly activate transcription but lost all PDIs and was therefore classified as a negative regulator (**Figure 7E**). Intriguingly, while the overall levels of *CREB1* gene expression were similar in breast tumors compared to matched normal controls (**Figure S7H-I**), there was a difference in relative isoform abundance: the alternative isoform was significantly down-regulated in tumors compared to the reference isoform (**Figure 7I, Figure S7J**). Moreover, *CREB1* was ubiquitously expressed, with both the reference and alternative isoforms being expressed in almost all healthy tissues (**Figure 7J**). Thus, our results are consistent with a model wherein the reference isoform of CREB1 acts as an oncogene, but the alternative isoform of CREB1–which we find to be a negative regulator–may have tumor suppressive properties. The fact that these two isoforms are co-expressed in the same tissues suggests that the alternative isoform of CREB1 may act as a dominant negative regulator of the reference isoform.

Taken together, our comprehensive, multi-dimensional characterization of hundreds of TF isoforms reveals that TF gene loci encode alternative proteoforms that fall into two primary categories: rewirers, which behave distinctly from their cognate reference isoforms, and negative regulators, which have the capacity to act either independently or in competition with their cognate reference isoforms, depending on their expression profiles (**Figure 7K**).

## Discussion

Transcriptomic analyses have revealed that TFs are commonly expressed as a series of isoforms generated by alternative promoters, splicing, and polyadenylation sites. However, the extent to which TF isoforms differ across key molecular functions and properties has remained unclear. To understand the functional differences between TF isoforms, we generated a collection of 693 TF isoform ORFs across 246 TF genes and systematically assayed their DNA binding, protein-protein interactions, transcriptional activation/repression, localization, and condensate formation. We provide the integrated results of our multi-dimensional profiling of TF isoforms at the website tfisodb.org as a resource to the community.

We additionally integrated these functional assays with isoform-aware expression analyses across two main data sets: GTEx^53^ and a time course of human development.^54^ We found that most alternative TF isoforms showed a more subtle ‘shift’ in their expression between tissues and developmental time points, rather than a dramatic ‘switch’ (**Figure 1G**). However, these observations are limited by the available data. Additional long-read sequencing of individual cell types, multiple conditions, and specific developmental stages may identify specific contexts in which these alternative isoforms are highly expressed. Moreover, we note that in this study, we tested different protein isoforms, but differences in their UTRs can also affect isoform mRNA stability and protein expression.^126^

Other clone collections of TF isoforms^60,61^ have been generated using gene synthesis. Such approaches are entirely reliant on gene annotation datasets. Here, we instead used a PCR-based approach to generate our clone collection.^127^ This PCR-based approach has two major advantages: (1) it is more cost-effective, and (2) it is less reliant on annotation. As a result, our TF isoform clone collection includes 183 high-confidence novel alternative TF isoforms (26% of the total library). These novel TF isoforms are expressed at similar maximum levels and behave similarly in our assays compared to annotated alternative TF isoforms (**Figure 2C, 2F**).

Overall, we found that two-thirds of alternative TF isoforms differ from their cognate reference isoforms in at least one molecular function. This is likely an underestimate of the real differences between isoforms, due to limitations of our assays. For example, further differences in TF-DNA binding may be revealed by testing a larger set of DNA baits in the eY1H assays. Additionally, there may be context-specific differences in TF-protein interactions (*e.g.*, those that depend on specific post-translational modifications) that were missed in the Y2H assays or cell-type-specific differences in transcriptional activity that might be uncovered by performing M1H assays in additional cell lines or stimulation conditions. One aspect we did not investigate was that, in the case of coexpressed reference and alternative isoforms of a dimerizing TF, there are up to three potential dimers–reference-reference, alternative-alternative and reference-alternative–and these three might each have different DNA-binding and activation properties. For example a prior study has shown that murine Tbx5e^81^ (TBX5-2) can heterodimerize with Tbx5a (TBX5-1).^81^ Finally, we note that there remain other molecular functions of TFs–such as ligand binding and RNA binding–that are key to their roles in GRNs and that we have not explored here. The exogenous assays we used provide an approach to achieve isoform-level resolution of some of the most important molecular TF functions in high throughput. Improved approaches for assaying isoform-specific TF occupancies *in vivo* and proteome-wide techniques for assaying PPIs, such as proximity labeling followed by mass spectrometry, are needed to achieve isoform-level resolution in endogenous contexts at scale.

The different molecular functions of TF isoforms can be exceedingly difficult to predict from sequence and predicted structure alone. Indeed, we find that differences in regions far from the annotated DBD, often in IDRs, can affect TF-DNA interactions (**Figure 3A**). This is consistent with previous work showing that differences in IDRs can affect TF binding sites *in vivo*,^65,83^ and highlights the importance of studying full-length TFs rather than solely focusing on extended DBDs when assaying DNA binding *in vitro*. While prior *in vitro* studies using short naked oligonucleotides found that full-length TFs and extended DBD constructs typically recognize the same motifs,^3,45^ by performing eY1H assays using full-length TFs, we were able to determine the differences in DNA binding between isoforms in a chromatinized setting in high throughput. Furthermore, combining PBMs of full-length TFs with an updated analysis pipeline that allows for statistical inference of differential affinity across all 8-mers,^78^ allowed us to identify subtle differences in DNA binding affinity and specificity between isoforms of TF genes that differ in sequence regions outside of their DBDs. We also found that differences in transcriptional activation/repression are common across TF isoforms and were often not associated with changes in annotated effector domains (**Figure 4A**), underscoring the continued importance of performing experiments using full-length proteins to characterize the functions of TFs, complementing tiling-based peptide assays.^6,7^

Many of the most well-studied alternative TF isoforms are known to function as natural dominant negative regulators of their cognate reference isoforms.^27,36^ However, the extent to which natural dominant negative isoforms exist within the context of the global “TFome” has remained unclear. Here, we present evidence that negative regulator isoforms (*i.e.*, isoforms that lose at least one molecular function compared to their cognate reference isoform) are likely widespread among TFs and often misregulated in cancer (**Figure 7C, 7H**). Given that ubiquitously expressed TFs tend to have negative regulator isoforms (**Figure 7F**), we propose that in most cases negative regulator isoforms will exert dominant negative effects through diverse mechanisms of action that interfere with reference isoform function. Future studies focused on characterizing how negative regulator isoforms compete with their cognate reference isoforms for key molecular interactions (e.g. PDIs, PPIs) within the cell to affect downstream GRNs are needed to determine whether these negative regulators act as true dominant negatives. However, our findings are consistent with decades-old ideas that negative regulators–whether they arise from genetic mutations^128^ or aberrant splicing^129^–contribute to human disease and highlight the varied ways in which changes in molecular functions can result in negative regulator TF isoforms.

The majority of human TFs have undergone gene duplication and diverged throughout evolution, resulting in large families of TFs with highly similar DBDs.^20^ Paralogous TFs have been studied widely for several decades;^20^ they are known to have differential *in vivo* binding^130^ and expand GRNs.^131,132^ TF paralogs have been associated with organismal complexity and the emergence of novel cell types.^133^ Alternative isoforms increase the diversity of TF proteins as well–albeit through a different mechanism than gene duplication–but their effects on GRNs are less well understood. In this study, we reveal that TF isoforms can behave as distinctly as paralogous TFs across all major molecular functions (TF-DNA binding, TF-protein binding, and transcriptional activity) (**Figures 6F-H**). We propose that alternative isoforms of TFs may be more likely to act as negative regulators than paralogous TFs. Consistent with this idea, the two thyroid receptor paralogs THRA and THRB have each retained their ability to bind to thyroid hormone, but the alternative isoform of THRA does not bind to thyroid hormone and retains its ability to dimerize, thus acting as a dominant negative.^134^ Thus, TF isoforms should be considered for their potential to expand GRN complexity alongside TF paralogs.

In summary, our high-throughput exogenous assays shed light on the functional diversity that naturally exists within the human TFome and is encoded by alternative isoforms. A major current challenge in clinical genomics is to classify so-called variants of unknown significance (VUS), with most efforts–both experimental and computational variant effect predictors (VEP)–focused on either single amino acid changes^135,136^ or noncoding variants.^137,138^ However, splicing variants comprise a neglected but substantial portion (approximately 5%) of VUS, totalling > 11,000 VUS in ClinVar.^139^ Systematic experimental molecular function profiling of full-length isoforms, as performed in this study, could fill the lack of training data for VEP of splicing variants to address this need. Additionally, such isoform-level assessments of protein function may augment emerging methods that model disease associations with isoform-level RNA-seq expression.^140^ Finally, our approach has significant potential when applied to cancer, in which splicing is often dysregulated^141^ and TFs play key roles.^142^ Altogether, our work highlights the importance of moving beyond gene-level resolution and towards a more complex, proteoform-aware characterization of TF function.

## STAR Methods

STAR methods are in the supplementary file.

## Acknowledgements

We thank Norman Davey (Institute of Cancer Research, UK) for sharing his expertise on predicting disordered regions using AlphaFold and Yves Janin (Institut Pasteur, France) for providing the furimazine substrate of the NanoLuc for the mN2H assay. We thank members of CCSB, Bulyk lab and Fuxman Bass lab for helpful discussions. Schematic figures in this paper were created with BioRender.com. The results presented here are in part based upon data generated by the TCGA Research Network: https://www.cancer.gov/tcga.

This work was supported by NIH grant U01CA232161 (M.V., M.L.B., and J.I.F.B.) with additional funding from NIH grants U24HG011451 (M.V., D.E.H., and A.F.), R35GM128625 (J.I.F.B.), R01CA226802 (N.S.), R35GM142647 (G.S.), and R35GM137836 (N. Sahni). Additional support included an NIH Ruth L. Kirschstein Award F32HG012318-01 (K.M.), NIH Ruth L. Kirschstein training grant T32CA009361 (G.S., awardee, N.S. Gray, P.I.), a Charles A. King Trust Postdoctoral Research Fellowship (G.S.), a Melanoma Research Foundation Career Development Award (G.S.), a Belgian American Educational Foundation doctoral research fellowship (F.L.), a Wallonia-Brussels International (WBI)-World Excellence fellowship (F.L.), a Fonds de la Recherche Scientifique (FRS-FNRS)-Télévie grant FC31747 (Crédit n° 7459421F) (F.L., awarded to J.-C.T.), the Fondation Léon Fredericq (F.L. and J.-C.T.), a University of Liège mobility grant (F.L.), a Fonds de la Recherche Scientifique (FRS-FNRS) Mobility and Congress funding (n°40020393) (F.L.), a Josée and Jean Schmets Prize (F.L.), a Herman-van Beneden Prize (F.L.), a Canadian Institutes for Health Research Foundation grant FDN-159926 (M.G., awarded to F. P. Roth), Deborah F. Allinger Fellowship awarded to CCSB (L.L.). N. Sahni is a CPRIT Scholar in Cancer Research with funding from the Cancer Prevention and Research Institute of Texas (CPRIT) New Investigator Grant RR160021 and supported by the Andrew Sabin Family Foundation Fellowship. J.A.R. is a CPRIT Scholar in Cancer Research with funding from the Cancer Prevention and Research Institute of Texas (CPRIT) New Investigator Grant RR210040. Laser scanning confocal microscopy and image processing were performed at the MD Anderson Cancer Center Epigenetics & Molecular Carcinogenesis Flow Cytometry and Cell Imaging Core (EMC-FCCIC) and Advanced Microscopy Core (AMC) with funding support provided by the CPRIT core facility grant RP170628. M.V. is a Chercheur Qualifié Honoraire and J.-C.T. is a Maître de Recherche from the Fonds de la Recherche Scientifique (FRS-FNRS, Wallonia-Brussels Federation, Belgium). M.L.B. is a co-inventor on U.S. patents # 6,548,021 and # 8,530,638 on PBM technology and corresponding universal sequence designs, respectively. Universal PBM array designs used in this study are available via a Materials Transfer Agreement with The Brigham & Women’s Hospital, Inc. No other competing interests to report.

---

**Figure S1:**
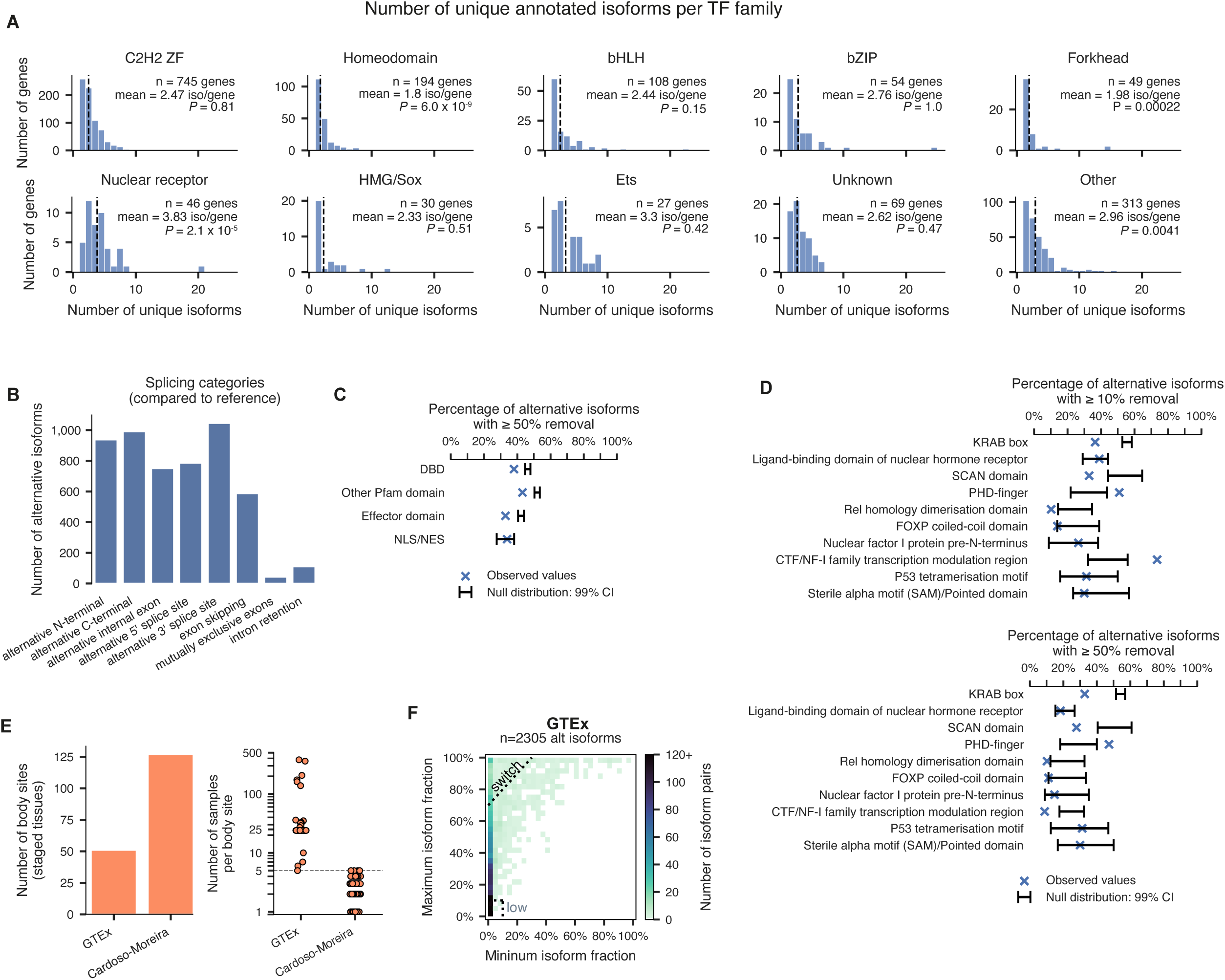
Sequence and expression diversity of annotated TF isoforms, related to Figure 1. **A.** Number of unique annotated protein isoforms per TF family. Mean number of isoforms per gene is shown as a dotted vertical line. Only TF families with ≥ 20 genes are shown; the remaining TF families are collapsed into the “other” category. **B.** Number of alternative isoforms that exhibit various sequence differences compared to their cognate reference isoforms. Categories are not mutually exclusive (so an alternative isoform could exhibit both an alternative N-terminal and exon skipping, for example). **C.** Barplot showing the observed fraction of alternative isoforms with ≥ 50% removal of various protein domains (green bars) compared to the expected fraction (black error bars, 99% CI) as defined by a null model assuming the domain is randomly positioned along the protein. DBD = DNA-binding domain; NLS/NES = nuclear localization/export signal. **D.** Analogous to **C**, but showing specific domains that are collapsed in the “Other Pfam domains” category in **C**. Only domains with ≥30 annotation instances are shown. **E.** Number of unique body sites (i.e., staged tissues) (left) and number of samples per body site (right) for both GTEx and developmental RNA-seq from Cardoso-Moreira et al.^54^ Cardoso-Moreira has more unique body sites, but fewer individual samples per body site, compared to GTEx. **F.** Heatmap showing the maximum isoform fraction (y-axis) compared to the minimum isoform fraction (x-axis) of alternative TF isoforms in re-sampled GTEx, where isoform fraction is defined as the expression level of an isoform normalized to the total expression level of its host gene. Dashed lines show the definitions used for isoforms that exhibit “switching” events and isoforms that remain lowly expressed. Only isoforms whose host genes are expressed at ≥ 1 TPM in ≥ 1 sample are shown.

**Figure S2:**
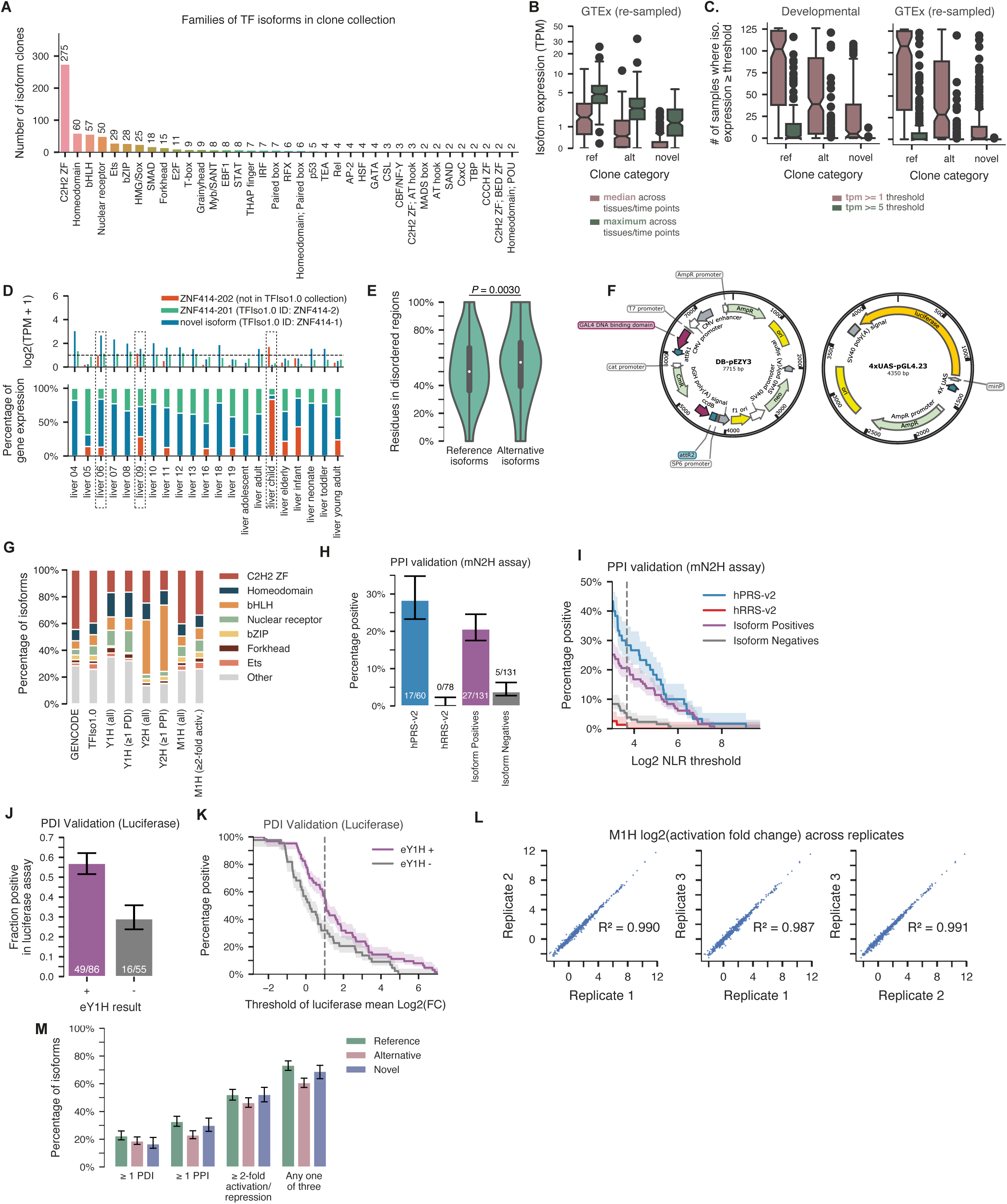
Overview of TFIso1.0 clone collection and TF molecular function assays, related to Figure 2. **A.** Histogram showing count of clones in TFIso1.0 across all observed TF families. **B.** Boxplot showing the median and maximum expression levels (in TPM) in re-sampled GTEx RNA-seq data of reference, annotated alternative, and novel alternative isoforms in TFIso1.0. **C.** Box plots showing the distribution of the number of samples where reference, alternative, or novel TF isoforms are expressed ≥ 1 TPM or ≥ 5 TPM in developmental RNA-seq and re-sampled GTEx. **D.** Example expression profile of a novel isoform in TFIso1.0, ZNF414-1. Top: log2 TPM values for each ZNF414 isoform; bottom: isoform expression as a percentage of total gene expression for each ZNF414 isoform. All liver samples from developmental RNA-seq data are shown. Samples where ZNF414-1 is expressed ≥ 1 TPM are outlined. **E.** Violin plot showing the fraction of residues predicted to be in disordered regions, per isoform, comparing reference and alternative isoforms. White dot indicates the median, dark-gray box indicates IQR. P-value calculated using a two-sided permutation test. **F.** Plasmids used in the M1H assay. **G.** Stacked barplot showing the percent of TF isoforms belonging to various TF families in GENCODE, the entire TFIso1.0 collection, those that have been successfully tested in each assay (“all” categories), and those that show evidence of function (≥ 1 PDI, ≥ 1 PPI, ≥ 2-fold M1H activity) in each assay. **H, I.** Results of testing our Y2H PPI data in the mN2H assay, along with positive and negative controls, displayed as a bar chart (**G**) and a titration across the readout value (**H**), with the cutoff displayed as a vertical dashed line. Error bars/bands are 68.3% Bayesian CI. PRS = positive reference set; RRS = random reference set; Lit-BM = Literature curated PPIs with binary and multiple evidence. **J, K.** Results of testing our Y1H PDI data in the luciferase assay, displayed as a bar chart (**I**) and a titration across the readout value (**J**). Error bars/bands are 68.3% Bayesian CI. **L.** Scatter plots showing the correlation across 3 independent transfection replicates of the mammalian one-hybrid experiment. **M.** Barplot showing the proportion of isoforms exhibiting ≥ 1 PPI, ≥ 1 PDI, ≥ 2-fold activation/repression in M1H, or any one of the three across reference, annotated alternative, and novel alternative isoforms, normalized to the total number of isoforms in TFIso1.0. Error bars are 68.3% Bayesian CI.

**Figure S3:**
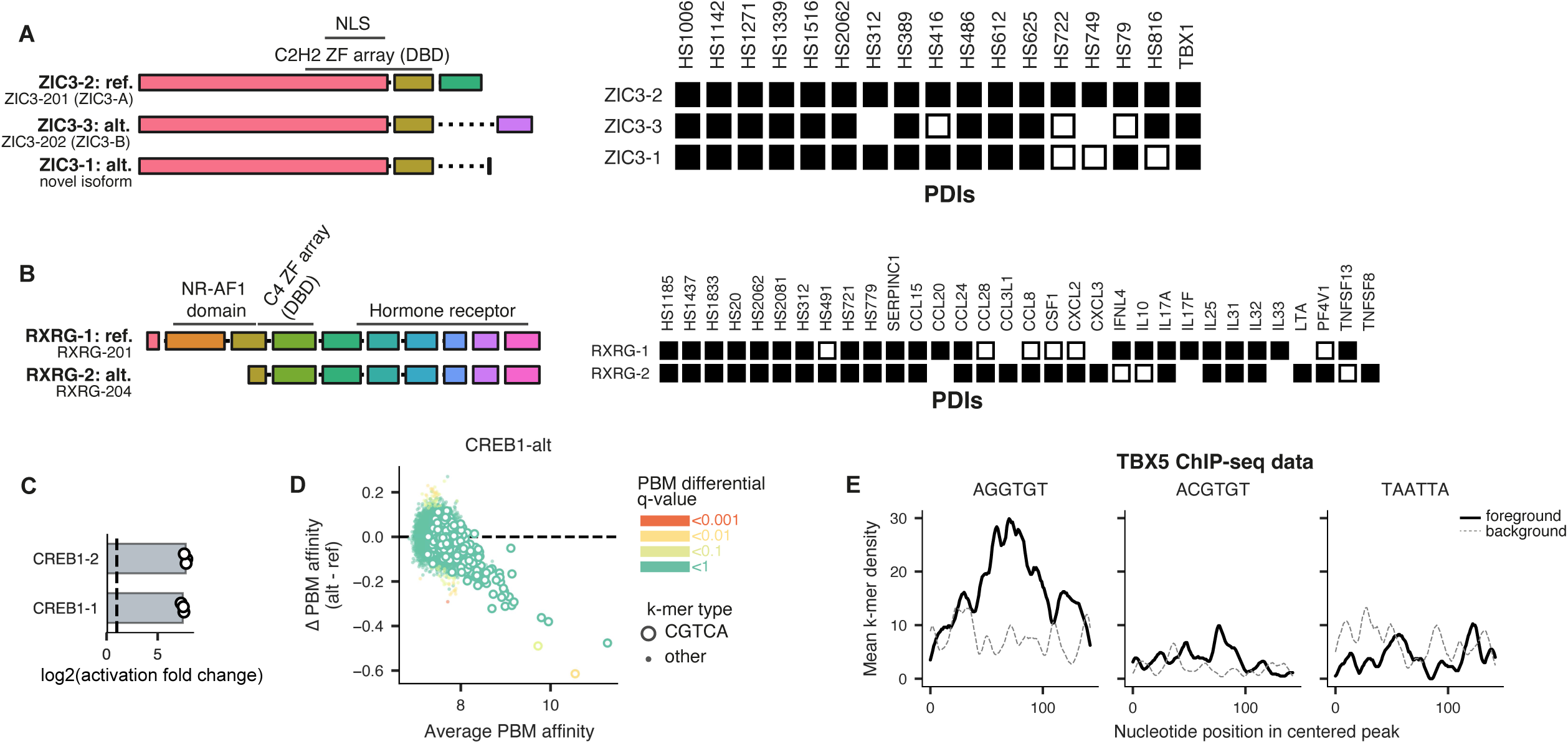
DNA binding preferences of TF isoforms, related to Figure 3. **A.** Left: exon diagrams of the 3 ZIC3 isoforms included in TFIso1.0. NLS = nuclear localization sequence. Right: PDI results from the Y1H assay for the 3 isoforms of ZIC3. Missing boxes correspond to baits that were not successfully tested against ZIC3-3. **B.** Left: exon diagrams of the 2 RXRG isoforms. Right: PDI results from the Y1H assay for the 2 isoforms of RXRG. Missing boxes correspond to baits that were not successfully tested against one of the isoforms. **C.** Mammalian one-hybrid (M1H) activity results for CREB1 isoforms. Both isoforms have high activation capacities. **D.** MA scatter plot showing the PBM results comparing the alternative and reference isoforms of CREB1 for every 8-mer. Points are colored by the differential affinity q-value calculated by the upbm package.^78^ Open circles correspond to 8-mers containing the canonical CREB1 5-mer CGTCA (or its reverse complement). Points below the dashed horizontal line correspond to 8-mers for which the alternative isoform shows reduced affinity compared to the reference isoform. **E.** Enrichment of the canonical TBX5 6-mer AGGTGT, the altered TBX5 6-mer ACGTGT, or a negative control Homeodomain 6-mer TAATTA (or each of their reverse complements) across TBX5 ChIP-seq peaks. Peaks were centered to the nucleotide corresponding to the highest ChIP enrichment over background and trimmed to 150 nucleotides. Solid black lines show enrichment in ChIP peaks (foreground); dotted grey lines show enrichment in matched genomic negative control regions (background). Lines show the moving average of k-mer density, using a window of 8 nucleotides.

**Figure S4:**
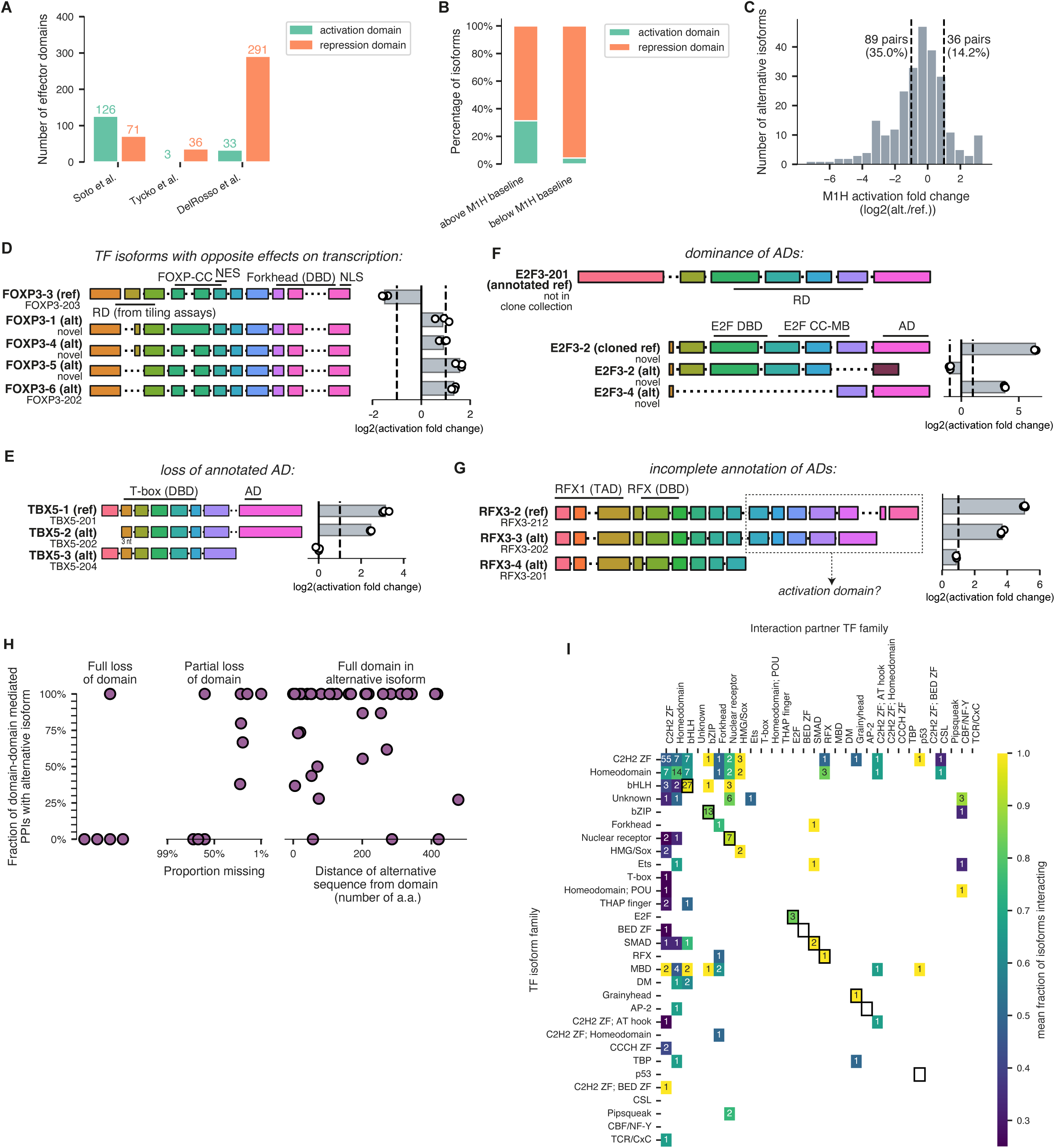
Transcriptional activity and protein binding preferences of TF isoforms, related to Figure 4. **A.** Bar plot showing the number of effector domains, broken up into activation and repression, annotated in each of the 3 studies used in this work. Note that Soto et al.^8^ is based primarily on literature curation, whereas Tycko et al.^6^ and DelRosso et al.^7^ are each large-scale tiling screens. **B.** Bar plot showing the percent of TF isoforms containing an either annotated activation or repression domain that are either above the mammalian one-hybrid (M1H) baseline activity levels (≥1) or below baseline activity levels (≤-1). **C.** Histogram showing the distribution of M1H activity changes (log2(alternate isoform M1H activity/reference isoform M1H activity)) across all pairs assayed. **D-G.** Example of TF genes with isoforms that have opposite effects on transcription (**D**), lose an annotated activation domain (**E**), show dominance of annotated activation domains over repression domains (**F**), and show potentially incomplete effector domain annotation (**G**). Left: exon diagrams of FOXP3, TBX5, E2F3, and RFX3 isoforms, respectively. Right: transcriptional activity from the M1H assay for the denoted isoforms. **H.** The fraction of the subset of PPIs mapped to domain-domain interactions that are retained in each alternative isoform, relative to the reference isoform, in cases where the alternative isoform fully or partially loses the interacting domain, or contains the full domain. **I.** Full heatmap showing the rewiring score for combinations of families of TF isoforms (y-axis) and families of TF PPI partners (x-axis). Within-family dimerizations are therefore denoted on the diagonal of the heatmap. TF families that bind DNA as obligate dimers are marked with outlined black boxes on the diagonal. The number within each cell indicates the number of PPIs that fall into that specific category, and the color denotes the rewiring score.

**Figure S5:**
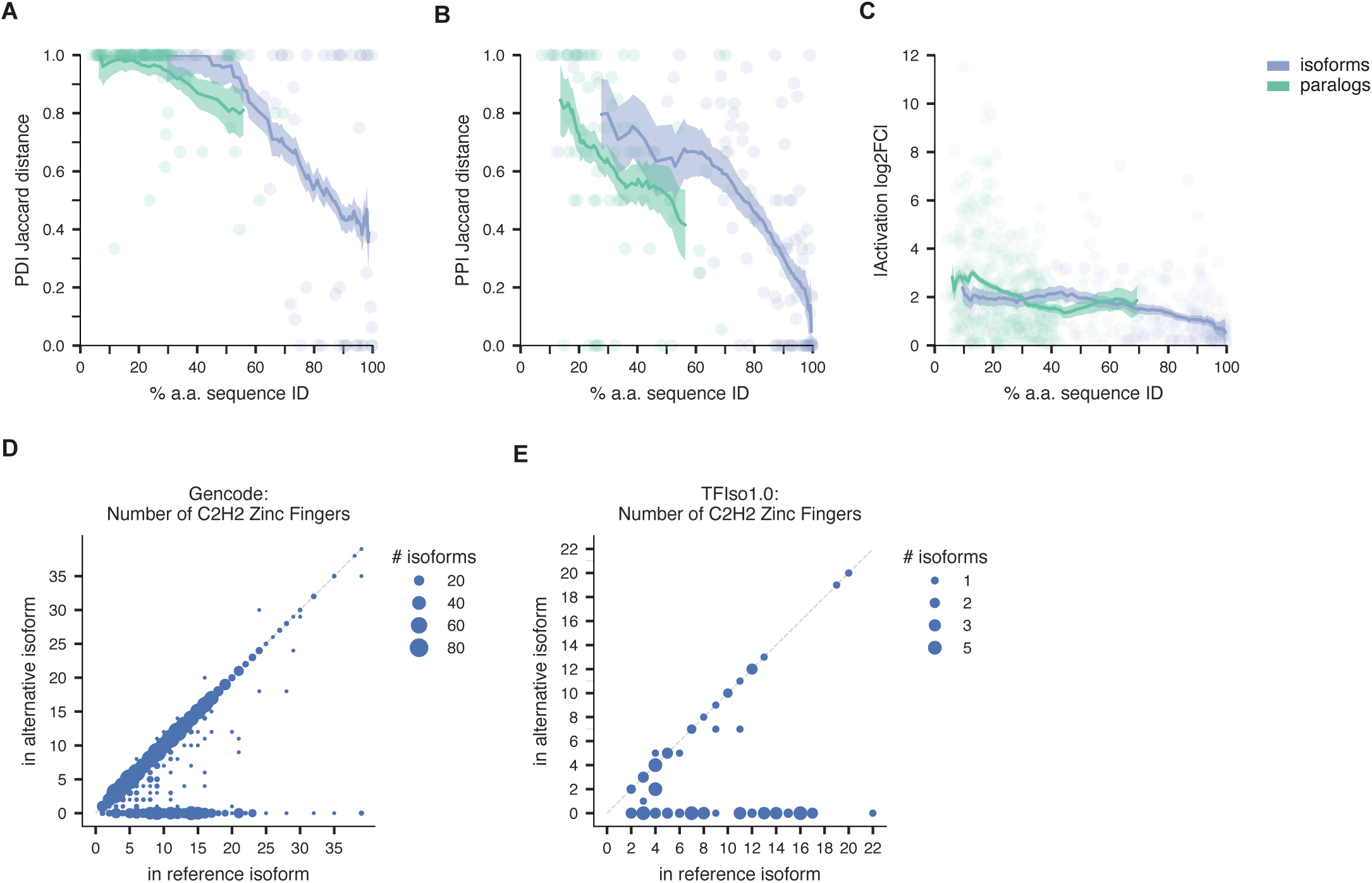
Functional differences between TF isoforms and TF paralogs, related to Figure 6. **A-C.** Scatter plots showing the Jaccard distance in PDIs (**A**) or PPIs (**B**) or the absolute log2 fold-change in M1H activity (**C**) between pairs of isoforms (blue) or paralogs (green) (y-axis) as compared to their pairwise amino acid sequence similarity (x-axis). Lines show mean values across a sliding window of 40%; error bands are 68.3% Bayesian CI; P-values are calculated using a two-sided permutation test. **D-E.** Bubble plots showing the number of zinc fingers in annotated zinc finger array TFs in either the reference isoform (x-axis) or alternative isoform (y-axis); size of the circles corresponds to the number of isoform pairs in each bin. **D**: considering all isoforms annotated in GENCODE; **E**: considering only isoforms in TFIso1.0.

**Figure S6:**
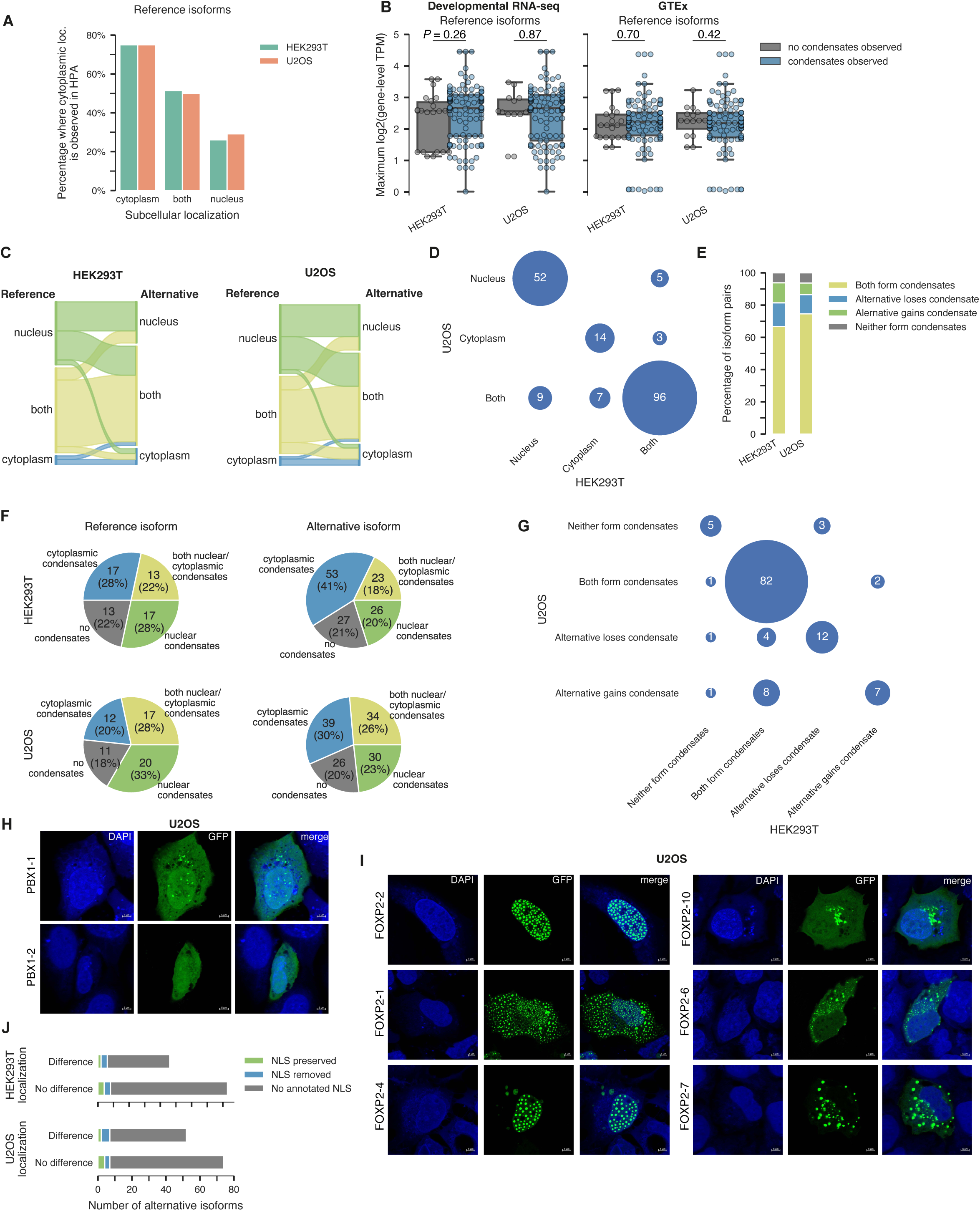
Condensate formation and subcellular localization differences between TF isoforms, related to Figure 6. **A.** Sankey plots showing how the localization of alternative isoforms change as compared to their cognate reference isoforms in HEK293T (left) and U2OS (right) cells. **B.** Heatmap showing the agreement in localization calls among all TF isoforms in HEK293T and U2OS cells. Size of the circle is proportional to the number of TF isoforms in that bin (shown in white). **C.** Stacked bar plot showing the percent of reference-alternative isoform pairs where both show condensates, the alternative gains or loses condensates compared to the reference, or neither isoform shows condensates in either HEK293T or U2OS cells. **D.** Pie charts showing the distribution of condensate localization among reference and alternative isoforms in HEK293T and U2OS cells. **E.** Heatmap showing the agreement in condensate call differences among reference-alternative TF isoform pairs in HEK293T and U2OS cells. Size of the circle is proportional to the number of TF isoforms in that bin (shown in white). **F.** Bar plot showing the percentage of reference isoforms that show cytoplasmic localization in the Human Protein Atlas (y-axis, **STAR Methods**) compared to their localization in our high-throughput imaging assay (x-axis) in either HEK293T or U2OS cells. **G.** Box plots showing the maximum expression of reference isoforms (in TPM, y-axis) in either Developmental RNA-seq (left) or GTEx (right) broken up by whether or not the reference isoform forms condensates in our high-throughput imaging assay (color) in either HEK293T or U2OS cells. P-values shown are from a two-sided Mann Whitney test. **H.** Representative images of PBX1 isoform expression in U2OS cells (63x magnification). **I.** Representative images of FOXP2 isoform expression in U2OS cells (63x magnification). **J.** Stacked bar plots showing the number of alternative isoforms with NLS preserved or lost, relative to the reference isoform, split by whether there was an observed difference in localization between the reference and alternative isoform in HEK293T (top) and U2OS cells (bottom).

**Figure S7:**
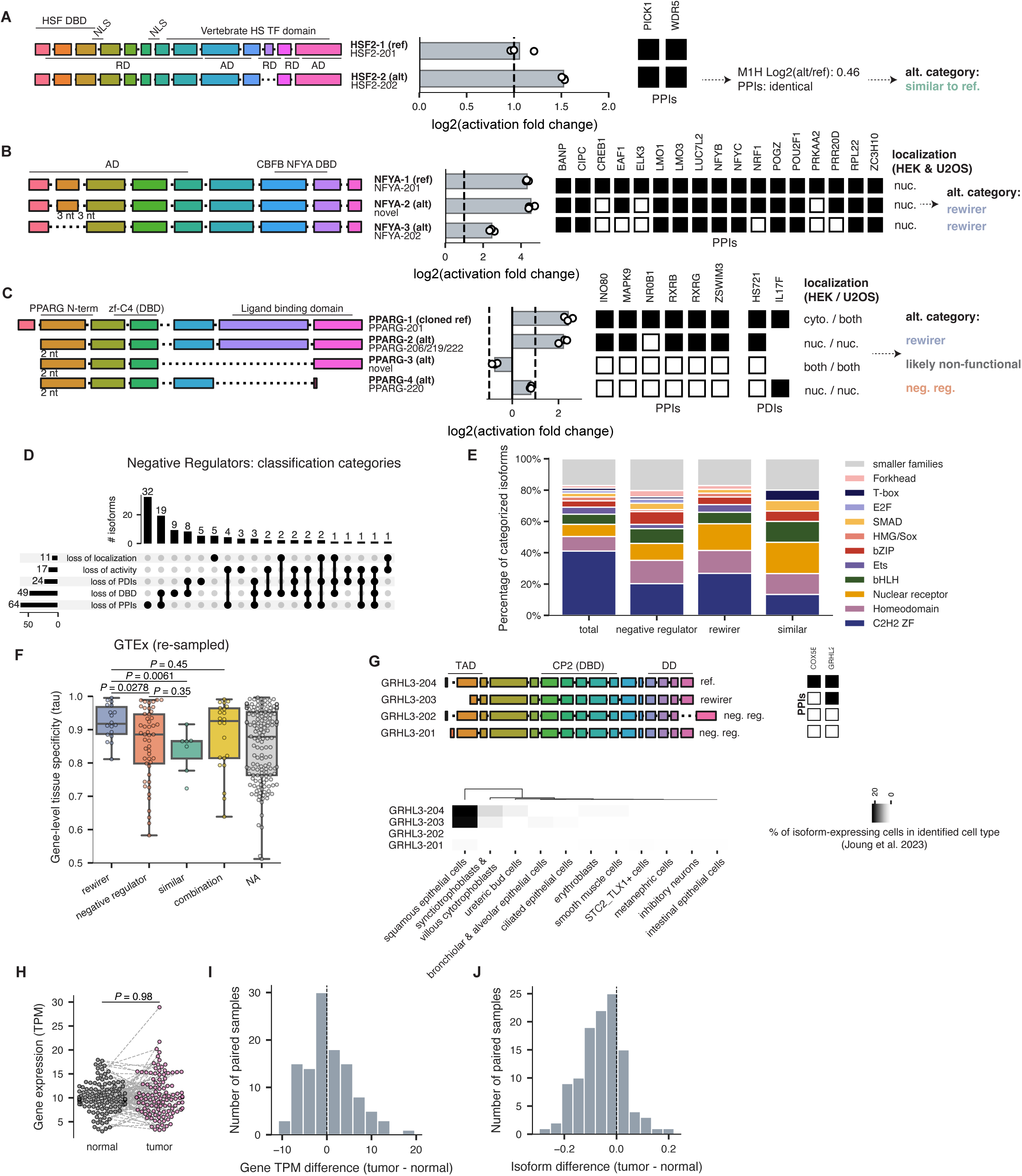
Alternative TF isoforms can function as negative regulators, related to Figure 7. **A-C.** Examples of TF genes with isoforms that are similar to the reference (A), rewirers (B, C), negative regulators (C), and likely non-functional (C). For each gene, all assays (Y1H, Y2H, M1H, localization) with data are shown. **D.** UpSet plot showing the reasons why negative regulator TF isoforms were classified as such, depending on their assay results. **E.** Stacked bar plot showing the distribution of TF families among each category of alternative isoform. **F.** Boxplot showing the gene-level tissue specificities (tau metric)^143^, calculated from the GTEx RNA-seq data, among TF genes with either only rewirer alternative isoforms, only negative regulator alternative isoforms, only alternative isoforms that are similar to reference, some combination of the above, or only alternative isoforms that were unable to be classified (NA). P-values shown are from a two-sided Mann Whitney test. **G.** Effect of GRHL3 isoforms on differentiation, from the TF mORF Atlas.^61^ Top: exon diagrams of GRHL3 isoforms in TFIso1.0 and their categorizations as negative regulators or rewirers. Right: PPI profiles of GRHL3 isoforms from Y2H assays. Bottom: heatmap showing the percent of GRHL3 isoform-expressing cells in Louvain clusters, from the TF mORF Atlas. **H.** Paired swarm plot showing the total CREB1 gene expression levels in matched breast cancer tumor and normal samples. P-value shown is from a two-sided Mann Whitney test. **I.** Histogram showing the paired difference in total CREB1 gene expression across matched breast cancer tumor and normal samples. **J.** Histogram showing the paired difference in the relative expression of the alternative isoform of CREB1 (as a fraction of total CREB1 gene expression) across matched breast cancer tumor and normal samples.

## References

1. Ptashne, M. (1988). How eukaryotic transcriptional activators work. Nature 335, 683–689.

2. Jolma, A., Kivioja, T., Toivonen, J., Cheng, L., Wei, G., Enge, M., Taipale, M., Vaquerizas, J.M., Yan, J., Sillanpää, M.J., et al. (2010). Multiplexed massively parallel SELEX for characterization of human transcription factor binding specificities. Genome Res. 20, 861–873.

3. Jolma, A., Yan, J., Whitington, T., Toivonen, J., Nitta, K.R., Rastas, P., Morgunova, E., Enge, M., Taipale, M., Wei, G., et al. (2013). DNA-binding specificities of human transcription factors. Cell 152, 327–339.

4. Fuxman Bass, J.I., Sahni, N., Shrestha, S., Garcia-Gonzalez, A., Mori, A., Bhat, N., Yi, S., Hill, D.E., Vidal, M., and Walhout A.J.M. (2015). Human gene-centered transcription factor networks for enhancers and disease variants. Cell 161, 661–673.

5. Barrera, L.A., Vedenko, A., Kurland, J.V., Rogers, J.M., Gisselbrecht, S.S., Rossin, E.J., Woodard, J., Mariani, L., Kock, K.H., Inukai, S., et al. (2016). Survey of variation in human transcription factors reveals prevalent DNA binding changes. Science 351, 1450–1454.

6. Tycko, J., DelRosso, N., Hess, G.T., Aradhana, Banerjee, A., Mukund, A., Van, M.V., Ego, B.K., Yao, D., Spees K., et al. (2020). High-Throughput Discovery and Characterization of Human Transcriptional Effectors. Cell 183, 2020–2035.e16.

7. DelRosso, N., Tycko, J., Suzuki, P., Andrews, C., Aradhana, Mukund, A., Liongson, I., Ludwig, C., Spees, K., Fordyce P., et al. (2023). Large-scale mapping and mutagenesis of human transcriptional effector domains. Nature 616, 365–372.

8. Soto, L.F., Li, Z., Santoso, C.S., Berenson, A., Ho, I., Shen, V.X., Yuan, S., and Fuxman Bass, J.I. (2022). Compendium of human transcription factor effector domains. Mol. Cell 82, 514–526.

9. Grove, C.A., De Masi, F., Barrasa, M.I., Newburger, D.E., Alkema, M.J., Bulyk, M.L., and Walhout, A.J.M. (2009). A multiparameter network reveals extensive divergence between C. elegans bHLH transcription factors. Cell 138, 314–327.

10. Ravasi, T., Suzuki, H., Cannistraci, C.V., Katayama, S., Bajic, V.B., Tan, K., Akalin, A., Schmeier, S., Kanamori-Katayama, M., Bertin, N., et al. (2010). An Atlas of Combinatorial Transcriptional Regulation in Mouse and Man. Cell 140, 744–752.

11. Göös, H., Kinnunen, M., Salokas, K., Tan, Z., Liu, X., Yadav, L., Zhang, Q., Wei, G.-H., and Varjosalo, M. (2022). Human transcription factor protein interaction networks. Nat. Commun. 13, 766.

12. Nilsen, T.W., and Graveley, B.R. (2010). Expansion of the eukaryotic proteome by alternative splicing. Nature 463, 457–463.

13. Blencowe, B.J. (2006). Alternative splicing: new insights from global analyses. Cell 126, 37–47.

14. Aebersold, R., Agar, J.N., Amster, I.J., Baker, M.S., Bertozzi, C.R., Boja, E.S., Costello, C.E., Cravatt, B.F., Fenselau, C., Garcia, B.A., et al. (2018). How many human proteoforms are there? Nat. Chem. Biol. 14, 206–214.

15. Cahan, P., Li, H., Morris, S.A., Lummertz da Rocha, E., Daley, G.Q., and Collins, J.J. (2014). CellNet: network biology applied to stem cell engineering. Cell 158, 903–915.

16. Sahni, N., Yi, S., Taipale, M., Fuxman Bass, J.I., Coulombe-Huntington, J., Yang, F., Peng, J., Weile, J., Karras, G.I., Wang, Y., et al. (2015). Widespread macromolecular interaction perturbations in human genetic disorders. Cell 161, 647–660.

17. Scarpato, M., Federico, A., Ciccodicola, A., and Costa, V. (2015). Novel transcription factor variants through RNA-sequencing: the importance of being “alternative.” Int. J. Mol. Sci. 16, 1755–1771.

18. Talavera, D., Orozco, M., and de la Cruz, X. (2009). Alternative splicing of transcription factors’ genes: beyond the increase of proteome diversity. Comp. Funct. Genomics, 905894.

19. Wang, E.T., Sandberg, R., Luo, S., Khrebtukova, I., Zhang, L., Mayr, C., Kingsmore, S.F., Schroth, G.P., and Burge, C.B. (2008). Alternative isoform regulation in human tissue transcriptomes. Nature 456, 470–476.

20. Lambert, S.A., Jolma, A., Campitelli, L.F., Das, P.K., Yin, Y., Albu, M., Chen, X., Taipale, J., Hughes, T.R., and Weirauch, M.T. (2018). The Human Transcription Factors. Cell 172, 650–665.

21. Frankish, A., Diekhans, M., Ferreira, A.-M., Johnson, R., Jungreis, I., Loveland, J., Mudge, J.M., Sisu, C., Wright, J., Armstrong, J., et al. (2019). GENCODE reference annotation for the human and mouse genomes. Nucleic Acids Res. 47, D766–D773.

22. Deveson, I.W., Brunck, M.E., Blackburn, J., Tseng, E., Hon, T., Clark, T.A., Clark, M.B., Crawford, J., Dinger, M.E., Nielsen, L.K., et al. (2018). Universal Alternative Splicing of Noncoding Exons. Cell Syst 6, 245–255.e5.

23. Sheynkman, G.M., Tuttle, K.S., Laval, F., Tseng, E., Underwood, J.G., Yu, L., Dong, D., Smith, M.L., Sebra, R., Willems, L., et al. (2020). ORF Capture-Seq as a versatile method for targeted identification of full-length isoforms. Nat. Commun. 11, 2326.

24. Cooper, T.A., Wan, L., and Dreyfuss, G. (2009). RNA and disease. Cell 136, 777–793.

25. Harrow, J., Frankish, A., Gonzalez, J.M., Tapanari, E., Diekhans, M., Kokocinski, F., Aken, B.L., Barrell, D., Zadissa, A., Searle, S., et al. (2012). GENCODE: the reference human genome annotation for The ENCODE Project. Genome Res. 22, 1760–1774.

26. Sinitcyn, P., Richards, A.L., Weatheritt, R.J., Brademan, D.R., Marx, H., Shishkova, E., Meyer, J.G., Hebert, A.S., Westphall, M.S., Blencowe, B.J., et al. (2023). Global detection of human variants and isoforms by deep proteome sequencing. Nat. Biotechnol. 10.1038/s41587-023-01714-x.

27. López, A.J. (1995). Developmental role of transcription factor isoforms generated by alternative splicing. Dev. Biol. 172, 396–411.

28. Kelemen, O., Convertini, P., Zhang, Z., Wen, Y., Shen, M., Falaleeva, M., and Stamm, S. (2013). Function of alternative splicing. Gene 514, 1–30.

29. Taneri, B., Snyder, B., Novoradovsky, A., and Gaasterland, T. (2004). Alternative splicing of mouse transcription factors affects their DNA-binding domain architecture and is tissue specific. Genome Biol. 5, R75.

30. Vuzman, D., and Levy, Y. (2012). Intrinsically disordered regions as affinity tuners in protein-DNA interactions. Mol. Biosyst. 8, 47–57.

31. Li, J., Wang, Y., Rao, X., Wang, Y., Feng, W., Liang, H., and Liu, Y. (2017). Roles of alternative splicing in modulating transcriptional regulation. BMC Syst. Biol. 11, 89.

32. Gabut, M., Samavarchi-Tehrani, P., Wang, X., Slobodeniuc, V., O’Hanlon, D., Sung, H.-K., Alvarez, M., Talukder, S., Pan, Q., Mazzoni, E.O., et al. (2011). An alternative splicing switch regulates embryonic stem cell pluripotency and reprogramming. Cell 147, 132–146.

33. Barbaux, S., Niaudet, P., Gubler, M.C., Grünfeld, J.P., Jaubert, F., Kuttenn, F., Fékété, C.N., Souleyreau-Therville, N., Thibaud, E., Fellous, M., et al. (1997). Donor splice-site mutations in WT1 are responsible for Frasier syndrome. Nat. Genet. 17, 467–470.

34. Gregoire, E.P., De Cian, M.-C., Migale, R., Perea-Gomez, A., Schaub, S., Bellido-Carreras, N., Stévant, I., Mayère, C., Neirijnck, Y., Loubat, A., et al. (2023). The-KTS splice variant of WT1 is essential for ovarian determination in mice. Science 382, 600–606.

35. Bickmore, W.A., Oghene, K., Little, M.H., Seawright, A., van Heyningen, V., and Hastie, N.D. (1992). Modulation of DNA binding specificity by alternative splicing of the Wilms tumor wt1 gene transcript. Science 257, 235–237.

36. Belluti, S., Rigillo, G., and Imbriano, C. (2020). Transcription Factors in Cancer: When Alternative Splicing Determines Opposite Cell Fates. Cells 9. 10.3390/cells9030760.

37. Schaefer, T.S., Sanders, L.K., Park, O.K., and Nathans, D. (1997). Functional differences between Stat3alpha and Stat3beta. Mol. Cell. Biol. 17, 5307–5316.

38. Zhang, H.-X., Yang, P.-L., Li, E.-M., and Xu, L.-Y. (2019). STAT3beta, a distinct isoform from STAT3. Int. J. Biochem. Cell Biol. 110, 130–139.

39. Flouriot, G., Brand, H., Denger, S., Metivier, R., Kos, M., Reid, G., Sonntag-Buck, V., and Gannon, F. (2000). Identification of a new isoform of the human estrogen receptor-alpha (hER-α) that is encoded by distinct transcripts and that is able to repress hER-α activation function 1. EMBO J. 19, 4688–4700.

40. Shi, L., Dong, B., Li, Z., Lu, Y., Ouyang, T., Li, J., Wang, T., Fan, Z., Fan, T., Lin, B., et al. (2009). Expression of ER-α36, a novel variant of estrogen receptor α, and resistance to tamoxifen treatment in breast cancer. J. Clin. Oncol. 27, 3423–3429.

41. Gadea, G., Arsic, N., Fernandes, K., Diot, A., Joruiz, S.M., Abdallah, S., Meuray, V., Vinot, S., Anguille, C., Remenyi, J., et al. (2016). TP53 drives invasion through expression of its Δ133p53β variant. Elife 5. 10.7554/eLife.14734.

42. Yang, X., Coulombe-Huntington, J., Kang, S., Sheynkman, G.M., Hao, T., Richardson, A., Sun, S., Yang, F., Shen, Y.A., Murray, R.R., et al. (2016). Widespread Expansion of Protein Interaction Capabilities by Alternative Splicing. Cell 164, 805–817.

43. Lee, T.I., Rinaldi, N.J., Robert, F., Odom, D.T., Bar-Joseph, Z., Gerber, G.K., Hannett, N.M., Harbison, C.T., Thompson, C.M., Simon, I., et al. (2002). Transcriptional regulatory networks in Saccharomyces cerevisiae. Science 298, 799–804.

44. Weirauch, M.T., Yang, A., Albu, M., Cote, A.G., Montenegro-Montero, A., Drewe, P., Najafabadi, H.S., Lambert, S.A., Mann, I., Cook, K., et al. (2014). Determination and inference of eukaryotic transcription factor sequence specificity. Cell 158, 1431–1443.

45. Badis, G., Berger, M.F., Philippakis, A.A., Talukder, S., Gehrke, A.R., Jaeger, S.A., Chan, E.T., Metzler, G., Vedenko, A., Chen, X., et al. (2009). Diversity and complexity in DNA recognition by transcription factors. Science 324, 1720–1723.

46. Mukund, A.X., Tycko, J., Allen, S.J., Robinson, S.A., Andrews, C., Sinha, J., Ludwig, C.H., Spees, K., Bassik, M.C., and Bintu, L. (2023). High-throughput functional characterization of combinations of transcriptional activators and repressors. Cell Syst 14, 746–763.e5.

47. Reece-Hoyes, J.S., Pons, C., Diallo, A., Mori, A., Shrestha, S., Kadreppa, S., Nelson, J., Diprima, S., Dricot, A., Lajoie, B.R., et al. (2013). Extensive rewiring and complex evolutionary dynamics in a C. elegans multiparameter transcription factor network. Mol. Cell 51, 116–127.

48. Glinos, D.A., Garborcauskas, G., Hoffman, P., Ehsan, N., Jiang, L., Gokden, A., Dai, X., Aguet, F., Brown, K.L., Garimella, K., et al. (2022). Transcriptome variation in human tissues revealed by long-read sequencing. Nature 608, 353–359.

49. Veiga, D.F.T., Nesta, A., Zhao, Y., Deslattes Mays, A., Huynh, R., Rossi, R., Wu, T.-C., Palucka, K., Anczukow, O., Beck, C.R., et al. (2022). A comprehensive long-read isoform analysis platform and sequencing resource for breast cancer. Sci Adv 8, eabg6711.

50. Morales, J., Pujar, S., Loveland, J.E., Astashyn, A., Bennett, R., Berry, A., Cox, E., Davidson, C., Ermolaeva, O., Farrell, C.M., et al. (2022). A joint NCBI and EMBL-EBI transcript set for clinical genomics and research. Nature 604, 310–315.

51. Kriventseva, E.V., Koch, I., Apweiler, R., Vingron, M., Bork, P., Gelfand, M.S., and Sunyaev, S. (2003). Increase of functional diversity by alternative splicing. Trends Genet. 19, 124–128.

52. Alekseyenko, A.V., Kim, N., and Lee, C.J. (2007). Global analysis of exon creation versus loss and the role of alternative splicing in 17 vertebrate genomes. RNA 13, 661–670.

53. GTEx Consortium (2015). Human genomics. The Genotype-Tissue Expression (GTEx) pilot analysis: multitissue gene regulation in humans. Science 348, 648–660.

54. Cardoso-Moreira, M., Halbert, J., Valloton, D., Velten, B., Chen, C., Shao, Y., Liechti, A., Ascenção, K., Rummel, C., Ovchinnikova, S., et al. (2019). Gene expression across mammalian organ development. Nature 571, 505–509.

55. Raj, B., O’Hanlon, D., Vessey, J.P., Pan, Q., Ray, D., Buckley, N.J., Miller, F.D., and Blencowe, B.J. (2011). Cross-regulation between an alternative splicing activator and a transcription repressor controls neurogenesis. Mol. Cell 43, 843–850.

56. Wilanowski, T., Tuckfield, A., Cerruti, L., O’Connell, S., Saint, R., Parekh, V., Tao, J., Cunningham, J.M., and Jane, S.M. (2002). A highly conserved novel family of mammalian developmental transcription factors related to Drosophila grainyhead. Mech. Dev. 114, 37–50.

57. Weber, D., Wiese, C., and Gessler, M. (2014). Hey bHLH transcription factors. Curr. Top. Dev. Biol. 110, 285–315.

58. Vanorny, D.A., and Mayo, K.E. (2017). The role of Notch signaling in the mammalian ovary. Reproduction 153, R187–R204.

59. Melé, M., Ferreira, P.G., Reverter, F., DeLuca, D.S., Monlong, J., Sammeth, M., Young, T.R., Goldmann, J.M., Pervouchine, D.D., Sullivan, T.J., et al. (2015). Human genomics. The human transcriptome across tissues and individuals. Science 348, 660–665.

60. Ng, A.H.M., Khoshakhlagh, P., Rojo Arias, J.E., Pasquini, G., Wang, K., Swiersy, A., Shipman, S.L., Appleton, E., Kiaee, K., Kohman, R.E., et al. (2021). A comprehensive library of human transcription factors for cell fate engineering. Nat. Biotechnol. 39, 510–519.

61. Joung, J., Ma, S., Tay, T., Geiger-Schuller, K.R., Kirchgatterer, P.C., Verdine, V.K., Guo, B., Arias-Garcia, M.A., Allen, W.E., Singh, A., et al. (2023). A transcription factor atlas of directed differentiation. Cell 186, 209–229.e26.

62. Liu, J., Perumal, N.B., Oldfield, C.J., Su, E.W., Uversky, V.N., and Dunker, A.K. (2006). Intrinsic disorder in transcription factors. Biochemistry 45, 6873–6888.

63. Boija, A., Klein, I.A., Sabari, B.R., Dall’Agnese, A., Coffey, E.L., Zamudio, A.V., Li, C.H., Shrinivas, K., Manteiga, J.C., Hannett, N.M., et al. (2018). Transcription Factors Activate Genes through the Phase-Separation Capacity of Their Activation Domains. Cell 175, 1842–1855.e16.

64. Sabari, B.R., Dall’Agnese, A., Boija, A., Klein, I.A., Coffey, E.L., Shrinivas, K., Abraham, B.J., Hannett, N.M., Zamudio, A.V., Manteiga, J.C., et al. (2018). Coactivator condensation at super-enhancers links phase separation and gene control. Science 361. 10.1126/science.aar3958.

65. Brodsky, S., Jana, T., Mittelman, K., Chapal, M., Kumar, D.K., Carmi, M., and Barkai, N. (2020). Intrinsically Disordered Regions Direct Transcription Factor In Vivo Binding Specificity. Mol. Cell 79, 459–471.e4.

66. Reece-Hoyes, J.S., Diallo, A., Lajoie, B., Kent, A., Shrestha, S., Kadreppa, S., Pesyna, C., Dekker, J., Myers, C.L., and Walhout, A.J.M. (2011). Enhanced yeast one-hybrid assays for high-throughput gene-centered regulatory network mapping. Nat. Methods 8, 1059–1064.

67. Luck, K., Kim, D.-K., Lambourne, L., Spirohn, K., Begg, B.E., Bian, W., Brignall, R., Cafarelli, T., Campos-Laborie, F.J., Charloteaux, B., et al. (2020). A reference map of the human binary protein interactome. Nature 580, 402–408.

68. Albert, R. (2005). Scale-free networks in cell biology. J. Cell Sci. 118, 4947–4957.

69. Bedard, J.E.J., Haaning, A.M., and Ware, S.M. (2011). Identification of a novel ZIC3 isoform and mutation screening in patients with heterotaxy and congenital heart disease. PLoS One 6, e23755.

70. Ma, L., Sham, Y.Y., Walters, K.J., and Towle, H.C. (2007). A critical role for the loop region of the basic helix-loop-helix/leucine zipper protein Mlx in DNA binding and glucose-regulated transcription. Nucleic Acids Res. 35, 35–44.

71. Fuke, S. (2005). Identification and Characterization of the Hesr1/Hey1 as a Candidate trans-Acting Factor on Gene Expression through the 3’ Non-Coding Polymorphic Region of the Human Dopamine Transporter (DAT1) Gene. Preprint, 10.1093/jb/mvi02010.1093/jb/mvi020.

72. Englert, C., Vidal, M., Maheswaran, S., Ge, Y., Ezzell, R.M., Isselbacher, K.J., and Haber, D.A. (1995). Truncated WT1 mutants alter the subnuclear localization of the wild-type protein. Proc. Natl. Acad. Sci. U. S. A. 92, 11960–11964.

73. Larsson, S.H., Charlieu, J.P., Miyagawa, K., Engelkamp, D., Rassoulzadegan, M., Ross, A., Cuzin, F., van Heyningen, V., and Hastie, N.D. (1995). Subnuclear localization of WT1 in splicing or transcription factor domains is regulated by alternative splicing. Cell 81, 391–401.

74. Siggers, T., Reddy, J., Barron, B., and Bulyk, M.L. (2014). Diversification of Transcription Factor Paralogs via Noncanonical Modularity in C2H2 Zinc Finger DNA Binding. Mol. Cell 55, 640–648.

75. Eddy, S.R. (2011). Accelerated Profile HMM Searches. PLoS Comput. Biol. 7, e1002195.

76. Berger, M.F., Philippakis, A.A., Qureshi, A.M., He, F.S., Estep, P.W., 3rd, and Bulyk, M.L. (2006). Compact, universal DNA microarrays to comprehensively determine transcription-factor binding site specificities. Nat. Biotechnol. 24, 1429–1435.

77. Berger, M.F., and Bulyk, M.L. (2009). Universal protein-binding microarrays for the comprehensive characterization of the DNA-binding specificities of transcription factors. Nat. Protoc. 4, 393–411.

78. Kock, K.H., Kimes, P.K., Gisselbrecht, S.S., Inukai, S., Phanor, S.K., Anderson, J.T., Ramakrishnan, G., Lipper, C.H., Song, D., Kurland, J.V., et al. (2023). DNA binding analysis of rare variants in homeodomains reveals novel homeodomain specificity-determining residues. bioRxiv, 2023.06.16.545320. 10.1101/2023.06.16.545320.

79. Steimle, J.D., and Moskowitz, I.P. (2017). TBX5: A Key Regulator of Heart Development. Curr. Top. Dev. Biol. 122, 195–221.

80. Li, Q.Y., Newbury-Ecob, R.A., Terrett, J.A., Wilson, D.I., Curtis, A.R., Yi, C.H., Gebuhr, T., Bullen, P.J., Robson, S.C., Strachan, T., et al. (1997). Holt-Oram syndrome is caused by mutations in TBX5, a member of the Brachyury (T) gene family. Nat. Genet. 15, 21–29.

81. Yamak, A., Georges, R.O., Sheikh-Hassani, M., Morin, M., Komati, H., and Nemer, M. (2015). Novel exons in the tbx5 gene locus generate protein isoforms with distinct expression domains and function. J. Biol. Chem. 290, 6844–6856.

82. Oki, S., Ohta, T., Shioi, G., Hatanaka, H., Ogasawara, O., Okuda, Y., Kawaji, H., Nakaki, R., Sese, J., and Meno, C. (2018). ChIP-Atlas: a data-mining suite powered by full integration of public ChIP-seq data. EMBO Rep. 19. 10.15252/embr.201846255.

83. Brodsky, S., Jana, T., and Barkai, N. (2021). Order through disorder: The role of intrinsically disordered regions in transcription factor binding specificity. Curr. Opin. Struct. Biol. 71, 110–115.

84. Mosca, R., Céol, A., Stein, A., Olivella, R., and Aloy, P. (2014). 3did: a catalog of domain-based interactions of known three-dimensional structure. Nucleic Acids Res. 42, D374–D379.

85. Shen, W.-K., Chen, S.-Y., Gan, Z.-Q., Zhang, Y.-Z., Yue, T., Chen, M.-M., Xue, Y., Hu, H., and Guo, A.-Y. (2023). AnimalTFDB 4.0: a comprehensive animal transcription factor database updated with variation and expression annotations. Nucleic Acids Res. 51, D39–D45.

86. Zhang, C.-H., Gao, Y., Jadhav, U., Hung, H.-H., Holton, K.M., Grodzinsky, A.J., Shivdasani, R.A., and Lassar, A.B. (2021). Creb5 establishes the competence for Prg4 expression in articular cartilage. Commun Biol 4, 332.

87. Hornbeck, P.V., Zhang, B., Murray, B., Kornhauser, J.M., Latham, V., and Skrzypek, E. (2015). PhosphoSitePlus, 2014: mutations, PTMs and recalibrations. Nucleic Acids Res. 43, D512–D520.

88. Naqvi, S., Martin, K.J., and Arthur, J.S.C. (2014). CREB phosphorylation at Ser133 regulates transcription via distinct mechanisms downstream of cAMP and MAPK signalling. Biochem. J 458, 469–479.

89. Kopelman, N.M., Lancet, D., and Yanai, I. (2005). Alternative splicing and gene duplication are inversely correlated evolutionary mechanisms. Nat. Genet. 37, 588–589.

90. Grishkevich, V., and Yanai, I. (2014). Gene length and expression level shape genomic novelties. Genome Res. 24, 1497–1503.

91. Lister, J.A., Close, J., and Raible, D.W. (2001). Duplicate mitf genes in zebrafish: complementary expression and conservation of melanogenic potential. Dev. Biol. 237, 333–344.

92. Hultman, K.A., Bahary, N., Zon, L.I., and Johnson, S.L. (2007). Gene Duplication of the zebrafish kit ligand and partitioning of melanocyte development functions to kit ligand a. PLoS Genet. 3, e17.

93. Marshall, A.N., Montealegre, M.C., Jiménez-López, C., Lorenz, M.C., and van Hoof, A. (2013). Alternative splicing and subfunctionalization generates functional diversity in fungal proteomes. PLoS Genet. 9, e1003376.

94. Laudet, V. (2011). The origins and evolution of vertebrate metamorphosis. Curr. Biol. 21, R726–R737.

95. Ortiga-Carvalho, T.M., Sidhaye, A.R., and Wondisford, F.E. (2014). Thyroid hormone receptors and resistance to thyroid hormone disorders. Nat. Rev. Endocrinol. 10, 582–591.

96. Talavera, D., Vogel, C., Orozco, M., Teichmann, S.A., and de la Cruz, X. (2007). The (in)dependence of alternative splicing and gene duplication. PLoS Comput. Biol. 3, e33.

97. Cramer, P. (2019). Organization and regulation of gene transcription. Nature 573, 45–54.

98. Ji, D., Shao, C., Yu, J., Hou, Y., Gao, X., Wu, Y., Wang, L., and Chen, P. (2023). FOXA1 forms biomolecular condensates that unpack condensed chromatin to function as a pioneer factor. Mol. Cell. 10.1016/j.molcel.2023.11.020.

99. Tripathi, S., Shirnekhi, H.K., Gorman, S.D., Chandra, B., Baggett, D.W., Park, C.-G., Somjee, R., Lang, B., Hosseini, S.M.H., Pioso, B.J., et al. (2023). Defining the condensate landscape of fusion oncoproteins. Nat. Commun. 14, 6008.

100. Thul, P.J., Åkesson, L., Wiking, M., Mahdessian, D., Geladaki, A., Ait Blal, H., Alm, T., Asplund, A., Björk, L., Breckels, L.M., et al. (2017). A subcellular map of the human proteome. Science 356, eaal3321.

101. Sabari, B.R. (2020). Biomolecular Condensates and Gene Activation in Development and Disease. Dev. Cell 55, 84–96.

102. Wei, M.-T., Chang, Y.-C., Shimobayashi, S.F., Shin, Y., Strom, A.R., and Brangwynne, C.P. (2020). Nucleated transcriptional condensates amplify gene expression. Nat. Cell Biol. 22, 1187–1196.

103. Veiga, R.N., de Oliveira, J.C., and Gradia, D.F. (2021). PBX1: a key character of the hallmarks of cancer. J. Mol. Med. 99, 1667–1680.

104. Mary, L., Leclerc, D., Gilot, D., Belaud-Rotureau, M.-A., and Jaillard, S. (2022). The TALE never ends: A comprehensive overview of the role of PBX1, a TALE transcription factor, in human developmental defects. Hum. Mutat. 43, 1125–1148.

105. Linares, A.J., Lin, C.-H., Damianov, A., Adams, K.L., Novitch, B.G., and Black, D.L. (2015). The splicing regulator PTBP1 controls the activity of the transcription factor Pbx1 during neuronal differentiation. Elife 4, e09268.

106. Remesal, L., Roger-Baynat, I., Chirivella, L., Maicas, M., Brocal-Ruiz, R., Pérez-Villalba, A., Cucarella, C., Casado, M., and Flames, N. (2020). PBX1 acts as terminal selector for olfactory bulb dopaminergic neurons. Development 147. 10.1242/dev.186841.

107. Xu, Y., Zhao, W., Olson, S.D., Prabhakara, K.S., and Zhou, X. (2018). Alternative splicing links histone modifications to stem cell fate decision. Genome Biol. 19, 133.

108. Riback, J.A., Zhu, L., Ferrolino, M.C., Tolbert, M., Mitrea, D.M., Sanders, D.W., Wei, M.-T., Kriwacki, R.W., and Brangwynne, C.P. (2020). Composition-dependent thermodynamics of intracellular phase separation. Nature 581, 209–214.

109. Qian, D., Ausserwoger, H., Arter, W.E., Scrutton, R.M., Welsh, T.J., Kartanas, T., Ermann, N., Qamar, S., Fischer, C., Sneideris, T., et al. (2023). Linking modulation of bio-molecular phase behaviour with collective interactions. bioRxiv, 2023.11.02.565376. 10.1101/2023.11.02.565376.

110. Hsiao, P.W., and Chang, C. (1999). Isolation and characterization of ARA160 as the first androgen receptor N-terminal-associated coactivator in human prostate cells. J. Biol. Chem. 274, 22373–22379.

111. Volpe, M., Shpungin, S., Barbi, C., Abrham, G., Malovani, H., Wides, R., and Nir, U. (2006). trnp: A conserved mammalian gene encoding a nuclear protein that accelerates cell-cycle progression. DNA Cell Biol. 25, 331–339.

112. Esgleas, M., Falk, S., Forné, I., Thiry, M., Najas, S., Zhang, S., Mas-Sanchez, A., Geerlof, A., Niessing, D., Wang, Z., et al. (2020). Trnp1 organizes diverse nuclear membrane-less compartments in neural stem cells. EMBO J. 39, e103373.

113. Fisher, S.E., and Scharff, C. (2009). FOXP2 as a molecular window into speech and language. Trends Genet. 25, 166–177.

114. Bruce, H.A., and Margolis, R.L. (2002). FOXP2: novel exons, splice variants, and CAG repeat length stability. Hum. Genet. 111, 136–144.

115. Schroeder, D.I., and Myers, R.M. (2008). Multiple transcription start sites for FOXP2 with varying cellular specificities. Gene 413, 42–48.

116. Burkett, Z.D., Day, N.F., Kimball, T.H., Aamodt, C.M., Heston, J.B., Hilliard, A.T., Xiao, X., and White, S.A. (2018). FoxP2 isoforms delineate spatiotemporal transcriptional networks for vocal learning in the zebra finch. Elife 7. 10.7554/eLife.30649.

117. Mizutani, A., Matsuzaki, A., Momoi, M.Y., Fujita, E., Tanabe, Y., and Momoi, T. (2007). Intracellular distribution of a speech/language disorder associated FOXP2 mutant. Biochem. Biophys. Res. Commun. 353, 869–874.

118. Tanabe, Y., Fujiwara, Y., Matsuzaki, A., Fujita, E., Kasahara, T., Yuasa, S., and Momoi, T. (2012). Temporal expression and mitochondrial localization of a Foxp2 isoform lacking the forkhead domain in developing Purkinje cells. J. Neurochem. 122, 72–80.

119. UniProt Consortium (2023). UniProt: the Universal Protein Knowledgebase in 2023. Nucleic Acids Res. 51, D523–D531.

120. Zammarchi, F., de Stanchina, E., Bournazou, E., Supakorndej, T., Martires, K., Riedel, E., Corben, A.D., Bromberg, J.F., and Cartegni, L. (2011). Antitumorigenic potential of STAT3 alternative splicing modulation. Proc. Natl. Acad. Sci. U. S. A. 108, 17779–17784.

121. Gasperoni, J.G., Fuller, J.N., Darido, C., Wilanowski, T., and Dworkin, S. (2022). Grainyhead-like (Grhl) Target Genes in Development and Cancer. Int. J. Mol. Sci. 23. 10.3390/ijms23052735.

122. Peyrard-Janvid, M., Leslie, E.J., Kousa, Y.A., Smith, T.L., Dunnwald, M., Magnusson, M., Lentz, B.A., Unneberg, P., Fransson, I., Koillinen, H.K., et al. (2014). Dominant mutations in GRHL3 cause Van der Woude Syndrome and disrupt oral periderm development. Am. J. Hum. Genet. 94, 23–32.

123. Frisch, S.M., Farris, J.C., and Pifer, P.M. (2017). Roles of Grainyhead-like transcription factors in cancer. Oncogene 36, 6067–6073.

124. Sands, W.A., and Palmer, T.M. (2008). Regulating gene transcription in response to cyclic AMP elevation. Cell. Signal. 20, 460–466.

125. Sapio, L., Salzillo, A., Ragone, A., Illiano, M., Spina, A., and Naviglio, S. (2020). Targeting CREB in Cancer Therapy: A Key Candidate or One of Many? An Update. Cancers 12. 10.3390/cancers12113166.

126. Spies, N., Burge, C.B., and Bartel, D.P. (2013). 3’ UTR-isoform choice has limited influence on the stability and translational efficiency of most mRNAs in mouse fibroblasts. Genome Res. 23, 2078–2090.

127. Walhout, A.J.M., Temple, G.F., Brasch, M.A., Hartley, J.L., Lorson, M.A., van den Heuvel, S., and Vidal, M. (2000). GATEWAY recombinational cloning: Application to the cloning of large numbers of open reading frames or ORFeomes. Methods in Enzymology 328, 575–592.

128. Herskowitz, I. (1987). Functional inactivation of genes by dominant negative mutations. Nature 329, 219–222.

129. Koenig, R.J., Lazar, M.A., Hodin, R.A., Brent, G.A., Larsen, P.R., Chin, W.W., and Moore, D.D. (1989). Inhibition of thyroid hormone action by a non-hormone binding c-erbA protein generated by alternative mRNA splicing. Nature 337, 659–661.

130. Shen, N., Zhao, J., Schipper, J.L., Zhang, Y., Bepler, T., Leehr, D., Bradley, J., Horton, J., Lapp, H., and Gordan, R. (2018). Divergence in DNA Specificity among Paralogous Transcription Factors Contributes to Their Differential In Vivo Binding. Cell Syst 6, 470–483.e8.

131. Zhang, Y., Ho, T.D., Buchler, N.E., and Gordân, R. (2021). Competition for DNA binding between paralogous transcription factors determines their genomic occupancy and regulatory functions. Genome Res. 31, 1216–1229.

132. Feng, S., Rastogi, C., Loker, R., Glassford, W.J., Tomas Rube, H., Bussemaker, H.J., and Mann, R.S. (2022). Transcription factor paralogs orchestrate alternative gene regulatory networks by context-dependent cooperation with multiple cofactors. Nat. Commun. 13, 3808.

133. Nitta, K.R., Jolma, A., Yin, Y., Morgunova, E., Kivioja, T., Akhtar, J., Hens, K., Toivonen, J., Deplancke, B., Furlong, E.E.M., et al. (2015). Conservation of transcription factor binding specificities across 600 million years of bilateria evolution. Elife 4. 10.7554/eLife.04837.

134. Brent, G.A. (2012). Mechanisms of thyroid hormone action. J. Clin. Invest. 122, 3035–3043.

135. Cheng, J., Novati, G., Pan, J., Bycroft, C., Žemgulytė, A., Applebaum, T., Pritzel, A., Wong, L.H., Zielinski, M., Sargeant, T., et al. (2023). Accurate proteome-wide missense variant effect prediction with AlphaMissense. Science 381, eadg7492.

136. Esposito, D., Weile, J., Shendure, J., Starita, L.M., Papenfuss, A.T., Roth, F.P., Fowler, D.M., and Rubin, A.F. (2019). MaveDB: an open-source platform to distribute and interpret data from multiplexed assays of variant effect. Genome Biol. 20, 223.

137. Rentzsch, P., Witten, D., Cooper, G.M., Shendure, J., and Kircher, M. (2019). CADD: predicting the deleteriousness of variants throughout the human genome. Nucleic Acids Res. 47, D886–D894.

138. Tewhey, R., Kotliar, D., Park, D.S., Liu, B., Winnicki, S., Reilly, S.K., Andersen, K.G., Mikkelsen, T.S., Lander, E.S., Schaffner, S.F., et al. (2016). Direct Identification of Hundreds of Expression-Modulating Variants using a Multiplexed Reporter Assay. Cell 165, 1519–1529.

139. Truty, R., Ouyang, K., Rojahn, S., Garcia, S., Colavin, A., Hamlington, B., Freivogel, M., Nussbaum, R.L., Nykamp, K., and Aradhya, S. (2021). Spectrum of splicing variants in disease genes and the ability of RNA analysis to reduce uncertainty in clinical interpretation. Am. J. Hum. Genet. 108, 696–708.

140. Bhattacharya, A., Vo, D.D., Jops, C., Kim, M., Wen, C., Hervoso, J.L., Pasaniuc, B., and Gandal, M.J. (2023). Isoform-level transcriptome-wide association uncovers genetic risk mechanisms for neuropsychiatric disorders in the human brain. Nat. Genet. 55, 2117–2128.

141. Bonnal, S.C., López-Oreja, I., and Valcárcel, J. (2020). Roles and mechanisms of alternative splicing in cancer-implications for care. Nat. Rev. Clin. Oncol. 17, 457–474.

142. Bushweller, J.H. (2019). Targeting transcription factors in cancer-from undruggable to reality. Nat. Rev. Cancer 19, 611–624.

143. Kryuchkova-Mostacci, N., and Robinson-Rechavi, M. (2017). A benchmark of gene expression tissue-specificity metrics. Brief. Bioinform. 18, 205–214.

