## Supplementary material for "Widespread variation in molecular interactions and regulatory properties among transcription factor isoforms": Methods

### STAR Methods

#### Resource Availability

##### Lead Contact

##### Materials Availability

The TFIso1.0 collection of TF isoform clones generated in this study are currently available through CCSB. Yeast strains used for eY1H assays are available from Juan I. Fuxman Bass upon request.

##### Data and Code Availability

All original code is available at github.com/CCSB-DFCI/TF_isoforms_paper. DOIs are listed in the key resources table. The code is additionally available as a repository on Github at. PBM data is available on GEO at accession GSE253638.

#### Experimental Model and Subject Details

##### Yeast strains

DNA-bait strains had been generated in the Y1HaS2 yeast background and were previously published (22037705, 25910213, 33179750). TF-prey strains were generated in the Yα1867 yeast background.

##### Human cell lines and cell culture

HEK293T cells were maintained in DMEM supplemented with 10% FBS and 1% antibiotic-antimycotic. Cells were passaged every 2 days at a ratio of 1:4, were kept in a sterile incubator at 37ºC and 5% $CO_2$, and regularly tested for mycoplasma contamination. Only low passage number cells were used in mammalian one-hybrid and NanoLuc two-hybrid experiments.

#### Method Details

##### TFIso1.0 clone collection generation and validation

A PCR-based method was used to amplify and clone the coding regions of TF isoforms, similarly to that in Yang *et al*.[1] Gene-specific anchoring primers (**Table S16**) were used to PCR amplify ORF sequences from reverse transcribed RNA obtained from fetal and adult brain, heart, and liver tissues obtained from BioChain.

| Catalog number | Description | Lot number |
|---|---|---|
| R1244149-50 | Total RNA - Human Fetal Normal Tissue: Liver | A601607 |
| R1234149-50 | Total RNA - Human Adult Normal Tissue: Liver | B705065 |
| R1244035-50 | Total RNA - Human Fetal Normal Tissue: Brain | B210035 |
| R1234035-50 | Total RNA - Human Adult Normal Tissue: Brain | B805061 |
| R1244122-50 | Total RNA - Human Fetal Normal Tissue: Heart | B512118 |
| R1234122-50 | Total RNA - Human Adult Normal Tissue: Heart | B604038 |

Primers were targeted against TF protein-coding transcripts based on sequences from GENCODE v21, with TFs defined as a union of two datasets: Reece-Hoyes *et al*.[2] and TFClass.[3] In subsequent data analysis, we updated the list of TFs used to Lambert *et al*.[4] and the GENCODE version to v30.

Cloning was performed in two stages: an initial pilot stage followed by the main stage, with clones resulting from both stages combined in the final collection. In the pilot stage, TFs targeted for cloning were selected based on long-read RNA-seq data of brain, heart, and liver, obtained from PacBio (private communication). In the full stage, the majority (> 90%) of the TFs targeted for cloning were selected based on having either protein-protein interactions (PPIs) or protein-DNA interactions (PDIs) in other ongoing single-isoform-per-gene PPI and PDI mapping projects from our labs. An additional 45 TF genes with the potential to have measurable differences in DNA binding were selected, the criteria for this was having an annotated alternative isoform with differences, relative to its reference isoform, in the DBD or in the 15 a.a. either side of the DBD, with both isoforms less than 80 KDa and less than 80% predicted disorder. Finally, an additional 25 TF genes with variants associated with neurodegenerative diseases (private communication) were added.

Cloning of isoforms of selected target genes was carried out as described previously.[1,5] Reverse transcription (RT) was carried out using a SMARTer® PCR cDNA Synthesis Kit (Takara) with oligo (dT)16 primers according to the manufacturer's instructions. The resultant cDNAs were used as templates for PCR amplification using KOD HotStart Polymerase (Novagen) and ORF-specific primers (Table S16). Up to 4 primer pairs targeting alternative N- and C-termini were used for each TF gene. The resulting amplicons, which may contain more than one alternatively spliced isoform were transferred into pDONR223 by Gateway™ BP reaction (Life Technologies) followed by transformation into *E. coli* DH5α. Transformed *E. coli* cells were plated on LB agar containing 50 mg/L spectinomycin for overnight growth at 37°C, after which up to 24 colonies were isolated for each primer pair using a Qpix 2 XT colony picking robot (Molecular Devices - Genetix).

These cloned TF isoforms were then combined with existing cloned TFs from our ORFeome collections.[1,6] PCR artifacts, duplicates, cases where there were multiple isoforms per well, and incomplete or otherwise erroneous cloned isoforms, were removed following Illumina short-read sequencing of the corresponding clones. The final set of ORFs was then chosen based on subsequent multiplexed, full-length, long-read sequencing.

The short-read sequencing step was performed mostly as described in Yang et al.,[1] pooling individual *E. coli* strains carrying plasmids encoding different TF genetic loci. Plasmid DNA minipreps from the pooled *E. coli* were prepared on a Qiagen BioRobot® Universal System according to the manufacturer's instructions and processed to make an Illumina sequencing library, during which Illumina adapter sequences, i7 and i5, were incorporated as plate indexes. The library was then paired-end-sequenced using an Illumina platform (MiSeq or NextSeq 500).

The long-read sequencing templates were generated by PCR using a method where the forward primers contained well specific barcodes and the reverse primers contained plate specific barcodes, enabling the clones to be pooled into a small number of aliquots that were processed for long-read sequencing on a PacBio RS II system (Pacific Biosciences of California, Inc.).

All isoforms were assessed by expert human annotators using methods and standards developed by GENCODE[7,8] to identify well supported isoforms suitable for further study. To assess orthogonal support for transcript structures, each isoform was aligned to the human reference genome (GRCh38) and compared to the contemporaneous GENCODE gene annotation, all available long transcriptomic data, RNAseq data and RNAseq supported introns from the Intropolis dataset[9] and CAGE transcription start site data.[10] Having determined support for transcript structure, putative novel CDSs were similarly investigated to assess support for translation initiation sites and novel coding exons using aligned protein sequence data, Ribo-seq data, and PhyloCSF[11] constraint data.

There were 28 cases where two or more clones encoded identical amino acid sequences. These were tested in the Y1H, M1H, and Y2H assays, and data from the duplicate clones were filtered out, keeping the clone that had the least drop-outs in the assays.

#### TF isoform annotations

Transcription factors were defined by Lambert et al. downloaded from http://humantfs.ccbr.utoronto.ca/ v1.01.[4]

Isoforms were matched to the CDS sequence of transcripts in the GENCODE basic set of GENCODE v30. Two cloned isoforms matched identical sequences to isoforms of two genes, HSFY1 and HSFY2, where the reference and alternative isoforms have identical CDS. We arbitrarily annotated these clones as HSFY1. For analysis, GENCODE transcripts with identical amino acid sequences but differences in the UTRs were merged into one protein isoform.

The reference isoform of a gene was defined in the vast majority of cases by the MANE select transcript. In the cases where a MANE select transcript was not available, the APPRIS principle isoform was used. If that was also not available, the longest isoform was chosen. The cloned reference isoform for a gene in TFiso1.0 was defined as the reference isoform, if it was cloned. If the reference isoform was not cloned, the cloned reference isoform was defined by the APPRIS annotated isoforms preferring principle over alternative. If no MANE or APPRIS annotated isoforms were cloned or available, then the cloned reference isoform was defined as the longest isoform matched to a GENCODE transcript. In the final case, if no GENCODE-matched isoforms were cloned, then the cloned reference isoform was the longest cloned isoform.

Alignment of TFIso1.0 isoforms with prior curated isoforms from the literature were manually determined based on the original evidence provided, including reported exon position and sequence length.

#### Detection of Protein-DNA interactions using enhanced yeast one-hybrid assays

Enhanced yeast one-hybrid (eY1H) assays were performed as described previously.[12–14] TF isoform ORF clones from TFIso1.0 were transferred by Gateway LR cloning (ThermoFisher #11791100) to the destination vector pDEST-AD2μ-*TRP1* (Walhout Lab)[12–14] to generate fusion clones of each TF isoform with the yeast Gal4 activation domain (AD).

To generate TF-prey yeast strains, cloned TF isoform ORFs were transformed into haploid MATα type yeast strain Yα1867, as previously described ([12–14]) and as follows. Yeast were inoculated in 1 L liquid YAPD media to a concentration of OD600 = 0.15 and were then incubated at 30°C shaking at 200 rpm until they reached OD600 = 0.5, washed with sterile water, and washed again with TE + 0.1 M lithium acetate (TE/LiAc). Yeast were resuspended in TE/LiAc with salmon sperm DNA (ThermoFisher #15632011) at a dilution of 1:10 before adding ~250 ng of the TF isoform clone. Six volumes of TE/LiAc + 40% polyethylene glycol were then added and samples were mixed gently ten times. Yeast were incubated at 30°C without shaking for 30 min followed by 42°C for 20 min, then resuspended in sterile water. Transformed yeast were plated on selective media lacking tryptophan to select for transformants.

DNA-bait yeast strains for 330 human enhancer and promoter sequences were previously generated using the Y1Has2 yeast strain.[13,15] For a complete list of baits with positive interactions, see **Table S2**. Each DNA-bait strain carries two integrated copies of the enhancer or promoter cloned upstream of two reporter genes: *HIS3*, which allows yeast to grow in the absence of histidine and overcome inhibition by 3-Amino-1,2,4-triazole (3AT), and *LacZ*, which causes yeast colonies to turn blue in the presence of 5-bromo-4-chloro-3-indolyl-beta-D-galacto-pyranoside (X-gal).

eY1H assays were performed in 1,536-colony format using a high-density array ROTOR robot (Singer Instruments), which facilitated the comparison between TF isoforms by allowing simultaneous testing on the same array plate. TF-prey yeast strains were mated in a pairwise manner with 211 DNA-bait strains on permissive YAPD agar plates and incubated at 30°C for one day. Yeast were then transferred to selective media agar plates lacking uracil and tryptophan and incubated at 30°C for two days to select for successfully mated diploid yeast. The resulting diploid yeast colonies were finally transferred to selective media agar plates lacking uracil, tryptophan, and histidine, with 320 mg/L X-gal and 5mM 3AT. Readout plates were imaged 2, 3, 4, and 7 days after plating. Binding of the TF-AD fusion to the DNA-bait region results in expression of the *HIS3* and *LacZ* reporter genes, allowing colonies to visibly grow and turn blue on readout plates. In addition, TF isoforms corresponding to MAX, STAT1, STAT3, PPARG, RARG, and RXRG, were tested against a collection of 119 cytokine promoter DNA-baits using paired yeast one-hybrid assays as previously described.[16] eY1H and paired yeast one-hybrid assay images were manually analyzed by three independent researchers to identify interactions. Array coordinate "holes" - where yeast mating or transfer was unsuccessful - were identified and removed from analysis.

#### Yeast one-hybrid protein-DNA interaction validation using luciferase assays

For a random subset of TF isoform series (i.e., all isoforms of a TF in TFiso1.0) a random subset of DNA baits were selected that had at least one interaction identified by eY1H assays with at least one of the isoforms. These all-by-all combinations for each selected TF isoform series were validated in an orthogonal system by luciferase assays in HEK293T cells (see **Table S7**). Briefly, DNA-bait sequences were cloned upstream of the firefly luciferase reporter in a Gateway compatible pGL4.23[luc2/minP] vector.[13] TF isoform ORFs were cloned into the Gateway compatible pEZY3-VP160 vector[15] such that TF isoforms are fused to 10 copies of the VP16 activation domain. HEK293T cells were plated in 96-well white opaque plates at a seeding density of ~10,000 cells/well and incubated for one day at 37ºC with 5% $CO_2$. Cells were transfected using Lipofectamine 3000 (Invitrogen) according to the manufacturer's protocol, with 80 ng of TF isoform (pEZY3-VP160) plasmid, 20 ng of DNA bait (pGL4.23) plasmid, and 10 ng of the renilla luciferase plasmid as a transfection normalization control. An empty pEZY3-VP160 plasmid co-transfected with the corresponding recombinant firefly luciferase plasmid were used as negative controls. Transfected cells were incubated for two days at 37ºC with 5% $CO_2$. Firefly and renilla luciferase activities were measured using the Dual-Glo Luciferase Assay System (Promega) according to the manufacturer's protocol. Non-transfected cells were used to subtract background firefly and renilla luciferase activities, and then firefly luciferase activity was normalized to renilla luciferase activity in each well. Each TF isoform-DNA bait pair was tested in three biological replicates. In the event of TF isoform-DNA binding, the VP16 activation domains promote the expression of firefly luciferase, increasing the normalized luminescence over background levels.

#### Protein-DNA interaction assay with protein-binding microarrays

Full-length TF isoforms (3 isoforms of TBX5 and 2 isoforms of CREB1) were cloned from Gateway compatible Entry vectors (pDONR223) into N-terminal GST protein fusion expression Destination vectors and sequence-verified by long-read DNA sequencing via Plasmidsaurus. Specifically, TBX5 isoforms were cloned into a modified pT7CFE1-NHis-GST vector (Thermo 88871), which is compatible with mammalian IVT, and CREB1 isoforms were cloned into pDEST15-NGST (Thermo 11802014), which is compatible with PURExpress IVT. Proteins were then expressed using either the 1-Step Human Coupled IVT Kit (for TBX5 isoforms) or PURExpress In Vitro Protein Synthesis Kit (for CREB1) (NEB E6800L), using the manufacturers' recommended protocols, with the exception of an addition of 1.5 µL of custom tRNA mix (NEB N6842Z) to the CREB1 PURExpress reactions. Protein expression was verified and quantified by Western blot using recombinant GST protein (Sigma G5663) as expression standards. Primary rabbit anti-GST polyclonal antibody (Sigma G7781) (1:160,000) and secondary goat horseradish peroxidase-conjugated IgG monoclonal antibody (Pierce 31460) (1:200,000) were used for Western blotting.

Universal PBMs containing all 10-mers in 8 x 60K, GSE format (Agilent Technologies: AMADID #030236) were used. For TBX5 arrays, PBM double-stranding was performed as previously described.[17,18] The polymerase used for the TBX5 arrays (Thermo Fisher Thermo Sequenase Cycle Sequencing Kit 785001KT) was discontinued by the manufacturer, and so, for CREB1 arrays PBM double stranding was performed using Cytiva Thermo Sequenase DNA polymerase (Cytiva E790000Y) using 3x polymerase but otherwise following the manufacturer's protocol. PBMs were then performed as described,[17,18] using a 50 µg/mL dilution of Alexa-488-conjugated rabbit polyclonal anti-GST antibody (Invitrogen A11131) in PBS / 2% (wt/vol) milk. CREB1 isoforms were assayed at 400 nM final concentration and TBX5 isoforms were assayed at 750 nM final concentration. PBMs were scanned in a GenePix 4400A microarray scanner. Each isoform was assayed on ≥ 2 independent arrays, and alternative isoforms were always assayed with their cognate reference isoforms on the same array. Given the higher level of noise when assaying full-length TFs compared to extended DBDs, each reference isoform was assayed on an additional ≥ 2 independent arrays to ensure robust quantification and differential comparisons.

#### Protein-protein interaction assay with yeast two-hybrid (Y2H)

##### Y2H Screens

The Y2H screens were performed mostly as described in Luck *et al.*[6] with some modifications.

***Yeast strains and transformation***

Competent yeast strain Y8800, mating type MATa (*leu2-3,112 trp1-901 his3Δ200 ura3-52 gal4Δ gal80Δ GAL2::ADE2 GAL1::HIS3@LYS2 GAL7::lacZ@MET2cyh2*$^R$) were transformed with individual AD-ORF constructs and plated onto yeast synthetic complete media[19] lacking tryptophan (SC-Trp) to select for AD-ORF plasmids.

Competent yeast strain Y8930, mating type MATα, (*leu2-3,112 trp1-901 his3Δ200 ura3-52 gal4Δ gal80Δ GAL2::ADE2 GAL1::HIS3@LYS2 GAL7::lacZ@MET2cyh2*$^R$) were transformed with individual DB-ORF constructs and plated onto SC-Leu to select for DB-ORF plasmids. Haploid DB-ORF yeast strains were tested for auto-activation of the *GAL1::HIS3* reporter gene. Individual DB-ORF yeast strains were spotted on SC-Leu-His+1mM 3AT media and any strains showing growth were considered auto-activators (AAs) and removed from the collection of strains to be screened.

***Primary Y2H Screens***

Two first-pass Y2H screens were performed, in which all TF isoforms were tested against (1) the hORFeome collection of ~17,500 ORF clones[6,19] and (2) a subset of the hORFeome collection that were annotated as TFs or co-factors. The list of co-factors was taken from the union of the TcoF database[20] using the January 2017 update, and from Heinäniemi *et al.*[21]

In both Y2H screens, TF isoforms were tested as fusions to the Gal4 activation domain (AD) in the Gateway compatible pDEST-AD-*CYH2* vector, and screened against the hORFeome fused to the Gal4 DNA-binding domain in the Gateway compatible pDEST-DB vector. To perform the screen pools of Y8930:DB-ORF yeast strains (baits) were mated against pools of Y8800:AD-ORF strains (preys). The TF isoforms AD-ORF yeast strains were combined into pools of 100 individual strains. In the large screen against the hORFeome the DB-ORF strains were combined into pools of 8 DB-ORF strains, and in the second focused screen DB-ORF strains were screened individually. These first-pass screens represent a systematic interrogation of ~13 million possible PPIs. To perform the mating, fresh overnight cultures of DB-ORF strains (either pools or individual) were mixed with AD-ORF strain pools and grown overnight at 30ºC in liquid rich media (YEPD). After overnight growth, the mated yeast cells were transferred into liquid SC-Leu-Trp media to select for diploids and again grown overnight at 30ºC. Finally the yeast cells were spotted onto SC-Leu-Trp-His+1mM 3AT solid media to select for activation of the *GAL1::HIS3* reporter gene. In parallel, diploid yeast cells were transferred onto SC-Leu-His+1mM 3AT solid media supplemented with 1 mg/l cycloheximide (CHX) to test for spontaneous DB-ORF auto-activators. All AD-ORF plasmids carry the counter-selectable marker *CYH2*, which allows selection on CHX-containing media of yeast cells that do not contain any AD-ORF plasmid. After 72h incubation at 30ºC, yeast that grew on SC-Leu-Trp-His+1mM 3AT media but not on SC-Leu-His+1mM 3AT+ 1 mg/l CHX media were picked into SC-Leu-Trp grown overnight and then processed to determine the identity of the respective bait and prey proteins. To identify the interacting bait and prey we used SWIM-Seq as described in Luck *et al.*[6]

The hits from the screen were then combined with PPIs from the subset of HuRI[6] that was detected using Y2H v1 with the TF as the AD fusion, and Lit-BM-17,[6] a dataset of literature-curated PPIs with multiple evidence including at least one experimental method that detects binary PPIs. These pairs were then tested in a series of initial pairwise Y2H experiments, testing each isoform of a TF gene against all interaction partners. These experiments were used to filter out pairs that were not positive with any of the isoforms of a TF gene, TF isoforms that were not positive with any interaction partner, and profiles of TF genes that were not positive (for at least one isoform) with at least two different partners and had at least two different isoforms with at least one positive interaction. These experiments were described as below, with the exception that the plate position of pairs was randomized, rather than keeping all isoforms of the same gene with the same partner on the same plate.

In the final pairwise test, we included additional pairs to test, so that we could compare the PPI profiles of paralogs to those of isoforms without the confounding effect of the sampling sensitivity of the screening. For a subset of paralogous TF genes and a control set of random paired non-paralogous genes (see the section on Paralogs definition), we additionally tested all isoforms of each paired gene with any additional interaction partners tested for the other gene.

##### Y2H Pairwise Test

Following first-pass screening, each protein isoform was pairwise tested for interaction with the candidate partners identified not only for itself but also for all first-pass partners of all other protein isoforms encoded by the same gene, thus minimizing biases due to incomplete sampling sensitivity.[22] To generate a final dataset of verified Y2H pairs, pairs were accepted if they showed (1) a valid growth score and (2) their ORF identities were confirmed by sequencing of the PCR products amplified from the tested colonies.

Briefly, interactors were inoculated in 200 µL corresponding selection media and mated overnight at 30ºC in 150 µL liquid rich media (YEPD). The following day, mated yeast cells were transferred into 150 µL liquid SC-Leu-Trp media to select for diploids. After overnight incubation at 30ºC, 5 µL diploid yeast cells were spotted onto SC-Leu-Trp-His+1mM 3AT solid media to select for activation of the *GAL1::HIS3* reporter gene as well as on SC-Leu-Trp to control for successful mating. AA tests were included by mating each Y8930:DB-ORF against a Y8800:AD-null (containing no ORF), which was included on each test plate for each individual Y8930:DB-ORF. Spots were scored for growth[6] with scores of 0, 1, 2, 3, 4, and NA for cases where the spotting had failed. Growth scores of 0 and 1 were considered not growing. If a spot corresponding to either the pair or the corresponding AA-test did not grow on SC-Leu-Trp, the pair was scored NA. If the AA-test had a growth score of 4, the corresponding pair was scored NA. Interactions were scored positive if they had higher growth scores on SC-Leu-Trp-His+1mM 3AT solid media compared to the auto-activator test and had a growth score of at least 2; otherwise they were scored negative. All positive scored colonies were picked, lysed and the identity of the two interacting proteins was confirmed performing SWIM-seq.[6] In addition, all SC-Leu-Trp plates were also sequenced. On each test plate, internal controls were included and in each batch of tested plates a positive reference set (PRS) and random reference set (RRS) was tested alongside the actual experiment.

#### Protein-protein interaction validation using mammalian NanoLuc two-hybrid (mN2H)

A random sample of PPIs identified by Y2H assays were validated in an orthogonal system by luciferase complementation assays in HEK293T cells,[23] along with positive and negative controls (**Table S6**). The tested positive Y2H pairs were a random sample of 300 pairs. The sample of negative pairs to test were selected in two steps: (1) negative pairs involving the same PPI partner with different isoforms of the same TF genes as the sampled positive pairs, where all negative pairs for each TF gene and partner combination were randomly selected to be included using a probability with a value of the fraction of positive pairs sampled for that gene; (2) an additional random sample of 150 negative pairs. The rationale behind this approach is that: by pairing the positive to negative pairs in step (1) we should reduce the variance when comparing positive to negative. The random selection in step (1) ensures a uniform random sample. Without it, the sample would be biased towards negative pairs with a larger number of positive PPIs involving other isoforms of the same TF gene. The additional pairs from step (2) were needed to increase the size of the negative sample to match the size of the positive sample.

In this assay, interacting partners were cloned into the corresponding N2H gateway plasmids (pDEST-N1 or pDEST-N2). Briefly, 30,000 HEK293T cells were seeded in a 96-well, flat bottom, cell culture microplate (Greiner Bio-One, #655083), and cultured in Dulbecco's modified Eagle's medium (DMEM) supplemented with 10% fetal calf serum at 37°C and 5% $CO_2$. Next day, cells were transfected with 100 ng of each N2H plasmid using linear polyethylenimine (PEI) to co-express the protein pairs fused with complementary NanoLuc fragments, F1 and F2. The following day, the media was removed and 50 µL of 100× diluted NanoLuc substrate (Promega, #N1110) or 100 × diluted furimazine substrate (Yves Janin) was added to each well of a 96-well microplate containing the transfected cells. Plates were incubated for 3 min at room temperature. Luciferase enzymatic activity was measured using a TriStar luminometer (Berthold; 2 s integration time).

#### Transcriptional activity using mammalian one-hybrid assays

To measure TF isoform transcriptional activity, modified mammalian one-hybrid assays (M1H) were performed in HEK293T cells. Briefly, TF isoform ORFs were cloned into a Gateway compatible DB-pEZY3 vector such that TF isoforms would be N-terminally fused to the Gal4 DNA-binding domain (DB). Four copies of the yeast UAS site corresponding to the Gal4 DB were then cloned upstream of the firefly luciferase reporter gene in a Gateway compatible pGL4.23[luc2/minP] vector. The DB-pEZY3 and 4xUAS-pGL4.23 backbone vectors were generated for this study (**Figure S2F**). HEK293T cells were plated in 96-well white opaque plates at a seeding density of ~10,000 cells/well and incubated for one day at 37$^o$C with 5% $CO_2$. Cells were then transfected using Lipofectamine 3000 (Invitrogen) according to the manufacturer's protocol, with 80 ng of TF isoform (DB-pEZY3) plasmid, 20 ng of 4xUAS-pGL4.23 plasmid, and 10 ng of the renilla luciferase plasmid as a transfection normalization control. An empty DB-pEZY3 plasmid co-transfected with the 4xUAS-pGL4.23 were used as negative controls. Cells were incubated for 2 days after transfection at 37$^o$C with 5% $CO_2$, and then firefly and renilla luciferase activities were measured using the Dual-Glo Luciferase Assay System (Promega) according to the manufacturer's protocol. Non-transfected cells were used to subtract background firefly and renilla luciferase activities, and then firefly luciferase activity was normalized to renilla luciferase activity in each well. In this assay, if the TF isoform is recruited to the UAS by the fused Gal4 DB this would lead to the expression or repression of the downstream firefly luciferase reporter genes depending on the endogenous activating or repressing activity of the TF isoform.

#### Condensate formation assay

##### Selection of isoforms for condensate assay

We selected 192 of our cloned isoforms (two 96 well-plates) to profile for condensate formation and localization in the condensates assay. We prioritized alternative isoforms based on showing differences from the reference isoform in either the PDI, PPI, or transcriptional activation assays. We restricted to genes where the MANE select isoform was cloned and the alternative isoform was cataloged in GENCODE. The assay criteria were: a difference in PDI profile with at least three DNA baits positive in at least one isoform of the gene; a difference in PPI profile, with at least three successfully tested PPIs, and at least one positive PPI for both the reference and alternative isoform; an 8-fold or greater difference in activation. These criteria were selected to try and get a roughly even split between differences in the three assays. The cloned reference and all alternative isoforms of a TF gene were selected, such that many other alternative TF isoforms that didn't show differences in the three assays were also included in those to be tested. This selection resulted in 50 TF genes. We then added an additional 11 TF genes which did not pass the selection above, but on manual inspection, showed differences between reference and alternative isoforms in our assay readouts that we judged to be interesting. We removed four cloned alternative isoforms that were not successfully tested in any of the three assays.

We transferred the reference and alternative isoform clones by Gateway LR reactions into a mammalian expression vector pcDNA3.1-ccdB-EGFP containing a C-terminal EGFP tag. All these clones were subjected to high-content imaging for condensate formation in two cell lines, HEK293T and U2OS. HEK293T and U2OS cells were transfected using standard protocols with FuGENE HD Transfection Reagent (Promega, Cat. No. E2311) in a 96-well plate format in DMEM and RPMI media, respectively, supplemented with 10% FBS and appropriate amounts of penicillin and streptomycin. 48 hours after transfection, cells were stained with DAPI, and imaging was performed using a ZEISS LSM 880 confocal microscope using a 63x objective. For comparative purposes, all available reference and alternative isoforms of the same gene were included in the same 96-well plate, for high-content imaging. We filtered out proteins that were not expressed from our imaging screen analysis. Alternative isoform-mediated condensate calls (Gain-of-condensate or GOC, Loss-of-condensate or LOC, and unchanged) were obtained, by comparing to their reference isoform profile (i.e., condensate or non-condensate). All phase separation experiments were performed in duplicate.

### Quantification and Statistical Analysis

#### Gene annotation

TF families were defined by Lambert *et al.*[4] MANE select transcripts were obtained from the file MANE.GRCh38.v0.95.summary.txt.[24] APPRIS transcript annotations were obtained from the file APPRIS-annotations_human_GRCh38.p13_ensembl104.tsv.[25]

#### Protein domain annotation

Pfam domains were mapped to protein isoforms from GENCODE v30 and our TFiso1.0 clone collection using HMMER version 3.3 and Pfam version 32.0. Domain matches were filtered for E-value < 0.01 and c-Evalue < 0.01. Overlapping Pfam domains were removed, keeping the domain with the lowest E-value. Zinc Finger domains, defined by membership of Pfam clan CL0361, that were separated by 10 amino acids or less, were merged into a single ZF array domain. We manually curated a list of Pfam domains that corresponded to DNA binding domains (Table S17). For TFiso1.0, we manually inspected each cloned reference isoform that did not have an annotated DBD, finding that for two TFs, HIF1A and ZNF207, their DBDs were above the E-value cutoff, and so we implemented a manual override of the filter in those two cases. Effector (activation and repression) domains were obtained from the literature-curated database TFRegDB[26] and from two published systematic tiling screens.[27,28] Nuclear localization and export sequence motifs (NLS/NES) were downloaded from UniProt on 2023-10-02. For the protein interaction partners, Pfam domains were filtered for E-value ≤ $10^{-5}$.

#### Proportion of alternative isoforms with domain affected

P-values and error bars for the fraction of domains in reference isoforms which are affected in alternative isoforms, were calculated using a null model where the domain is randomly positioned along the reference isoform. This is calculated by first, for each domain/reference-isoform/alternative-isoform combination, calculating a probability as the fraction of cases in which the alternative isoform affects a dummy domain, a contiguous set of amino acids the same length as the real domain in the reference isoform, of all possible positions of that dummy domain along the reference isoform. In the case of multiple domains of the same type on a single reference isoform, the probabilities of at least one of the domains being affected was calculated, assuming independence. This array of probabilities for a specific type of domain, with one value for each reference/alternative isoform pair, was used in a Poisson Binomial distribution to calculate p-values and confidence intervals. Because the independence assumption, in the case of multiple domains, is violated by the fact that domains do not overlap, we compared our approach with a more computationally intensive and less numerically precise approach of repeatedly randomly shuffling the positions of the domains along the reference isoform, not allowing overlap in the case of multiple domains of the same type, and we found that the two approaches gave consistent results (data not shown).

#### AlphaFold structural prediction

Predicted 3D structures of cloned TF isoforms were obtained using AlphaFold version 2.3.1. With options: --model_preset=monomer_ptm --db_preset=full_dbs --max_template_date=2023-05-05. To produce figures showing the approximate position of DNA relative to the isoform structure, the AlphaFold structures were aligned to experimental structures of the TF, or a homologous TF, bound to DNA. The experimental structures were manually selected after searching for the amino acid sequence of the DNA binding domain of the reference isoform on the PDB website. HEY1 was aligned to human CLOCK in CLOCK BMAL1 heterodimer, PDB ID 4H10; CREB1 was aligned to mouse CREB1 homodimer, PDB ID 1DH3; TBX5 was aligned to mouse TBX5, interacting with NKX2-5, PDB ID 5FLV.

#### Predicted disorder values

Binary per-residue predictions of being in a disordered region, for each cloned isoform, were derived from the AlphaFold predicted structures by[29]:
(i) calculating accessible surface area (ASA) and secondary structure using DSSP[30]
(ii) normalizing to relative solvent accessibility (RSA), using maximum ASA values from [31]
(iii) calculating a sliding-window average RSA value for each residue, using a window of 20 aa both sides of the residue in question
(iv) residues with this average RSA ≥ 0.5 were categorized as 'disordered', with RSA < 0.5 as 'structured'
(v) performed a correction for long alpha helices, which would generally be structured in binding, but have high solvent accessibility when looking at the monomer structure, for example in bZIP TFs. Residues within contiguous regions classed as alpha helix of 20 amino acids or longer were set to 'structured'.

#### PPI partner classification

Protein interaction partners of the TF isoforms were categorized into one of four categories: *TF*, *cofactor*, *signaling*, or *other*. *TF* was based on the Lambert *et al*.[4] list, *cofactor* were proteins that were not classed as *TF* but appeared in the list of human cofactors from Animal TF DB v4.[32] *Signaling* were those partners not already classed as *TF* or *cofactor*, that were annotated with the gene ontology (GO) term 'signaling' (GO:0023052) or one of its related lower terms. GO annotations were generated by UniProt on 2023-07-28 and the ontology file was released 2023-07-27. All remaining partner proteins were classed as *other*.

We obtained a list of TF families that typically bind DNA as obligate heterodimers from Jolma et al. Nat Meth. 2013.[33] The 22 families are: AP-2, ARID/BRIGHT, BED ZF, bHLH, bZIP, CENPB, E2F, EBF1, GCM, Grainyhead, HSF, IRF, MADF, MADS box, Myb/SANT, Nuclear receptor, p53, RFX, Rel, SAND, SMAD, STAT.

#### Domain-domain PPI annotation

A list of interacting domain pairs was obtained from 3did[34] 2022-05 release. All possible domain pairs matching to the TF isoform and partner protein were initially mapped. These 61 domain pairs and their corresponding evidence were manually inspected, filtered for quality and duplicates were removed, resulting in a filtered list of 42 domain pairs. Further de-duplication was performed by collapsing the multiple different Pfam domains corresponding to bZIPs (bZIP_1/bZIP_2/bZIP_Maf), homeobox (Homeobox/Homeobox_KN), and PAS (PAS/PAS_3/PAS_9/PAS11) to a single domain each.

#### RNA-seq analyses

RNA-seq analyses were performed by pseudoaligning reads to transcriptome indices made using the following reference fasta files: (1) to estimate the relative abundance of annotated transcription factor isoforms (GENCODE version 30) alone (i.e., analyses in Figure 1), we used the GENCODE version 30 protein-coding transcripts fasta file (which includes full transcript sequences, including UTRs) as the reference, and (2) to estimate the abundance of both annotated TF isoforms and unannotated cloned isoforms in the TFIso1.0 collection (i.e., analyses in Figures 2-7), we produced a consensus fasta reference that includes the aforementioned GENCODE version 30 protein-coding transcripts as well as the CDS sequences of any unannotated, novel clones in our collection. In both cases, we generated index files for the software kallisto (version 0.46.0) using the "kallisto index" command with default parameters. Pseudoalignment was then performed on fastq files using the "kallisto quant" command with default parameters to estimate transcript per million (TPM) estimates per isoform. To estimate relative isoform abundance, gene-level TPM values were computed as the sum of all isoform TPM values for a given gene, and individual isoform ratios were determined relative to the total gene TPM for any gene with a TPM > 1. GTEx data were downloaded from the Sequence Read Archive prior to migration of the data to ANVIL following dbGAP approval (phs000424.v8.p2). Developmental RNA-seq data from Cardoso-Moreira et al.[35] were downloaded from ArrayExpress (accession number E-MTAB-6814). TCGA breast cancer data were downloaded as paired-end bam files from the NCI Genomics Data Commons portal following dbGAP approval (phs000178.v10.p8) and converted to paired-end fastq files using samtools (version 1.15). For these analyses, we included a subset of representative GTEx samples (n=1,201 samples), spanning the same 30 patients (where possible) for all 51 tissue regions (excluding cell lines).

##### Re-sampling GTEx data

Since the number of samples per condition and the number of conditions in the GTEx and Developmental RNA-seq datasets was very different, in order to compare isoform expression between adult and developing tissues, we created a randomly sampled subset of the GTEx dataset. To clarify: a condition in GTEx is one adult tissue type (e.g. "Liver") and a condition in Developmental RNA-seq is a tissue/time-point (e.g. "Liver 10 weeks post conception"). There were 1-5 samples per condition, with a median of 2 samples and a total of 127 conditions in Developmental RNA-seq, and 5-379 samples per condition, with a median of 24 and a total of 51 conditions in GTEx. To generate the resampled GTEx dataset we cycled through the GTEx tissues creating dummy conditions by randomly sampling the total number and number of samples per condition of the Developmental RNA-seq dataset.

#### Pairwise sequence identity analyses

Amino acid sequences were aligned using the pairwise2 module of biopython, with the blosum62 substitution matrix, an open gap penalty of -10, an extend gap penalty of -0.5, and penalize_end_gaps=False.

#### Paralogs definition

Paralogs were downloaded from Ensembl Compara on 2023-10-26. The non-paralog control set of pairs was generated from the list of paralog pairs, by randomly re-pairing the genes, removing any pairs that were in the original paralogs list of any cases where a gene was paired with itself. Note that there are some pairs in the non-paralog control of TF genes from the same family. We repeated the analysis with a different non-paralog control that specifically excludes within-family pairs and it produced very similar results (data not shown).

#### Functional assay quantification

Jaccard distances of PPI and PDI data for a pair of TF isoforms were calculated as 1 - number of common interaction partners / total number of interaction partners. Only partners that were successfully tested in both isoforms were included. For Y2H PPI data, we did not use values where one of the isoforms had no interactions, in order to try and avoid artifacts where the clone was not functional in the assay.

#### Violin plots

Violin plots were drawn with the Gaussian KDE in the python package seaborn but modified to fit bounded data by reflecting the probability density back from the bounds, with the bounds being 0-1 in the case of PDI/PPI profile Jaccard distance, a lower bound of 0 for absolute log2 fold change of activation, and 0%-100% for sequence similarity. The kernel bandwidth was set to 0.1/10% for Jaccard distance/sequence similarity and 0.5 for absolute log2 fold change of activation.

#### PBM analysis

For all PBM replicates, a scan corresponding to a photomultiplier tube (PMT) gain of 500 was selected, as this consistently resulted in the lowest proportion of both over-saturated and under-saturated probes. PBM pre-processing was then performed using the upbm data analysis pipeline as described previously.[36] Briefly, probe intensities are background-subtracted, Cy3-normalized (to account for any biases resulting from double-stranding the array), and spatially de-biased. Reference isoforms then served as “anchors” for cross-array normalization for each gene. PBM inference to determine differentially bound DNA 8-mers was also performed using the upbm pipeline, which tests for a difference in the aggregated 8-mer affinity scores against a null hypothesis of zero. GPR files are available on GEO at accession GSE253638.

#### TBX5 ChIP-seq analysis

Uniformly processed human TBX5 ChIP-seq peaks were downloaded (bed and bigwig files) from ChIP-Atlas.[37] Only studies profiling wild-type TBX5 were considered (see **Resources Table** for a list of Accession IDs), and only peaks with MACS2 q-values < $1 \times 10^{-5}$ were considered. Any overlapping peaks were de-duplicated using the bedtools[38] merge command, such that only one peak from one study was considered (randomly sampled) in any overlapping regions, resulting in a final list of 2,074 non-overlapping TBX5 ChIP peaks. Peak regions were then centered to the nucleotide with the highest ChIP signal and trimmed to 150 nucleotides using bwtool[39] and hg38 genomic sequences were extracted for each centered peak using bedtools.[38] GENRE[40] was then used to generate a list of matched genomic background sequences (of the same length) for k-mer enrichment analyses.

#### CSat analysis

Microscopy images were acquired on a Zeiss LSM 880 confocal laser scanning microscope equipped with a Plan-Apochromat 63x/1.4 oil DIC M27 objective with a pinhole size of 46 µm (1 Airy unit). mEGFP (expressed fused on the c-terminal of constructs) was excited with a 488 laser and imaged with emission filters 498-552 nm. As Csat curve analysis requires (1) construct expression levels that differ >10 fold and (2) ensuring a linear range of intensities in both condensates and the dilute phase (i.e., nucleoplasm or cytoplasm) which typically differ by >10 fold, laser power is optimized for each field. This optimization is done by the user in real-time at the scope with the guiding principles that the max pixel should be roughly 10% of the max of the detector, maximizing signal to noise while maintaining the linearity of the image digital units with a concentration of construct. Quantitative comparison between the different laser settings is achieved by converting each image to digital units (after background subtraction) referenced at 1% laser power (reported as AU). This conversion was done using an empirically measured conversion factor formula determined with conversion factors calculated from the slope of the pixel values when imaging the same field of view at 1% laser power and various other laser settings, only including those pixels within linear range in both images. Note that these images were not used for Csat analysis as pixels were frequently out of linear range, and each field was only imaged once to avoid the complication of photobleaching. All other microscope and camera settings were kept constant. For analysis, cells or nuclei at different expression levels were found and the rough region containing them was hand-segmented with polygons to remove extra-cellular debris and other nearby cells. The designation of containing condensates was user decided at this point prior to Csat quantification. Measurement of the dilute phase concentration was done manually by choosing a non-foci-containing location and getting the value at that pixel with a maximal Gaussian blur to lower that noise but without including foci. To approximate the total concentration (x-axis) the average pixel value was calculated in a 2D image in a binary mask for the cell. This binary mask for the cell was determined by taking the image in the polygon segmented region, blurring it with a pixel radius of 5, doing a morphological binarization using the dilute phase value, filling all holes in the object, and deleting small components. The exact command in Mathematica is DeleteSmallComponents[FillingTransform[MorphologicalBinarize[Blur[image, 5], dil]]] where image and dil is the polygon segmented image region and digital value for the dilute phase chosen, respectively. This was done using a custom-made Mathematica GUI based on that used in Riback et al. Nature 2020[41].

#### Human Protein Atlas localization validation

Subcellular localization data from the Human Protein Atlas[42] were downloaded on October 26, 2023. The following Human Protein Atlas subcellular localization annotations were considered “cytoplasmic”: Actin filaments, Cleavage furrow, Focal adhesion sites, Intermediate filaments, Centriolar satellite, Centrosome, Cytokinetic bridge, Microtubule ends, Microtubules, Midbody, Midbody ring, Mitotic spindle, Aggresome, Cytoplasmic bodies, Cytosol, Rods & rings, Mitochondria, Endoplasmic reticulum, Vesicles, Endosomes, Lipid droplets, Lysosomes, Peroxisomes, Golgi apparatus, Cell junctions, Plasma membrane. If one of the above localizations was observed in any localization column (Approved, Enhanced, Supported, Uncertain), we considered the protein to be “cytoplasmic” in **Figure S6F**.

#### Classification of negative regulators and rewirers

We defined negative regulator alternative isoforms as those exhibiting one or more of the following: (1) those that show 0 PDIs while their cognate reference isoform shows ≥ 1 PDI or those that lose ≥ 10% of the DBD; (2) those that show loss of activation compared to their reference isoform (reference has M1H signal ≥ 1, alternative has M1H signal between 1 and -1, and alternative log2FC ≤ -1 compared to reference) or loss of repression compared to their reference isoform (reference has M1H signal ≤ -1, alternative has M1H signal between 1 and -1, and alternative log2FC ≥ 1 compared to reference); (3) those that show 0 PPIs while their cognate reference isoform shows ≥ 1 PPI or those that lose all of 1 key type of PPI (within-family TFs of obligate dimers, signaling proteins, or transcriptional cofactors). To classify an alternative isoform as a negative regulator, we required that it show evidence of functionality in 1 additional assay as follows: (1) ≥ 1 PDI, (2) ≥ 1 PPI, or (3) M1H signal ≥ 1 (activation) or ≤ -1 (repression). Thus, only alternative isoforms with data from ≥ 2 assays (eY1H, Y2H, or M1H) were classified based on assay data alone; unless we considered the isoform a negative regulator due to loss of DBD (in those cases, evidence of functionality could come from Y2H or M1H assays alone). Finally, we layered subcellular localization on top of these categorizations (we did not consider it with the same initial weight as the Y1H, Y2H, and M1H assays as localization in and of itself is not evidence of TF functionality): any alternative isoforms whose localization changed from nuclear or both nuclear/cytoplasmic in the reference to solely cytoplasmic in either HEK293T or U2OS imaging assays were considered negative regulators. Any alternative isoforms with data in ≥ 2 functional assays (eY1H, Y2H, or M1H) that had ≥ 1 difference in PPIs or PDIs, or log2FC ≥ 1 or ≤ -1 (with at least 1 isoform having M1H signal above baseline), or a difference in subcellular localization in either cell line, were considered to be rewirers; any with 0 difference in PPIs or PDIs, and log2FC between -1 and 1, and no differences in localization in either cell line were considered to be similar to the reference isoform. Only one isoform loses function across all tested axes (**PPARG-3**, see Figure S7C) and was filtered out of downstream analyses as likely non-functional.

#### TF Atlas mORF analyses

The following datasets were downloaded from the TF over-expression atlas (Joung et al.[43]): (1) TF ORF library sequences (Supplemental Table S1A), (2) TF over-expression scores (processed, Supplemental Table S2B), and (3) mapping of TF ORFs to Louvain clusters from their Figure 3B (direct communication with authors). To intersect our clone collection with their library, we first attempted to match based on amino acid sequence; if this failed, we attempted to match based on annotated Ensembl transcript IDs. To examine the effect of TF over-expression on differentiation, we used their processed “Diffusion difference” as the effect size and “Diffusion P-value” as the p-value. To examine the enrichment of TF ORFs in differentiated cell types, we calculated the percentage of TF-expressing cells in a given Louvain cluster relative to the total number of TF-expressing cells in their subsampled library.

##### Paired tumor/normal TCGA analysis

We considered a set of 112 paired (i.e., from the same patient) tumor/normal breast cancer samples (***SuppTable_BRCASamps.txt***). In 5 cases, there were repeat normal control samples from the same patient – we randomly sampled 1 of these control samples in each case to make the final list. To find isoforms that show significant differences between tumor and normal controls, we performed paired Wilcoxon tests between the fractional isoform expression in the 112 normal samples compared to the fractional isoform expression in the 112 tumor samples, requiring a minimum gene-level expression of 1 TPM in both normal and tumor samples in a minimum of 10 such paired samples. We then corrected these p-values for multiple hypothesis testing using the Benjamini-Hochberg method and an FDR of 0.05.

#### Additional Resources

We have made the website tfisodb.org to enable users to browse and download the data presented in this paper, including isoform diagrams, expression levels, and assay results.

#### References


1. Yang, X., Coulombe-Huntington, J., Kang, S., Sheynkman, G.M., Hao, T., Richardson, A., Sun, S., Yang, F., Shen, Y.A., Murray, R.R., et al. (2016). Widespread Expansion of Protein Interaction Capabilities by Alternative Splicing. Cell *164*, 805–817.

2. Reece-Hoyes, J.S., Barutcu, A.R., McCord, R.P., Jeong, J.S., Jiang, L., MacWilliams, A., Yang, X., Salehi-Ashtiani, K., Hill, D.E., Blackshaw, S., et al. (2011). Yeast one-hybrid assays for gene-centered human gene regulatory network mapping. Nat. Methods *8*, 1050–1052.

3. Wingender, E., Schoeps, T., Haubrock, M., Krull, M., and Dönitz, J. (2018). TFClass: expanding the classification of human transcription factors to their mammalian orthologs. Nucleic Acids Res. *46*, D343–D347.

4. Lambert, S.A., Jolma, A., Campitelli, L.F., Das, P.K., Yin, Y., Albu, M., Chen, X., Taipale, J., Hughes, T.R., and Weirauch, M.T. (2018). The Human Transcription Factors. Cell *172*, 650–665.

5. Salehi-Ashtiani, K., Yang, X., Derti, A., Tian, W., Hao, T., Lin, C., Makowski, K., Shen, L., Murray, R.R., Szeto, D., et al. (2008). Isoform discovery by targeted cloning, “deep-well” pooling and parallel sequencing. Nat. Methods *5*, 597–600.

6. Luck, K., Kim, D.-K., Lambourne, L., Spirohn, K., Begg, B.E., Bian, W., Brignall, R., Cafarelli, T., Campos-Laborie, F.J., Charloteaux, B., et al. (2020). A reference map of the human binary protein interactome. Nature *580*, 402–408.

7. Harrow, J., Frankish, A., Gonzalez, J.M., Tapanari, E., Diekhans, M., Kokocinski, F., Aken, B.L., Barrell, D., Zadissa, A., Searle, S., et al. (2012). GENCODE: the reference human genome annotation for The ENCODE Project. Genome Res. *22*, 1760–1774.

8. Frankish, A., Carbonell-Sala, S., Diekhans, M., Jungreis, I., Loveland, J.E., Mudge, J.M.,

Sisu, C., Wright, J.C., Arnan, C., Barnes, I., et al. (2023). GENCODE: reference annotation for the human and mouse genomes in 2023. Nucleic Acids Res. *51*, D942–D949.

9. Nellore, A., Jaffe, A.E., Fortin, J.-P., Alquicira-Hernández, J., Collado-Torres, L., Wang, S., Phillips, R.A., III, Karbhari, N., Hansen, K.D., Langmead, B., et al. (2016). Human splicing diversity and the extent of unannotated splice junctions across human RNA-seq samples on the Sequence Read Archive. Genome Biol. *17*, 266.

10. FANTOM Consortium and the RIKEN PMI and CLST (DGT), Forrest, A.R.R., Kawaji, H., Rehli, M., Baillie, J.K., de Hoon, M.J.L., Haberle, V., Lassmann, T., Kulakovskiy, I.V., Lizio, M., et al. (2014). A promoter-level mammalian expression atlas. Nature *507*, 462–470.

11. Lin, M.F., Jungreis, I., and Kellis, M. (2011). PhyloCSF: a comparative genomics method to distinguish protein coding and non-coding regions. Bioinformatics *27*, i275–i282.

12. Reece-Hoyes, J.S., Diallo, A., Lajoie, B., Kent, A., Shrestha, S., Kadreppa, S., Pesyna, C., Dekker, J., Myers, C.L., and Walhout, A.J.M. (2011). Enhanced yeast one-hybrid assays for high-throughput gene-centered regulatory network mapping. Nat. Methods *8*, 1059–1064.

13. Fuxman Bass, J.I., Sahni, N., Shrestha, S., Garcia-Gonzalez, A., Mori, A., Bhat, N., Yi, S., Hill, D.E., Vidal, M., and Walhout, A.J.M. (2015). Human gene-centered transcription factor networks for enhancers and disease variants. Cell *161*, 661–673.

14. Sahni, N., Yi, S., Taipale, M., Fuxman Bass, J.I., Coulombe-Huntington, J., Yang, F., Peng, J., Weile, J., Karras, G.I., Wang, Y., et al. (2015). Widespread macromolecular interaction perturbations in human genetic disorders. Cell *161*, 647–660.

15. Santoso, C.S., Li, Z., Lal, S., Yuan, S., Gan, K.A., Agosto, L.M., Liu, X., Pro, S.C., Sewell, J.A., Henderson, A., et al. (2020). Comprehensive mapping of the human cytokine gene regulatory network. Nucleic Acids Res. *48*, 12055–12073.

16. Berenson, A., Lane, R., Soto-Ugaldi, L.F., Patel, M., Ciausu, C., Li, Z., Chen, Y., Shah, S., Santoso, C., Liu, X., et al. (2023). Paired yeast one-hybrid assays to detect DNA-binding cooperativity and antagonism across transcription factors. Nat. Commun. *14*, 6570.

17. Berger, M.F., and Bulyk, M.L. (2006). Protein binding microarrays (PBMs) for rapid, high-throughput characterization of the sequence specificities of DNA binding proteins. Methods Mol. Biol. *338*, 245–260.

18. Berger, M.F., Philippakis, A.A., Qureshi, A.M., He, F.S., Estep, P.W., 3rd, and Bulyk, M.L. (2006). Compact, universal DNA microarrays to comprehensively determine transcription-factor binding site specificities. Nat. Biotechnol. *24*, 1429–1435.

19. Dreze, M., Monachello, D., Lurin, C., Cusick, M.E., Hill, D.E., Vidal, M., and Braun, P. (2010). High-quality binary interactome mapping. Methods Enzymol. *470*, 281–315.

20. Schmeier, S., Alam, T., Essack, M., and Bajic, V.B. (2017). TcoF-DB v2: update of the database of human and mouse transcription co-factors and transcription factor interactions. Nucleic Acids Res. *45*, D145–D150.

21. Heinäniemi, M., Nykter, M., Kramer, R., Wienecke-Baldacchino, A., Sinkkonen, L., Zhou, J.X., Kreisberg, R., Kauffman, S.A., Huang, S., and Shmulevich, I. (2013). Gene-pair

expression signatures reveal lineage control. Nat. Methods *10*, 577–583.

22. Venkatesan, K., Rual, J.-F., Vazquez, A., Stelzl, U., Lemmens, I., Hirozane-Kishikawa, T., Hao, T., Zenkner, M., Xin, X., Goh, K.-I., et al. (2009). An empirical framework for binary interactome mapping. Nat. Methods *6*, 83–90.

23. Choi, S.G., Olivet, J., Cassonnet, P., Vidalain, P.-O., Luck, K., Lambourne, L., Spirohn, K., Lemmens, I., Dos Santos, M., Demeret, C., et al. (2019). Maximizing binary interactome mapping with a minimal number of assays. Nat. Commun. *10*, 3907.

24. Morales, J., Pujar, S., Loveland, J.E., Astashyn, A., Bennett, R., Berry, A., Cox, E., Davidson, C., Ermolaeva, O., Farrell, C.M., et al. (2022). A joint NCBI and EMBL-EBI transcript set for clinical genomics and research. Nature. 10.1038/s41586-022-04558-8.

25. Rodriguez, J.M., Pozo, F., Cerdán-Vélez, D., Di Domenico, T., Vázquez, J., and Tress, M.L. (2022). APPRIS: selecting functionally important isoforms. Nucleic Acids Res. *50*, D54–D59.

26. Soto, L.F., Li, Z., Santoso, C.S., Berenson, A., Ho, I., Shen, V.X., Yuan, S., and Fuxman Bass, J.I. (2022). Compendium of human transcription factor effector domains. Mol. Cell *82*, 514–526.

27. Tycko, J., DelRosso, N., Hess, G.T., Aradhana, Banerjee, A., Mukund, A., Van, M.V., Ego, B.K., Yao, D., Spees, K., et al. (2020). High-Throughput Discovery and Characterization of Human Transcriptional Effectors. Cell *183*, 2020–2035.e16.

28. DelRosso, N., Tycko, J., Suzuki, P., Andrews, C., Aradhana, Mukund, A., Liongson, I., Ludwig, C., Spees, K., Fordyce, P., et al. (2023). Large-scale mapping and mutagenesis of human transcriptional effector domains. Nature. 10.1038/s41586-023-05906-y.

29. Akdel, M., Pires, D.E.V., Pardo, E.P., Jänes, J., Zalevsky, A.O., Mészáros, B., Bryant, P., Good, L.L., Laskowski, R.A., Pozzati, G., et al. (2022). A structural biology community assessment of AlphaFold2 applications. Nat. Struct. Mol. Biol. *29*, 1056–1067.

30. Kabsch, W., and Sander, C. (1983). Dictionary of protein secondary structure: pattern recognition of hydrogen-bonded and geometrical features. Biopolymers *22*, 2577–2637.

31. Tien, M.Z., Meyer, A.G., Sydykova, D.K., Spielman, S.J., and Wilke, C.O. (2013). Maximum Allowed Solvent Accessibilites of Residues in Proteins. PLoS One *8*, e80635.

32. Shen, W.-K., Chen, S.-Y., Gan, Z.-Q., Zhang, Y.-Z., Yue, T., Chen, M.-M., Xue, Y., Hu, H., and Guo, A.-Y. (2023). AnimalTFDB 4.0: a comprehensive animal transcription factor database updated with variation and expression annotations. Nucleic Acids Res. *51*, D39–D45.

33. Jolma, A., Yan, J., Whitington, T., Toivonen, J., Nitta, K.R., Rastas, P., Morgunova, E., Enge, M., Taipale, M., Wei, G., et al. (2013). DNA-binding specificities of human transcription factors. Cell *152*, 327–339.

34. Mosca, R., Céol, A., Stein, A., Olivella, R., and Aloy, P. (2014). 3did: a catalog of domain-based interactions of known three-dimensional structure. Nucleic Acids Res. *42*, D374–D379.

35. Cardoso-Moreira, M., Halbert, J., Valloton, D., Velten, B., Chen, C., Shao, Y., Liechti, A., Ascenção, K., Rummel, C., Ovchinnikova, S., et al. (2019). Gene expression across mammalian organ development. Nature *571*, 505–509.

36. Kock, K.H., Kimes, P.K., Gisselbrecht, S.S., Inukai, S., Phanor, S.K., Anderson, J.T., Ramakrishnan, G., Lipper, C.H., Song, D., Kurland, J.V., et al. (2023). DNA binding analysis of rare variants in homeodomains reveals novel homeodomain specificity-determining residues. bioRxiv, 2023.06.16.545320. 10.1101/2023.06.16.545320.

37. Oki, S., Ohta, T., Shioi, G., Hatanaka, H., Ogasawara, O., Okuda, Y., Kawaji, H., Nakaki, R., Sese, J., and Meno, C. (2018). ChIP-Atlas: a data-mining suite powered by full integration of public ChIP-seq data. EMBO Rep. *19*, e46255.

38. Quinlan, A.R., and Hall, I.M. (2010). BEDTools: a flexible suite of utilities for comparing genomic features. Bioinformatics *26*, 841–842.

39. Pohl, A., and Beato, M. (2014). bwtool: a tool for bigWig files. Bioinformatics *30*, 1618–1619.

40. Mariani, L., Weinand, K., Vedenko, A., Barrera, L.A., and Bulyk, M.L. (2017). Identification of Human Lineage-Specific Transcriptional Coregulators Enabled by a Glossary of Binding Modules and Tunable Genomic Backgrounds. Cell Syst *5*, 654.

41. Riback, J.A., Zhu, L., Ferrolino, M.C., Tolbert, M., Mitrea, D.M., Sanders, D.W., Wei, M.-T., Kriwacki, R.W., and Brangwynne, C.P. (2020). Composition-dependent thermodynamics of intracellular phase separation. Nature *581*, 209–214.

42. Thul, P.J., Åkesson, L., Wiking, M., Mahdessian, D., Geladaki, A., Ait Blal, H., Alm, T., Asplund, A., Björk, L., Breckels, L.M., et al. (2017). A subcellular map of the human proteome. Science *356*. 10.1126/science.aal3321.

43. Joung, J., Ma, S., Tay, T., Geiger-Schuller, K.R., Kirchgatterer, P.C., Verdine, V.K., Guo, B., Arias-Garcia, M.A., Allen, W.E., Singh, A., et al. (2023). A transcription factor atlas of directed differentiation. Cell *186*, 209–229.e26.
