## Supplementary tables for "Widespread variation in molecular interactions and regulatory properties among transcription factor isoforms": Supplementary table guide.pdf

### Table S1. TFIso1.0 library sequences, related to Figure 2.

- **clone_id:** TFIso1.0 isoform clone ID (used in most figures)
- **gene_symbol:** TF gene
- **isoform_status:** one of either: annotated reference, annotated alternative, novel reference, or novel alternative. Where the reference is chosen from the available cloned isoforms of a gene.
- **gencode_transcript_names:** GENCODE transcript names for annotated isoforms
- **ensembl_transcript_ids:** Ensembl transcript IDs for annotated isoforms
- **cds_seq**: nucleotide sequence of cloned isoform
- **aa_seq:** amino acid sequence of the isoform
- **tf_family:** TF DBD family

### Table S2. eY1H DNA Baits involved in PDIs, related to Figures 2 and 3.

- **bait_id:** internal bait ID
- **seq:** sequence of baits

### Table S3. PDI (eY1H) results, related to Figures 2 and 3.

- **gene_symbol**: TF gene
- **clone_id:** TFIso1.0 isoform clone ID (used in most figures)
- **all other columns:** internal bait IDs; cell will be TRUE if there is a PDI, FALSE if there is a verified non-PDI, or NA

### Table S4. Transcriptional activation/repression (M1H) results, related to Figures 2 and 4.

- **clone_id:** TFIso1.0 isoform clone ID (used in most figures)
- **gene_symbol:** TF gene symbol
- **M1H_rep1, 2, 3:** M1H data from each of the 3 replicates
- **M1H_mean**: mean of the 3 replicates

### Table S5. PPI (Y2H) results, related to Figures 2 and 4.

- **ad_clone_id:** TFIso1.0 isoform clone ID (used in most figures)
- **ad_gene_symbol:** gene symbol of the TF isoform (which is fused to Gal4-AD)
- **ad_orf_id**: our internal ID
- **db_gene_symbol:** gene symbol of the tested partner (which is fused to Gal4-DB)
- **db_orf_id**: our internal ID
- **Y2H_result**: TRUE if there is a PPI, FALSE if there is a verified non-PPI, or NA
- **db_gene_category**: category of the tested partner (one of: TF, cofactor, signaling, or other)

- **db_gene_cofactor_type**: for the cofactor genes, the cofactor type (one of: coactivator, corepressor, both, or unknown)

### Table S6: mN2H PPI validation results, related to Figure S2.

- **clone_id:** TF isoform ID.
- **gene_symbol_tf:** gene encoding TF isoform.
- **gene_symbol_partner:** PPI partner.
- **test_orf_ida:** internal ID of interactor**.**
- **test_orf_idb:** internal ID of interactor**.**
- **source:** *isoform positives/isoform negatives* Y2H positive/negative pairs randomly sampled from our dataset; *PRS - hPRS-v2/RRS - hRRS-v2* positive/random reference set controls used to compare across a range of previous experiments; *Lit-BM-13*/*RRS - from HuRI* a larger set of positive/negative controls used in Luck *et al*. *Nature* 2020; *Lit-BM - TF space specific*/*RRS - TF space specific* positive/negative controls restricted to PPIs involving transcription factors to test if these behave differently in the assay relative to the more general controls.
- **score_pair:** mN2H assay readout for the tested pair.
- **score_empty-N1:** readout testing one partner against an empty vector.
- **score_empty-N2:** readout testing the other partner against an empty vector.
- **log2 NLR:** log2(score_pair / max(score_empty-N1, score_empty-N2)).

### Table S7: PDI validation results, related to Figure S2.

- **gene_symbol:** TF gene
- **clone_id:** TF isoform ID
- **Bait:** DNA sequence ID
- **Y1H_result:** original PDI call from eY1H assay
- **Replicate1/2/3:** assay readout scores
- **Average (empty-pEZY3-VP160):** mean control value
- **Log2(FC):** log2(mean(Replicate1/2/3 / Average (empty-pEZY3-VP160)))

### Table S8. PDI (PBM) results for CREB1, related to Figure 3.

- **seq:** DNA 8-mer
- **CREB1-ref:** affinityEstimate, calculated using the upbm command kmerTestAffinity across replicate arrays, for the CREB1 reference isoform. This is the relative affinity that CREB1-ref has for a given DNA 8-mer.
- **CREB1-alt:** affinityEstimate, calculated using the upbm command kmerTestAffinity across replicate arrays, for the CREB1 alternative isoform. This is the relative affinity that CREB1-ref has for a given DNA 8-mer.
- **contrastAverage:** contrastAverage, calculated using the upbm command kmerTestContrast between CREB1-ref and CREB1-alt. This is the average affinity across CREB1-ref and CREB1-alt for a given DNA 8-mer.

- **contrastDifference:** contrastDifference, calculated using the upbm command kmerTestContrast between CREB1-ref and CREB1-alt. This is the difference in affinity between CREB1-alt and CREB1-ref for a given DNA 8-mer.
- **contrastQ**: contrastQ, calculated using the upbm command kmerTestContrast between CREB1-ref and CREB1-alt. This is the q-value of the measured difference in affinity between CREB1-alt and CREB1-ref for a given DNA 8-mer.

### Table S9. PDI (PBM) results for TBX5, related to Figure 3.

- **seq:** DNA 8-mer
- **TBX5-ref:** affinityEstimate, calculated using the upbm command kmerTestAffinity across replicate arrays, for the TBX5 reference isoform. This is the relative affinity that TBX5-ref has for a given DNA 8-mer.
- **TBX5-2:** affinityEstimate, calculated using the upbm command kmerTestAffinity across replicate arrays, for the TBX5-2 alternative isoform. This is the relative affinity that TBX5-2 has for a given DNA 8-mer.
- **TBX5-3:** affinityEstimate, calculated using the upbm command kmerTestAffinity across replicate arrays, for the TBX5-3 alternative isoform. This is the relative affinity that TBX5-3 has for a given DNA 8-mer.
- **contrastAverage_TBX5-2:** contrastAverage, calculated using the upbm command kmerTestContrast between TBX5-ref and TBX5-2. This is the average affinity across TBX5-ref and TBX5-2 for a given DNA 8-mer.
- **contrastDifference_TBX5-2:** contrastDifference, calculated using the upbm command kmerTestContrast between TBX5-ref and TBX5-2. This is the difference in affinity between TBX5-2 and TBX5-ref for a given DNA 8-mer.
- **contrastQ_TBX5-2**: contrastQ, calculated using the upbm command kmerTestContrast between TBX5-ref and TBX5-2. This is the q-value of the measured difference in affinity between TBX5-2 and TBX5-ref for a given DNA 8-mer.
- **contrastAverage_TBX5-3:** contrastAverage, calculated using the upbm command kmerTestContrast between TBX5-ref and TBX5-3. This is the average affinity across TBX5-ref and TBX5-3 for a given DNA 8-mer.
- **contrastDifference_TBX5-3:** contrastDifference, calculated using the upbm command kmerTestContrast between TBX5-ref and TBX5-3. This is the difference in affinity between TBX5-3 and TBX5-ref for a given DNA 8-mer.
- **contrastQ_TBX5-3**: contrastQ, calculated using the upbm command kmerTestContrast between TBX5-ref and TBX5-3. This is the q-value of the measured difference in affinity between TBX5-3 and TBX5-ref for a given DNA 8-mer.

### Table S10: Additional clones of a single isoform per TF gene used in the paralogs analysis, related to Figure 5.

- **clone_id:** isoform clone ID
- **gene_symbol:** TF gene
- **isoform_status:** one of either: annotated reference, annotated alternative, novel reference, or novel alternative

- **gencode_transcript_names:** GENCODE transcript names for annotated isoforms
- **ensembl_transcript_ids:** Ensembl transcript IDs for annotated isoforms
- **cds_seq**: nucleotide sequence of cloned isoform
- **aa_seq:** amino acid sequence of the isoform
- **tf_family:** TF DBD family

### Table S11. Paralogs results, related to Figure 5.

- **gene_symbol_a:** gene symbol of first tested protein
- **clone_id_a**: clone ID of first tested protein
- **gene_symbol_b:** gene symbol of second tested protein
- **clone_id_b**: clone ID of second tested protein
- **category**: whether the two tested proteins are paralogs, isoforms, or non-paralog/non-isoform controls
- **aa_seq_pct_identity**: percent amino acid sequence identity between the two tested proteins
- **PDI_Jaccard_d**: Jaccard distance between the two proteins' PDI profiles from eY1H assays
- **PPI_Jaccard_d**: Jaccard distance between the two proteins' PPI profiles from Y2H assays
- **activation_abs_fold_change_log2**: absolute log2 fold-change in activation between the two proteins from M1H assays

### Table S12. Condensate and localization results, related to Figure 6.

- **clone_id:** cloned isoform ID
- **cloned_reference_or_alternative:** Referece/Alternative
- **condensates_HEK293T/U2OS_r1/2:** whether condensates were observed and if they were in the nucleus and/or cytoplasm for two replicates of two cell lines
- **localization_HEK293T/U2OS_r1/2:** the localization of the GFP tagged TF isoform in the nucleus and/or cytoplasm for two replicates of two cell lines

### Table S13. Negative regulators and rewirers, related to Figure 7.

- **gene_symbol:** TF gene name
- **Ensembl_gene_ID:** TF Ensembl Gene ID
- **family:** TF family
- **reference_isoform:** ID for reference isoform (currently uses internal ID here e.g. TBX5-1)
- **alternative_isoform:** ID for alternative isoform (currently uses internal ID here e.g. TBX5-2)
- **DBD_pct_lost_in_alt:** percent of DBD lost in the alternative isoform as a fraction of amino acids (100 = full DBD loss)
- **PDI_category:** one of either: PDI loss (all PDIs lost in Y1H in alternative isoform), PDI gain (alternative isoform has PDIs but reference isoform does not), no PDI change

(compared to reference isoform, all baits are the same), PDI rewire (other change in PDIs), or NA (PDIs not successfully tested in both reference and alternative isoform)

- **PPI_category:** one of either: PPI loss (PPIs lost in the Y2H in alternative isoform, subdivided by the type of PPIs lost: all, cofactor, signalling, or dimerization partner), no PPI change (compared to reference isoform, PPIs are the same, subdivided by either all PPIs being retained or cofactor/signalling/dimerization PPIs being retained), PPI rewire (other change in PPIs), or NA (PPIs not successfully tested in both reference and alternative isoform)
- **M1H_activation_category:** one of either: activation loss (if reference isoform is above baseline and alternative isoform has a l2fc ≤ -1), repression loss (if reference isoform is below baseline and alternative isoform has a l2fc ≥ 1), activation GoF (gain of function) (if reference isoform is not above or below baseline but alternative isoform has a l2fc ≥ 1), repression GoF (gain of function) (if reference isoform is not above or below baseline but alternative isoform has a l2fc ≤ -1), similar (alternative isoform has an absolute l2fc of ≤ 1), or NA (activation/repression not successfully tested in both reference and alternative isoform)
- **alt_iso_classification:** one of either: negative regulator, rewirer, similar to ref., likely non-functional, or NA
- **detailed_alt_iso_classification:** same as above, with additional details about why an isoform was called a negative regulator when relevant

### Table S14. Paired TCGA BRCA samples, related to Figure 7.

- List of paired normal/tumor BRCA samples used for all TCGA analyses (one TCGA ID for each)

### Table S15. TF isoform BRCA results, related to Figure 7.

- **gene_symbol:** TF gene name
- **isoform:** ID for isoform (uses internal ID for anything in the TFiso1.0 clone collection and ENST ID for anything not in the clone collection; note that therefore this table includes annotated isoforms missing in TF1.0.)
- **median_tumor_iso_pct:** median isoform expression, as a fraction of total gene expression, across all BRCA tumor samples used (1.0 = 100% expression of gene)
- **median_normal_iso_pct:** median isoform expression, as a fraction of total gene expression, across all BRCA normal samples used (1.0 = 100% expression of gene)
- **median_iso_pct_difference:** median_tumor_iso_pct - median_normal_iso_pct
- **pval:** unadjusted p-value comparing tumor isoform percentages to normal isoform percentages using a paired Wilcoxon test
- **n_samps:** number of samples considered in the p-value test, as any samples where the total gene expression was < 1 TPM were removed
- **padj:** adjusted p-value (Benjamini-Hochberg correction method using an FDR of 0.05)

### Table S16. Primers used for cloning isoforms, related to Figure 2.

- **Plate_name:** Plate name
- **Row:** row position
- **Column:** column position
- **Sequence_name:** ID of the primer
- **Sequence:** nucleotide sequence

### Table S17. Pfam domains considered DNA-binding domains in this study, related to Figure 3.

- **dbd:** domain name
- **pfam:** Pfam accession
- **clan:** Pfam clan accession
